# Global population genomics and pangenome of the clubroot pathogen link clonal expansion and effector diversification to resistance breakdown in canola

**DOI:** 10.64898/2026.10.14.759018

**Authors:** Muhammad Asim Javed, Florent Sylvestre, Sandra Marcela Velasco Cuervo, Soham Mukhopadhyay, Rasha Salih, Anne-Sophie Brochu, Elisa Inés Fantino, Jiaxu Wu, Thies Marten Heick, Arne Schwelm, Yoann Aigu, Leonardo Galindo-Gonzalez, Sheau-Fang Hwang, Pierre Hellin, Marika L.T. Rastas, Nazanin Zamani-Noor, Ghita Cordsen Nielsen, Riinu Kiiker, Louise Aldén, Andrea Ficke, Andres Mäe, Danae Riquelme Toledo, Anh Tung Phan Lam, Shin-ichi Ito, Carlos Antonio dos Santos, Margarida Goréte Ferreira do Carmo, Ulrike Miersch, Godfrey Chongo, Jonny Jacobs, Matthew A. Campbell, Peri Tobias, Jacob Downs, Sera Choi, Brent N. Kaiser, Maxim Prokchorchik, Stephen Strelkov, Edel Pérez-López

**Author notes:** Submitting author: Edel Pérez-López. **Corresponding:** Maxim Prokchorchik, Stephen Strelkov, Edel Pérez-López.

## Abstract

Clubroot disease, caused by the obligate biotrophic protist *Plasmodiophora brassicae*, threatens Brassica crops worldwide. Novel pathogen virulence frequently breaks resistance in agricultural crops, and yet explanations for isolate evolution remain unknown. We combined whole-genome sequencing of 192 isolates from 24 countries with a pangenome analysis of 55 high-quality complete or nearly complete genome assemblies and performed functional effector analyses to investigate potential mechanisms of virulence emergence. Three major phylogenetic clusters were identified, including two highly clonal lineages with contrasting virulence profiles, and a genetically diverse population retaining signatures of recombination. Our findings support a model in which genome-wide purifying selection maintained core genome integrity, while effector genes evolved under less constraint. Despite a compact closed pangenome, we identified chromosome 17 as a major hotspot of single nucleotide and structural variations, transposable elements and expansion of ankyrin-repeat effectors. Within the variable chromosome 17, *PbAnk2* alleles from virulent and avirulent isolates encoded proteins with distinct predicted structures. PbAnk2 interacted with the Arabidopsis co-chaperone DJC31 in yeast two-hybrid and BiFC assays, whereas co-immunoprecipitation did not detect an interaction for the allele found in isolates that break *CRa*-mediated resistance, suggesting that allelic variation may alter PbAnk2’s association with DJC31. These findings reveal the clonal expansion of virulent isolates and connect it to effector diversification that led to resistance breakdown and establish a genomic framework for pathogen surveillance and durable clubroot resistance.

## INTRODUCTION

Plant-pathogen interactions are shaped by various evolutionary factors often driven by heterogeneous plant host landscapes, pathogen genome plasticity, and strong selection imposed by resistance deployment^1,2^. In many pathogens, the frequency of emerging new virulent genotypes, generated by recombination or mutation, increases rapidly under constant evolutionary pressure imposed by the resistant crop deployment, leading to the breakdown of bespoke crop resistance^3,4^. In several major fungal and oomycete plant pathogens, the most serious epiphytotic outbreaks have been recorded when the new clonal pathogen lineages broke the plant resistance in a matter of a few cropping seasons^5–8^. The diversity among successful genotypes driving these shifts extends beyond single nucleotide polymorphisms, with structural changes such as gene presence–absence variations and chromosomal rearrangements providing a major substrate for pathogen adaptation^9,10^. These structural variants frequently accumulate in dynamic genomic regions within the highly variable chromosomes enriched with transposable elements (TEs), where the evolutionary arms race between the pathogen and the plant accelerates the functional diversification of genes involved in host-pathogen interactions^11,12,13^.

Most knowledge about the eukaryotic plant pathogens is based on fungal or oomycete species, whereas deep insights into plant pathogenic protists remain sparse ^14^. *Plasmodiophora brassicae* is an obligate biotroph and unculturable Rhizarian protist that causes clubroot disease in *Brassicaceae* worldwide, and represents an extreme yet understudied example of pathogen genome evolution under intense selection imposed by agriculture^15–17^. Clubroot disease management relies almost exclusively on the deployment of clubroot-resistant (CR) cultivars, creating strong and recurrent selection pressure on pathogen populations^17,18^. Resistance breakdown has been repeatedly documented across production systems, indicating rapid pathogen adaptation despite its obligate lifestyle^18–22^. However, the evolutionary mechanisms underlying these adaptations remain poorly understood, largely due to the lack of geographically broad population-scale genomic resources and the organism’s complex life cycle^23^. The clubroot pathogen predominantly reproduces asexually throughout its life cycle^15,17,24^ , with the frequency and contribution of sexual reproduction to genetic diversity poorly understood^17,24,25^. The genome of *P. brassicae* is compact, with a limited number of recombination events, and as a Rhizarian protist, *P. brassicae* occupies a distinct evolutionary space compared to well-studied fungal and oomycete plant pathogens^17,23,25,26^. Comparative genomics studies of genetically and phenotypically diverse *P. brassicae* isolates are essential to understand the genomic architecture and evolutionary mechanisms underlying pathogen adaptation and host resistance breakdown within *P. brassicae* but also more broadly across other plant pathogens.

Plant pathogens use small secreted proteins, called effectors, to subvert the plant innate immunity and to manipulate the host plant physiology to reroute the development and nutrient flow, ultimately promoting disease development^27–30^. Some plants have evolved resistance genes that can recognize these effectors and activate robust immunity responses^31^. These responses create evolutionary pressure on the pathogen to diversify its effectors, evading immunity recognition while retaining their virulent function^32^. In filamentous plant pathogens, population genomic studies frequently show that effector repertoires are shaped by both purifying selection and localized diversification that is driven by structural variation, TE activity, and effector gene gain and loss^10,13,33,34^. Whether the *P. brassicae* effectorome follows similar evolutionary trajectories, and how genomic context shapes its virulence, has remained largely unknown. One of the factors driving selection is the deployment of resistant crop cultivars. An example is the *Cra* gene mediated canola and cruciferous vegetables clubroot resistance, which provides selection grounds for *P. brassicae* and leads to the breakdown of genetic resistance.

Here, we integrate global population genomics of 192 *P. brassicae* isolates from 24 countries with pangenome analyses of 55 chromosome-level genomes to resolve the evolutionary dynamics of *P. brassicae* virulence development. We identify three major genetic clusters with contrasting virulence profiles, including the canola *CRa* resistance gene-breaking clonal lineage, a predominantly avirulent lineage, and a genetically diverse population showing signatures of recombination with a mixed virulence profile. Through genome-wide analyses, we reveal that effector genes are highly polymorphic and concentrated within structurally dynamic genomic regions, where purifying selection is less effective and genetic variation accumulates. Finally, construction of the first *P. brassicae* pangenome pinpointed chromosome 17 as a hotspot of structural and single-nucleotide variation and identified a candidate ankyrin-repeat effector associated with resistance-breaking isolates. Together with our previous findings, its interaction with a plant co-chaperone suggests that *P. brassicae* ankyrin-repeat effectors may repeatedly target chaperone-mediated processes involved in protein folding, quality control, and stress responses. Altogether, these findings provide a genomic framework for understanding the plant pathogen virulence evolution and the emergence of resistance-breaking populations.

## RESULTS

### Global population structure reveals two major clonal *Plasmodiophora brassicae* lineages

To characterize the global genetic diversity of *P. brassicae*, we performed whole-genome sequencing of 192 isolates from worldwide collections. Using this sequencing data, we called the genomic variants and retained 141 isolates with less than 10% missing genotype variant data and a mean genome coverage greater than 30× for downstream population genomic analyses. The dataset spans six continents, with large representation from Canadian canola fields, and includes isolates from North and South America, Europe, Asia, Oceania, and Africa affecting a wide diversity of Brassica hosts (**Fig. 1a, Supplementary Table 1**).

**Fig. 1.**
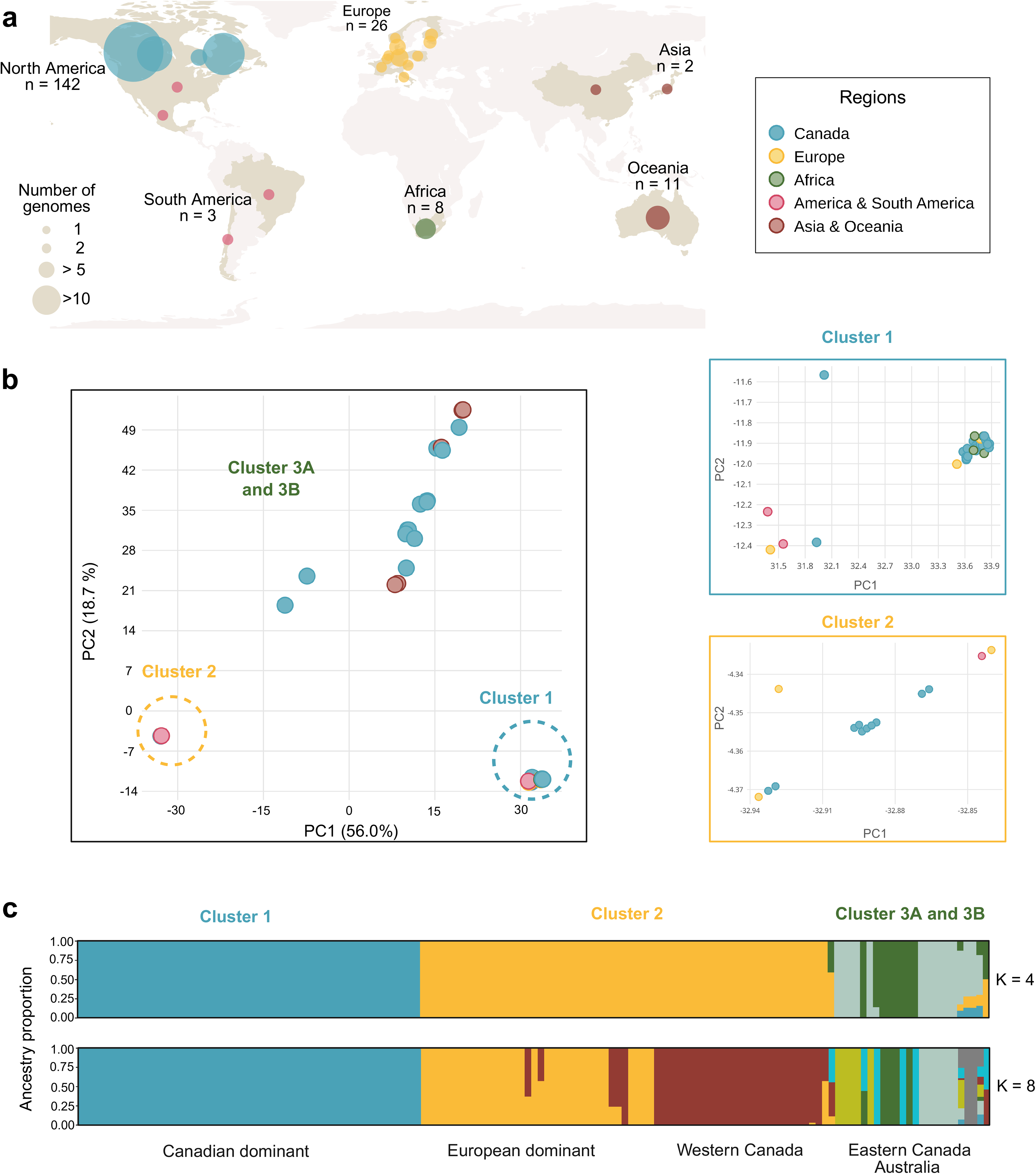
Global sampling and population structure of Plasmodiophora brassicae. **. a**, Geographic distribution of *P. brassicae* isolates across six continental regions. Circle area is proportional to the number of genomes per sampling location. Points are colored by region as indicated in the legend and sample sizes per region are labeled. **b**, Principal component analysis (PCA) of genome-wide SNPs. PC1 and PC2 explain 56.0% and 18.7% of total variance, respectively. Points are colored by geographic region as in a. Three genetic clusters (Clusters 1–3) are indicated and text color is designated to each cluster. Insets show magnified views of the tightly grouped Cluster 1 and Cluster 2, with axis scales reflecting the compressed variance within each cluster. **c**, ADMIXTURE analysis at K = 4 and 8. Each vertical bar represents one isolate where bar segment colors indicate the proportion of ancestry assigned to each of eight inferred ancestral components (numbered 1–8, legend, right). Isolates are ordered and grouped by predominant geographic origin as labeled below the x-axis.

After quality filtering, we obtained 314,193 high confidence biallelic SNPs, of which 3563 statistically independent SNPs were used for population structure analysis. Principal component analysis (PCA) resolved two major genetic clusters, with the first two principal components explaining 74.7% of the total genetic variation. A compact cluster (hereafter Cluster 1) comprised the highly aggressive Canadian pathotypes 5X and 4A, together with recently collected isolates from Alberta and Saskatchewan that predominantly infect commercial CR canola cultivars, defining a western Canadian lineage. The highly virulent European pathotype P1(+), predominantly distributed across central and northern Europe, together with several isolates from northeastern Germany, also grouped within Cluster 1. A second compact cluster (Cluster 2) consisted primarily of European isolates, along with isolates from Alberta and a small number of additional international collections. In contrast, the remaining isolates formed a genetically diverse group with evident two substructures, Cluster 3A and 3B, rather than discrete clusters. These isolates occupied intermediate positions in the PCA and included most eastern Canadian isolates (Québec and Ontario) as well as isolates from Australia (**Fig. 1b**). Running the PCA on isolates sharing less than 90% identity confirmed the separation of Clusters 3A and 3B along the first principal component, which explains 25.1% of the variance (**Supplementary Fig. 1, 2**).

Admixture analysis properly recovered these two Clusters 1 and 2 across all K values, further subdividing Cluster 2 into European and western Canadian isolates with limited admixture between them starting from K=7 (**Supplementary Fig. 4**). Clusters 3A and 3B separated mostly at K=4 and increasing K further substructures these clusters (**Fig. 1c**), reflecting shared ancestry and a more complex population structure. At K=8, the value with the lower cross-validation error (S**upplementary Fig. 3-4)**, Cluster 3A is distributed mainly in three subgroups separating Quebec, Ontario and Australia, with the exception of two isolates that are highly admixed. Similarly, Cluster 3B is divided into two main subclusters, one including isolates from Québec, Alberta and Australia and the other comprising Australian isolates only, with two additional isolates being highly admixed (**Fig. 1c**).

Altogether, these results suggest that Clusters 1 and 2 represent two lineages of *P. brassicae* that are widespread in Canada and most likely expanded through extensive clonal reproduction, potentially from a more diverse population represented by Clusters 3A and 3B, at different times.

### Phylogenomic relationships closely predict virulence profiles

The maximum-likelihood phylogeny built on variant sites across 1231 ortholog genes closely recapitulated the population structure inferred from the PCA and ADMIXTURE analyses, resolving the same three major genetic groups (**Fig. 2a**). Clusters 1 and 2 each formed well-supported monophyletic clades with very short internal branches, consistent with limited genetic diversity among isolates within each lineage. In contrast, Cluster 3A and 3B were genetically diverse and resolved into two respective principal clades. One clade 3B comprised a mixture of isolates from Alberta, Québec, and Australia, with several historical Australian herbarium isolates collected between the 1960s and 1988 occupying basal positions (**Fig. 2a, Supplementary Table 1**). The second clade was composed predominantly of isolates from Québec and Ontario and placed as a sister group of Cluster 1 on the phylogeny. Two Australian isolates grouped within this eastern Canadian clade, forming a distinct subclade (**Fig. 2a**).

**Fig. 2.**
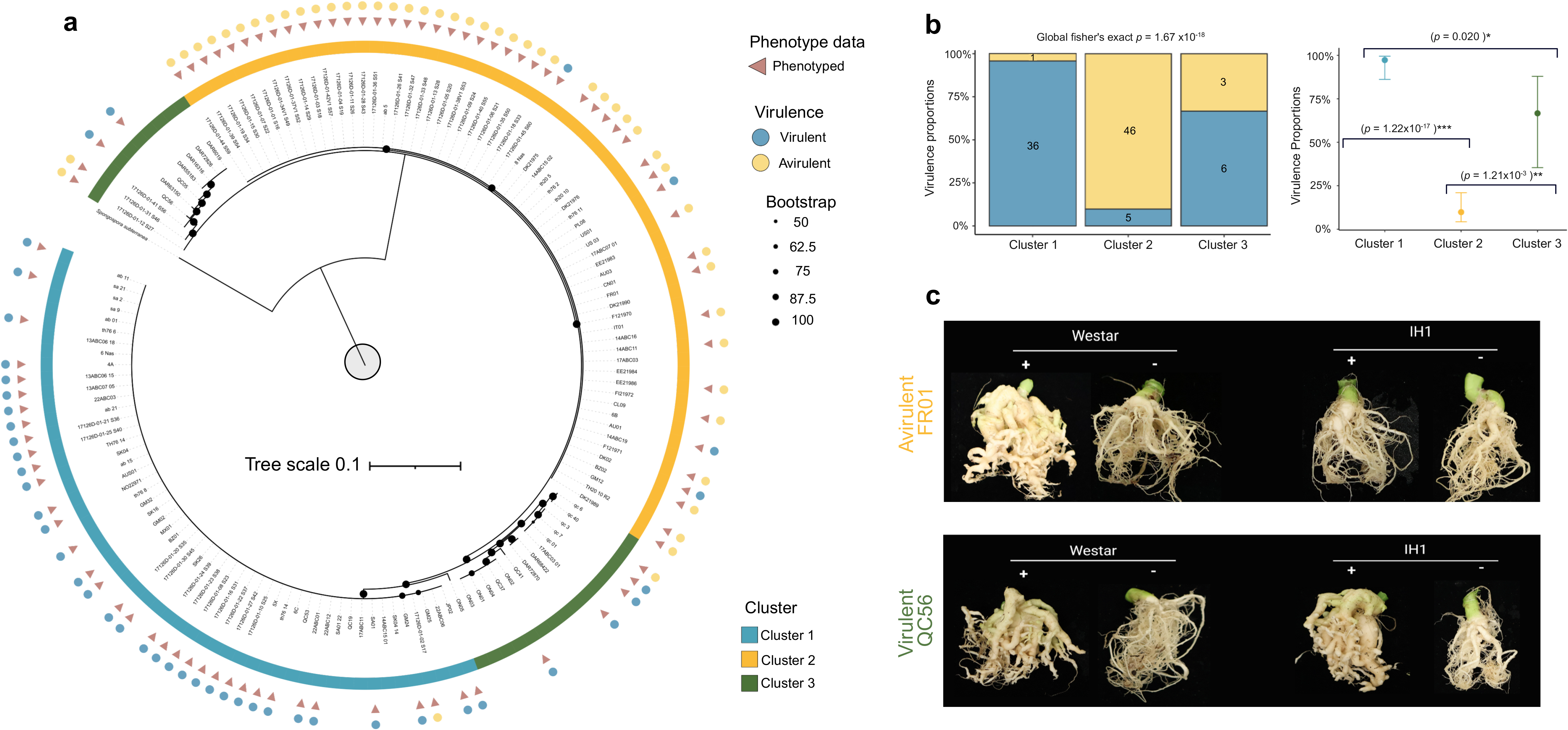
Phylogenetic structure and virulence phenotype distribution across *Plasmodiophora brassicae* population clusters. **a**, Maximum-likelihood (ML) circular phylogeny reconstructed from neutral unlinked single-copy genes across 141 *P. brassicae* isolates. Outer arc coloring denotes cluster assignment (Cluster 1, blue; Cluster 2, orange; Cluster 3A and 3B, green). Node size reflects bootstrap support (1–100). Outer symbols indicate isolates with phenotyping data (burgundy triangles) and virulence classification on resistant host set (filled circles: blue, virulent; yellow, avirulent). Tree scale bar represents 0.1 substitutions per site. **b,** Stacked bar charts (left) showing the proportion of virulent (blue) and avirulent (yellow) isolates within each cluster. Numbers indicate isolate counts per category. Cluster membership and virulence status were significantly associated by Fisher’s exact test (global **p** = 1.67 × 10⁻^18^). Dot plot with 95% Wilson confidence intervals (right) shows pairwise comparisons of virulence proportions between clusters. Significance levels reflect two-sided Fisher’s exact tests with Holm correction (\**p* < 0.05, \*\**p* < 0.01, \*\*\**p* < 0.001). **c,** Representative phenotypes of clubroot symptoms on susceptible (**Brassica napus** cv. Westar) and resistant (IH1 carrying **CRa** resistance gene) hosts. Plants were inoculated with resting spores (+) using isolate FR01 (avirulent on IH1) and QC56 (virulent on IH1), or mock-inoculated (−) with water.

To determine whether phylogenetic clustering was associated with virulence, 97 representative isolates spanning the three major genetic groups were tested on a host carrying the *CRa* resistance gene, either the *Brassica napus* inbred homozygous line IH1 or *Brassica rapa* ECD04 and classified as virulent or avirulent based on their ability to overcome *CRa*-mediated resistance. The susceptible *B. napus* cultivar Westar was used as a control and developed comparable symptoms when infected by virulent and avirulent isolates. Virulence phenotypes differed significantly among the three genetic clusters (global Fisher’s exact test, *p* = 1.67 × 10⁻^18^) (**Fig. 2b**, **Supplementary Table 1**). Cluster 1 contained a significantly higher proportion of virulent isolates than Cluster 2 (*p* = 1.22 × 10⁻^17^) or Cluster 3 (*p* = 0.02) (**Fig. 2b**), including the resistance-breaking pathotypes 5X, 6B, 4A, and P1(+). In contrast, Cluster 2 consisted predominantly of avirulent isolates, including the Canadian pathotypes 3H, 3D, 2A, and 2F. Cluster 3 exhibited a mixed pattern, comprising both virulent and avirulent isolates. Interestingly, the virulent members of Cluster 3 (3A) grouped phylogenetically closer to Cluster 1, whereas the avirulent members (3B) occupied more divergent positions within the lineage (**Fig. 2a**), consistent with multiple evolutionary trajectories towards virulence.

As expected, all isolates induced gall formation on the susceptible cultivar Westar. In contrast, avirulent isolates, such as the French isolate FR01, failed to overcome *CRa*-mediated resistance in IH1 and did not induce gall development (**Fig. 2c**). Highly virulent isolates, such as the eastern Canadian isolate QC56, produced severe galling on the *CRa*-resistant hosts comparable to that observed on the susceptible cultivar Westar, confirming their ability to overcome *CRa*-mediated resistance, consistent with their original field phenotype (**Fig. 2c**).

### Clonal lineages and recombining population exhibit contrasting evolutionary signatures

To investigate the role of clonal and sexual reproduction in the expansion of virulent lineages, we explore linkage disequilibrium (LD) and genetic diversity distribution in *P. brassicae* population. Clusters 1 and 2 showed no detectable LD decay, consistent with predominantly clonal reproduction. In Cluster 1, the fitted LD decay rate (*b*) was not significant (*p* = 0.674), indicating uniformly elevated LD across the 100-kb window (**Fig. 3a, b**). For Cluster 2, only 96 polymorphic SNP pairs were available, precluding reliable model fitting or estimation of LD half-decay distances (**Fig. 3a, b**). This significantly low level of polymorphism is itself consistent with extensive clonality. In contrast, the analysis of Cluster 3A and 3B shows a clear signature of recombination, with an LD half-decay distance of 5.31 kb (non-linear regression fit R^2^ = 0.9338) when grouped (**Fig. 3c**) and 7.76kb (R^2^ = 0.9829) and 34.26kb (R^2^ = 0.8998) for Cluster 3A and 3B respectively (**Supplementary Fig. 5**). Although the low sample size in Cluster 3B likely prevents a precise estimation of LD. Those results are supported by the split-network analysis (**Supplementary Fig. 6**), which showed no evidence of recombination in Cluster 1 and 2 (PHI test, *p* = 1.0) in opposition to the joint analysis of Cluster 3A and 3B, which shows a reticulate topology (PHI test, *p* < 0.001). Together, these analyses support Cluster 3 as a recombining population with mixed virulence profiles, in contrast to the predominantly clonal Clusters 1 and 2. Cluster 1 and 2 also exhibited lower genetic diversity in comparison to combined Cluster 3 (π = 4.112 × 10⁻⁶, 8.867 × 10⁻⁷ and 4.086 × 10⁻³ respectively, all p-value < 0.0001 after Benjamini-Hochberg (BH) correction). Additionally, a low percentage of windows are polymorphic in Cluster 1 and 2 throughout their genome (15.38% and 4.86% of 10 kb windows) where 97.5% of 10kb windows are polymorphic in Cluster 3A and 3B. (**Fig. 3d**, **Supplementary Fig. 7**). These results are concordant with Cluster 1 and 2 being highly clonal lineages. On the opposite, Cluster 3A and 3B genetic diversity seem to be partially shaped by recombination, which could be indicative of past recombination events. However, this result is to be nuanced by their low sample size and the complex substructure observed for these clusters, as suggested by the PERMANOVA results, which show that after controlling for cluster membership (R^2^ = 0.733, *p* < 0.0001), geographic origin (6.2%) and host type (2%) still explain a lower but significant proportion of observed variation (**Fig. 3e**).

**Fig. 3.**
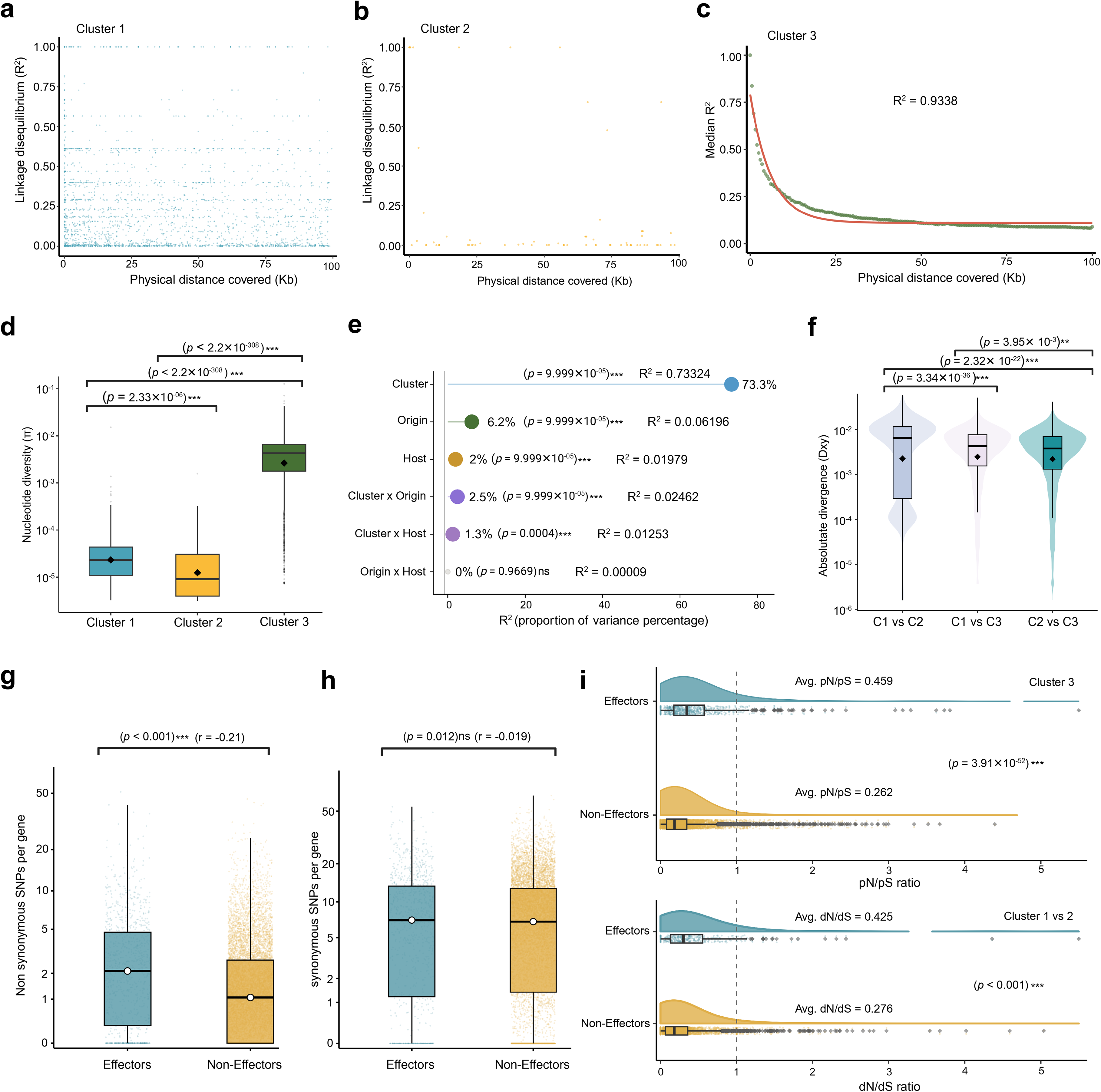
Linkage disequilibrium, genomic diversity, and selection signatures in *P. brassicae*. **a-c,** Linkage disequilibrium (LD) decay patterns revealing distinct reproductive modes across population clusters. Points represent median r² values per 500 bp distance bins; red lines show non-linear least squares (NLS) exponential decay model fits. **(a)** Cluster 1: no systematic decay (R² = 9.9×10⁻⁴) (**b)** Cluster 2: weak non-significant decay (R² = 0.688, ^ns^**p** = 0.089). **(c)** Cluster 3: strong exponential decay (R² = 0.934, \*\*\**p* < 2×10⁻¹, half-decay = 5.31 kb) with significant baseline LD, indicating active sexual recombination. Y-axis: median linkage disequilibrium (r²), linear scale 0-1.0; x-axis: physical distance (kb). **d,** Nucleotide diversity (π) distributions across three population clusters. Bonferroni-corrected pairwise comparisons from post-hoc Dunn tests represent Cluster 1 vs Cluster 3 (\*\*\**p* < 2.33×10⁻¹⁶⁰), Cluster 2 vs Cluster 3 (\*\*\**p* < 2.2×10⁻³⁰⁸), and Cluster 1 vs Cluster 2 (\*\*\**p* < 2.2×10⁻³⁰⁸ demonstrating statistically distinct from each other. Y-axis: nucleotide diversity (log₁₀ scale). **e,** PERMANOVA-based variance partitioning based on factors: cluster, origin, host, cluster × origin, cluster × host and origin × host. Cluster identity as the dominant factor explaining 73.3% of genomic variance (R² = 0.733, **P** < 0.0001). **f,** Absolute divergence (dXY) distributions for pairwise cluster comparisons reveal contrasting genomic divergence patterns. Benjamini–Hochberg corrected pairwise comparisons from post-hoc Dunn tests represent C1vsC2 – C1vsC3 (\*\*\**p* = 3.3×10⁻³⁶), C1vsC2 – C2vsC3 (\*\*\**p* = 2.3×10⁻²²), and C1vsC3 – C2vsC3 (\*\**p* = 3.9×10⁻³), demonstrating significantly different divergence patterns between all cluster pairs. Both b-c Y-axis values are log₁₀ scale. **g-h,** Distribution of synonymous and non-synonymous SNPs per gene across all 141 isolates based on SnpEff variant annotation of 314,193 polymorphic sites. **(g)** Non-synonymous SNPs per gene show significantly elevated counts in effector genes compared to non-effector genes (Mann-Whitney U test, \*\*\**p* < 0.001), **(h)** Synonymous SNPs per gene display non-significant differences between effector and non-effector genes (^ns^*p* = 0.012). **i,** Selection pressure analysis comparing pN/pS ratios (top) and dN/dS ratios (bottom) for effector versus non-effector genes. **Top**: pN/pS ratios in Cluster 3 show significantly elevated values in effectors (mean = 0.459) compared to non-effectors (mean = 0.262) (Mann-Whitney U test, \*\*\***p** = 3.91×10⁻⁵³). **Bottom**: dN/dS ratios between Clusters 1 and 2 similarly demonstrate elevated divergence in effectors (mean = 0.425) versus non-effectors (mean = 0.276, \*\*\***p** < 0.001). Dashed vertical line at ratio = 1.0 indicates neutral evolution threshold and violin plots show kernel density distributions with embedded box plots.

Pairwise absolute divergence (dxy) further highlighted contrasting evolutionary relationships among the clusters. Comparisons between Clusters 1 and 2 produced a strongly bimodal distribution, with 30.6% of genomic windows showing no sequence divergence, consistent with extensive identity by descent. The polymorphic regions of the genome exhibit strong differentiation between Cluster 1 and 2 (mean dxy = 7.094 × 10⁻³). By contrast, comparisons involving Cluster 3 (Clusters 1 vs. 3 and 2 vs. 3) showed a unimodal dxy distribution averaging 5.110 × 10⁻³ and 4.650 × 10⁻³, respectively (**Fig. 3f)**. The strong differentiation between Cluster 1 and 2 appears to be circumscribed to certain genomic regions and could reflect fast differentiation events in response to natural selection (**Supplementary Fig. 8**). To explore the role of natural selection in fast genomic differentiation between clonal lineages and its role in resistance-breaking, we used SNPEff to functionally annotate genetic variants. This revealed a disproportionate accumulation of functional mutations in effector genes. Although putative effectors comprised only 964 of the 10,489 predicted genes (9.2%), they accumulated significantly more non-synonymous mutations per gene than the remaining 9,525 non-effector genes (Mann-Whitney U test, *p* = 1.66 × 10⁻²⁶) (**Fig. 3g,h**). Effector genes contained 74,085 missense variants, 246 nonsense mutations, and 277 splice-site variants, including 517 predicted high-impact mutations, indicating that a substantial proportion of genetic divergence among populations is concentrated within the effector repertoire (**Supplementary Table 3**).

We next compared pN/pS ratios to quantify selective pressure acting on effector and non-effector genes within clusters. Because Clusters 1 and 2 contained very few polymorphic sites, reliable pN/pS estimates could only be obtained for Cluster 3. Effectors exhibited significantly higher pN/pS ratios than non-effectors (0.459 versus 0.262; Mann-Whitney U test, *p* = 3.91 × 10⁻⁵³) (**Fig. 3i**). Nevertheless, both groups showed mean pN/pS values below one, consistent with pervasive purifying selection across the genome. The elevated pN/pS ratios observed in effectors therefore suggest reduced selective constraint, allowing greater accumulation of amino acid-changing mutations relative to the genomic background.

Consistent with this pattern and previous dxy analysis, comparisons between the two clonal lineages (Clusters 1 vs. 2) revealed significantly higher dN/dS values for effectors than for non-effectors (0.425 versus 0.276; Mann-Whitney U test, *p* < 0.001) (**Fig. 3i**). Only a small proportion of genes showed dN/dS values exceeding 1 (3.8% of effectors and 1.2% of non-effectors), indicating that the genome-wide increase in effector dN/dS ratio is primarily consistent with relaxed purifying selection rather than widespread positive selection. In contrast, comparisons involving Cluster 3 showed no significant differences between effectors and non-effectors (**Supplementary Fig. 9**), likely because genetic divergence is distributed uniformly across the recombining ancestral population, obscuring the contrast observed between the derived clonal lineages.

### The *Plasmodiophora brassicae* pangenome reveals a highly conserved genome architecture

To characterize large-scale structural variation across the global *P. brassicae* population, we generated chromosome-level genome assemblies for 53 representative isolates spanning the three major genetic clusters, diverse geographic origins, and contrasting virulence phenotypes. Two previously published long-read assemblies, e3 from Germany^35^ and Race4 from China^12^, were included, resulting in a pangenome comprising 55 genomes. Thirty-three assemblies were completely resolved at the chromosome level, eight contained only one or two fragmented chromosomes, and the remaining assemblies consisted predominantly of chromosome-scale contigs (**Supplementary Table 4**). Assembly sizes ranged from 24.0 to 25.8 Mb, while chromosome number remained highly conserved (**Fig. 4a**). Gene completeness was similarly high, with a mean BUSCO score of 90.4%, although four assemblies displayed lower completeness (**Fig. 4b**, **Supplementary Table 4**).

**Fig. 4.**
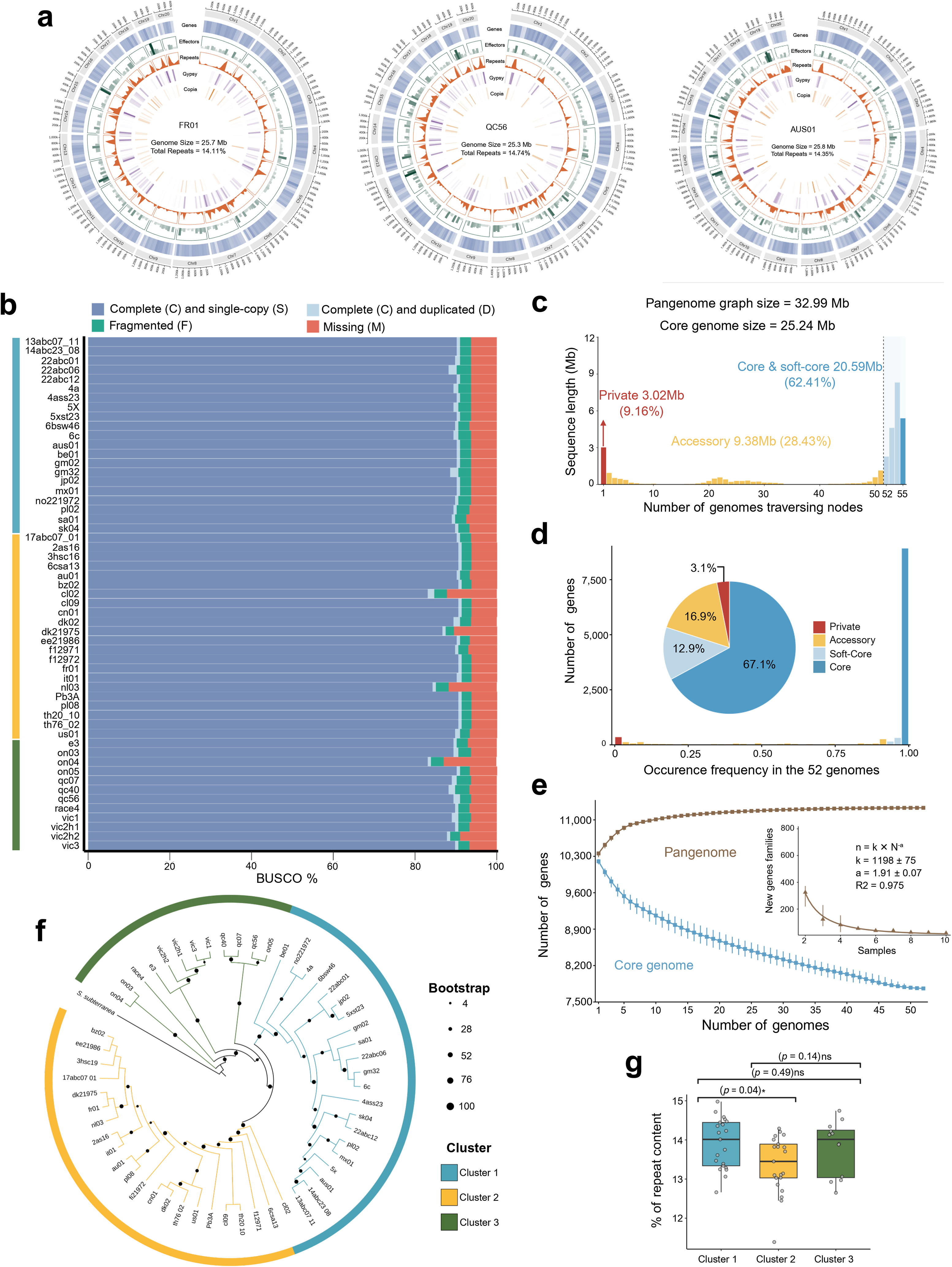
Pangenome and quality statistics of 55 *P. brassicae* complete and nearly complete assemblies. **a**, Circos plots for three representative assemblies (FR01, QC56 and AUS01). Concentric tracks from outermost to innermost display chromosomal coordinates, protein-coding gene density (blue), predicted effector gene positions, total repetitive element content, Gypsy retrotransposon density (orange) and Copia retrotransposon density. **b,** BUSCO completeness scores across all 55 *P. brassicae* assemblies. Stacked bars show the proportion of complete single-copy (dark blue), complete duplicated (light blue), fragmented (green) and missing (red) BUSCO genes. Coloured side bars indicate population cluster membership. **c,** Node-based decomposition of the *P. brassicae* pangenome graph constructed from all 55 assemblies using Minigraph-Cactus. Bars represent total sequence length (Mb) contributed by nodes traversed by a given number of genomes. Annotated compartments correspond to private sequences (3.0 Mb, 9.2%), accessory sequences (9.4 Mb, 28.4%) and core and soft-core sequences (20.6 Mb, 62.41%). **d,** Pangene frequency distribution across 52 assemblies retained after quality filtering. The bar chart shows the number of pangene clusters as a function of occurrence frequency. The pie chart inset summarises the proportional contribution of four pangene categories: core (71%), soft-core (9%), accessory (11%) and private (4%). **e,** Pangenome and core genome accumulation curves across 52 assemblies. Points represent median gene family counts from 1,000 random resampling iterations per sampling depth and error bars show the interquartile range. The pangenome (brown) approached saturation at approximately 25 accessions while the core genome (blue) stabilised at approximately 7,500–8,000 gene families. Inset shows the rate of new gene family discovery per additional genome fitted by a power-law decay model (n = k × N^a; k = 1,198 ± 75, a = 1.91 ± 0.07, R² = 0.975). **f,** ML phylogenetic tree of 55 *P. brassicae* isolates rooted by *S.subterranea*. Branch colors indicate population cluster assignment. **g,** repeat content (%) compared across three population clusters. Centre lines denote medians, box bounds denote interquartile ranges and whiskers extend to 1.5 times the interquartile range. Individual data points are overlaid. Pairwise comparisons were performed using a Kruskal–Wallis test with post hoc correction. Significance levels: \**P* = 0.04; ns, not significant.

The resulting pangenome graph comprised 32.99 Mb, representing approximately a 1.3-fold expansion relative to the 25.24-Mb core genome size. Despite this increase, the graph was dominated by core and soft-core sequences, which together accounted for 20.59 Mb (62.41%) of the pangenome. Accessory sequences contributed 9.38 Mb (28.43%), whereas isolate-specific private sequences represented only 3.02 Mb (9.16%) (**Fig. 4c**, **Supplementary Table 5**).

Gene-based pangenome analysis identified 11,564 orthogroups from an average of 10,371 annotated protein-coding genes per genome. Of these, 7,759 (67.1%) were present in all isolates, 1,489 (12.9%) were classified as soft-core, 1,959 (16.9%) as accessory, and only 357 (3.1%) were private (**Fig. 4d**, **Supplementary Table 5**). Thus, nearly 80% of the gene repertoire was shared among almost all isolates, highlighting the high conservation of gene content despite the global distribution and contrasting virulence phenotypes of the sampled populations.

The pangenome accumulation curve approached saturation after approximately 25 genomes, with additional assemblies contributing very few new gene families (**Fig. 4e**). Fitting Heaps’ law [F(N) = κ × N^γ] to the pangenome accumulation curve yielded an exponent γ = 0.018 (κ = 10,515; R² = 0.928), well below the threshold of γ = 0.5^36^, classifying the pangenome of *P. brassicae* as closed (**Supplementary Table 5**). Further, the rate of new gene family discovery was independently modelled as n = 1,198 ± 75 × N^(−1.91 ± 0.07) (R² = 0.975), the deceleration exponent 1 = 1.91 substantially surpassing the threshold of 1^36^, confirming rapid saturation of gene space by the 10th isolate, with each additional isolate genome contributing less than 15 new gene families (**Fig. 4e**).

Phylogenetic reconstruction using 888 pangenome wide single-copy non-effector orthogroups recovered Cluster 1, 2 and 3A (**Fig. 4f**) but placed Cluster 3A in a basal rather than intermediate position relative to Cluster 1 and 2, confirming the fast expansion of these lineages, which complicates the resolution of their evolutionary history. *De novo* repeat annotation showed that transposable elements (TEs) comprised 11–14% of each genome and were significantly more abundant in the virulent Cluster 1 than in the predominantly avirulent Cluster 2 (*p* = 0.04). LTR retrotransposons, particularly Gypsy and Copia elements, dominated the repeat landscape, whereas DNA transposons, LINEs, rolling circle (RC/Helitron) elements, and unclassified repeats accounted for the remainder (**Supplementary Fig. 10**).

### Structural variation is shaped by population structure and repeat-rich regions

To enable reference-free comparison across high-quality genome assemblies, we constructed a pangenome graph using Minigraph-Cactus^37^. The graph revealed a highly conserved genomic backbone across all 20 chromosomes, interspersed with localized regions of structural variation represented by alternative graph paths. Several large structurally variable regions were detected on chromosomes 10, 11, 16, and 17, many of which displayed lineage-specific patterns, indicating that structural variation is non-randomly distributed across the genome and associated with genetic cluster membership (**Fig. 5a**).

**Fig. 5.**
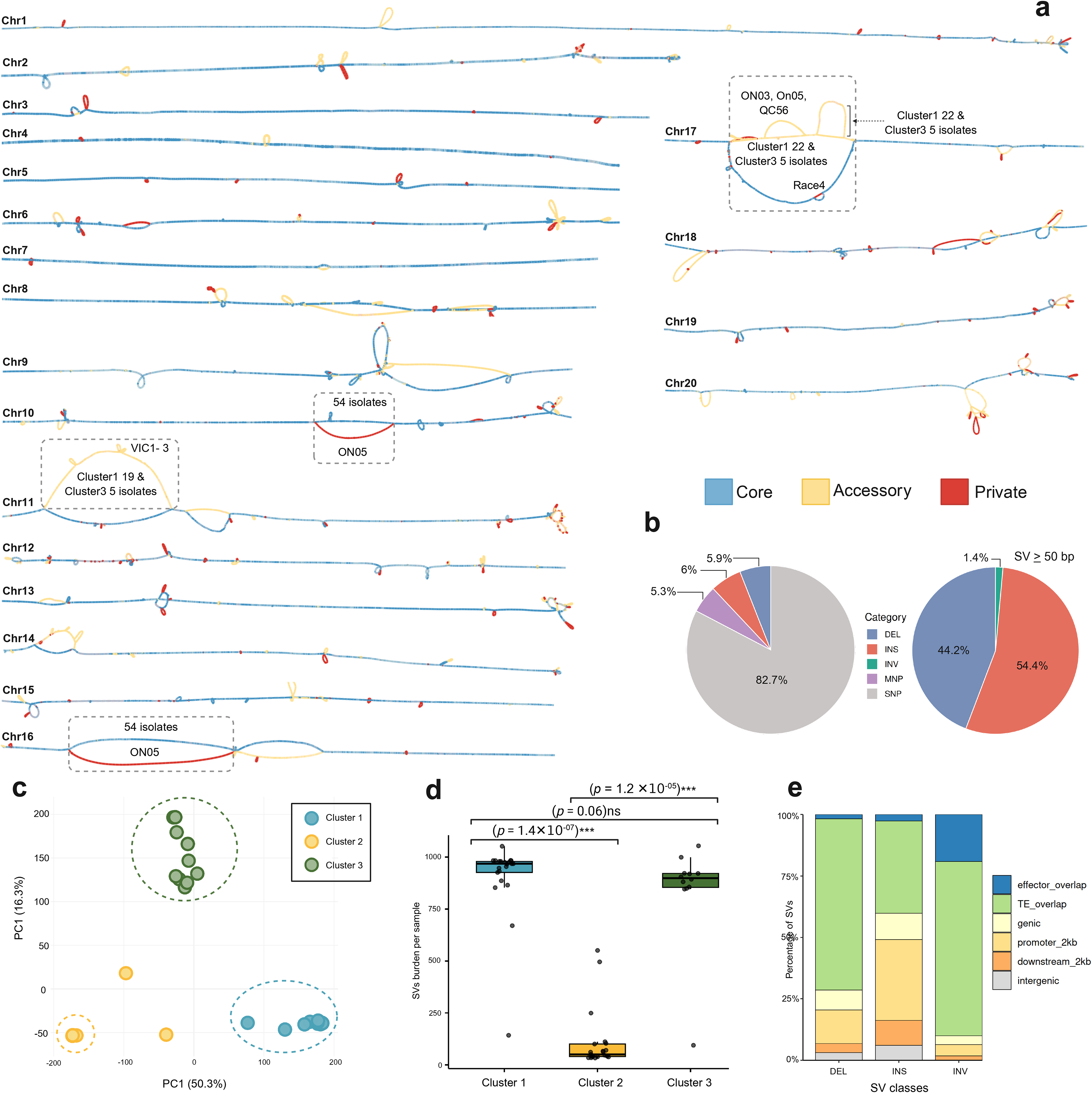
Pangenome graph construction and structural variation landscape. **a,** Graph representation of the *P. brassicae* pangenome across all 20 chromosomes constructed using Minigraph-Cactus. Segments are colored by pangenome category: core (blue, present in all isolates), accessory (yellow, present in a subset of isolates), and private (red, unique to individual isolates). Dashed boxes highlight notable structural variant bubbles on Chr10, Chr11, Chr16, and Chr17, with annotations indicating the isolates from different clusters contributing to each alternate path. **b,** Left, proportional composition of all genomic variants identified across the pangenome, comprising single-nucleotide polymorphisms (SNPs), deletions (DEL), insertions (INS), multi-nucleotide polymorphisms (MNPs), and inversions (INV). Right, proportion of structural variants ≥ 50 bp by class. **c**, PCA of structural variant profiles across **n** = 141 isolates. Each point represents one isolate, colored by population cluster (Cluster 1, blue; Cluster 2, yellow; Cluster 3, green). **d**, SV burden per isolate stratified by clusters. Box plots display median (centre line), interquartile range (box), and 1.5× IQR (whiskers); individual points represent isolates. Pairwise comparisons using the Kruskal–Wallis test with Dunn’s post-hoc correction show significant differences were detected between Cluster 1 and Cluster 2 (\*\*\***p** = 1.4×10⁻⁰⁷) and Cluster 2 × Cluster 3 (\*\*\***p** = 1.2×10⁻⁰⁵), with no significant difference between Cluster 1 and Cluster 3 (**p** = 0.06). **e**, Proportional distribution of DEL, INS, and INV events intersecting annotated genomic features: effectors, TEs, genic, promoter-proximal regions (2 kb upstream), downstream (2 kb), and intergenic regions.

Joint variant calling across 55 genomes yielded 386,555 high-quality variants distributed across all chromosomes. Variant composition was dominated by SNPs, which accounted for 319,091 sites (82.7%), followed by multi-nucleotide polymorphisms (MNPs) with 21,678 sites (5.3%) (**Fig. 5b**). Short insertions and deletions occurred at comparable frequencies, with 23,178 insertions (6.0%) and 22,608 deletions (5.9%). In addition, we identified 7,657 large structural variants (SVs ≥ 50 bp), comprising 4161 insertions (54.4%), 3,386 deletions (44.2%) and 110 inversions (1.4%) (**Fig. 5b, Supplementary Table 6**). In addition, PCA based on large SVs followed the phylogenetic tree and population genetic structure of the three clusters (**Fig. 5c**), described earlier.

Structural variant burden, defined as the number of large insertions or deletions (≥50 bp) carrying a non-reference genotype, was significantly higher in Clusters 1 and 3 than in Cluster 2 (mean = 903.4, 844.9, and 109.9 SVs per isolate, respectively; Kruskal–Wallis test with BH correction, *p* = 1.4 × 10⁻⁷ and 1.2 × 10⁻⁵). On average, genomes from Clusters 1 and 3 harboured approximately eightfold more large structural variants than genomes from Cluster 2 (**Fig. 5d**, **Supplementary Table 7**).

To characterize the genomic distribution of structural variation, we intersected 7,657 SVs (≥50 bp) with gene and repeat annotations. More than half of all SVs overlapped annotated transposable elements (TEs; 3996 total events), indicating that structural variation is concentrated within repeat-rich regions. Promoter regions (defined as 2 kb upstream of genes) represented the second most common category, with 1,835 SVs (24%), suggesting extensive structural remodelling of putative regulatory sequences. Smaller proportions of SVs occurred within genic regions (723 SVs; 9.4%), downstream regions (552 SVs; 7.2%), and intergenic regions (357 SVs; 4.7%). Only 194 SVs (2.5%) overlapped candidate effector genes, indicating that most structural variation is associated with repetitive and regulatory regions rather than effector coding sequences (**Fig. 5e**, **Supplementary Table 7**).

### Population structure shapes the *P. brassicae* pan-effectorome

To determine how effector gene content varies across the *P. brassicae* pangenome and whether effectors are organized in genomic clusters, as documented in other plant pathogens, including *Ustilago maydis* and *Peronospora effusa*^38–40^, we constructed an effector pangenome using 52 isolates representing all three major population structures. A total of 1,160 effector orthogroups were identified. Core and soft-core OGs accounted for 65.34% of the pan-effectorome, including 52.76% that were present in all isolates, indicating that most predicted effectors are highly conserved across the species. A 31.64% of the effectorome was shared across a subset of isolates, whereas only 3.02% were private to individual isolates (**Supplementary Table 8**).

Presence-absence variation (PAV) analysis of the 361 distributed OGs identified 46 lineage-specific effector OGs, including eight unique to the virulent Cluster 1, 10 unique to the avirulent Cluster 2, and 28 unique to Cluster 3, which exhibits a mixed virulence profile. Cluster 3 also contained the largest number of private effector OGs, indicating a substantially expanded accessory effector repertoire in this lineage, composed primarily of eastern Canadian and Australian isolates. Most lineage-specific and private effectors lacked recognizable functional domains, highlighting the large proportion of uncharacterized effector diversity within the species (**Supplementary Table 8**). We next examined six expanded effector families with reported or predicted roles in plant-pathogen interactions, and previously identified by us as the most abundant in gall-forming plasmodiophorids^23^, including ankyrin-repeat (ANK) proteins, chitin-binding proteins, glycosyl hydrolases, leucine-rich repeat (LRR) proteins, kinases, and RXLR proteins (**Fig. 6a, Supplementary Table 8**). Among these, ANK and LRR effectors displayed the greatest accessory variation, whereas the remaining families were largely conserved across isolates.

**Fig. 6.**
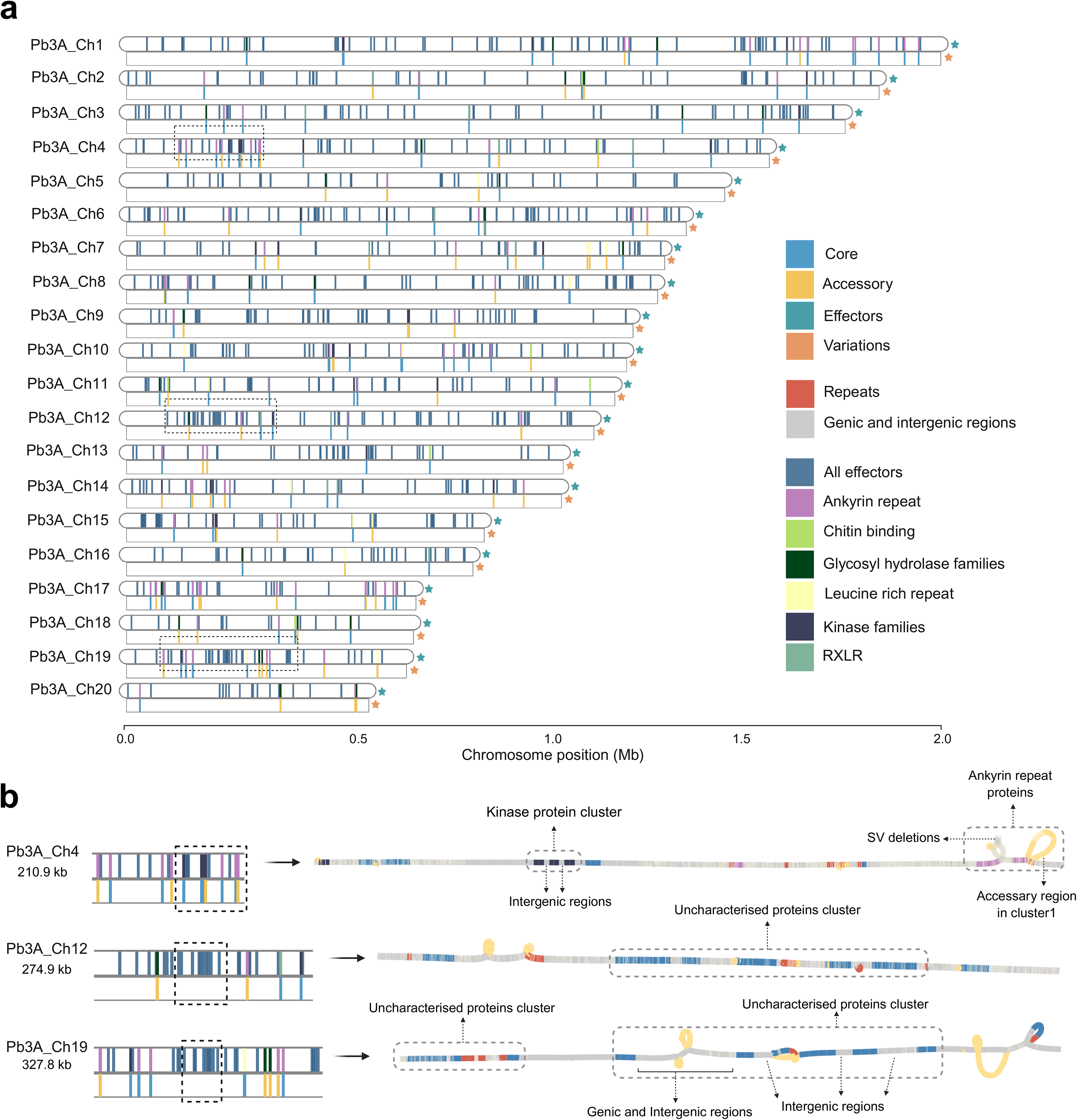
Genome-wide distribution of the pan-effectorome and effector gene clusters. **a**, Chromosomal ideogram of all 20 chromosomes of the *P. brassicae* reference isolate *Pb3A* showing pan-effectorome genomic positions and the presence/absence variations of six effector families. Genes are classified as core (light blue), accessory (yellow), effector-assigned (teal) based on presence-absence variation across the pangenome, with orange marks indicating variations. The positions of all predicted effectors are shown collectively (dark blue), with individual families depicted separately as ankyrin-repeat proteins (pink), chitin-binding proteins (yellow green), glycosyl hydrolase family proteins (dark green), leucine-rich repeat proteins (yellow), kinase family proteins (dark navy), and RXLR effectors (sage green). **b**, Expanded views of the effector-dense cluster windows highlighted in **a** for Chr4 (210.9 kb), Chr12 (274.9 kb), and Chr19 (327.8 kb). Effector clusters are annotated with dashed outlines, including a kinase protein cluster on Chr4 and uncharacterised protein clusters on Chr12 and Chr19. Repeat elements are shown in magenta and genic and intergenic regions in grey.

To determine whether candidate effectors are physically clustered within the genome based on their functions, we mapped the 964 predicted effectors onto the 20 chromosomes of the Pb3A reference genome. Effectors were broadly distributed across all chromosomes, with no chromosome exhibiting a clear enrichment. Nevertheless, small clusters of two to three adjacent effectors, separated by at most a single non-effector gene, were detected on most chromosomes. Prominent examples occurred on chromosomes 4, 12, and 19. The chromosome 4 cluster consisted of closely spaced kinase domain-containing effectors separated by short intergenic regions, whereas clusters on chromosomes 12 and 19 were primarily composed of effectors with unknown functions. Overlaying these annotations onto the pangenome graph revealed that nearly all identified effector clusters reside within core genomic regions, indicating that local effector clustering is evolutionarily conserved despite substantial lineage-specific differences in effector gene content (**Fig. 6a, b**, **Supplementary Table 8**).

### Chromosome 17 is a hotspot of structural variation and ankyrin effector diversification

To determine whether SVs were uniformly distributed across the genome, we compared chromosome length-normalized SV densities across all 20 chromosomes using chi-square goodness-of-fit tests followed by per-chromosome binomial exact tests with BH correction. SVs were distributed non-randomly for all major variant classes, including insertions (INS; χ² = 1,045.68, *p* = 9.34 × 10⁻²¹⁰), deletions (DEL; χ² = 892.82, *p* = 3.81 × 10⁻¹⁷⁷) (**Fig. 7a, b**), and SNPs (χ² = 26,793.22, *p* < 0.005) (**Fig. 7c**). Among all chromosomes, Chr17 (0.747 Mb) exhibited the strongest and most consistent enrichment, with 3.12-fold more insertions, 2.78-fold more deletions, and more than two-fold higher SNP density than expected based on chromosome size. Although inversions were most enriched on Chr14 (χ² = 149.06, *p* = 3.32 × 10⁻²² (**Supplementary Fig. 11**), their relatively low abundance (110 events) warrants cautious interpretation. Regression of fold enrichment against chromosome length further confirmed that the elevated insertion, deletion, and SNP densities on Chr17 cannot be explained by chromosome size alone, suggesting this chromosome as a major hotspot of genomic variation.

**Fig. 7.**
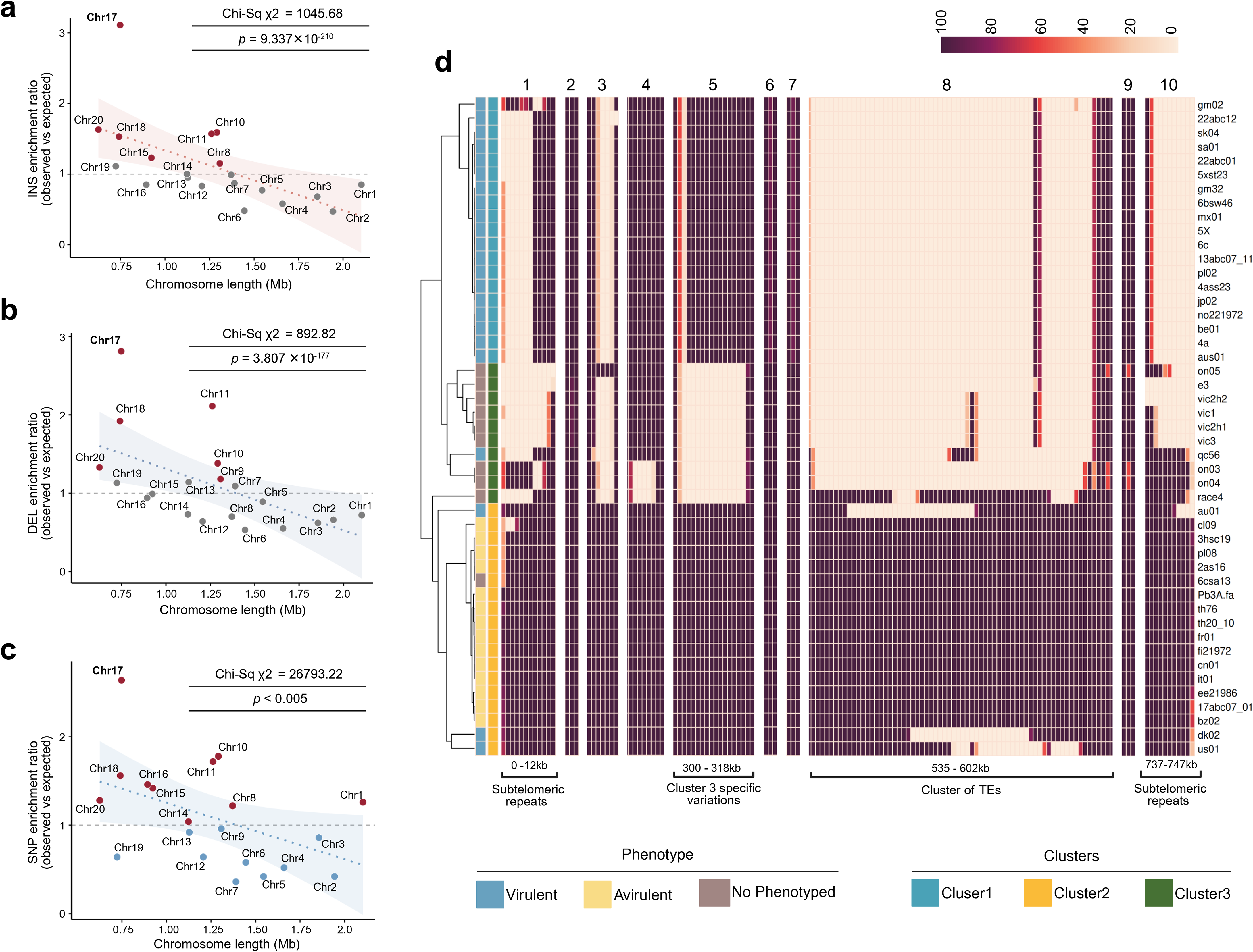
Genome-wide hotspots of structural and sequence variation on Chromosome 17. **a–c,** Scatter plots showing length-normalized density of insertions (INS) **a**, deletions (DEL) **b**, and single nucleotide polymorphisms (SNPs). **c**, per megabase (Mb) plotted against chromosome length (Mb) across all 20 chromosomes. Chi-square goodness-of-fit test (χ²) and associated **P**-values are indicated. Dotted regression lines with 95% confidence intervals (shaded) are shown. **d,** Heatmap of pairwise nucleotide identity (%) across ten low-identity regions on Chr17 for the pangenome dataset. Colour scale indicates per-window alignment identity to the *Pb3A* Chr17 reference (0–100%); dark purple = fully identical cream = diverged (zero values) indicate windows with no detectable alignment, reflecting structural absence or complete sequence divergence in that isolate. Isolates are hierarchically clustered based on Chr17 (left dendrogram) and annotated by phenotype (virulent, light blue; avirulent, yellow; no phenotype, taupe) and population cluster (Cluster 1, teal; Cluster 2, yellow; Cluster 3, dark green). Annotated genomic features include left sub-telomeric repeats (region 1; 0–12 kb), three variable regions (regions 2–4), Cluster 3-specific variation zones (regions 5–6; 300–318 kb), a transposable element cluster (region 8; 535–602 kb) representing the most diverged locus on Chr17, and right sub-telomeric repeats (regions 9–10; 737–747 kb).

Consistent with these observations, Chr17 also exhibited the lowest sequence identity across the genomes. Whereas most chromosomes showed mean pairwise nucleotide identities ranging from 79–92%, Chr17 averaged only ∼70% identity in comparisons between virulent and avirulent isolates (**Supplementary Fig. 12**, **Supplementary Table 9**). This divergence was associated with virulent–avirulent comparisons, while virulent–virulent and avirulent–avirulent comparisons retained nucleotide identities exceeding 80%, comparable to genome-wide levels. Sliding-window analysis across 48 isolates further resolved ten discrete regions of reduced nucleotide identity (**Fig. 7d**). The most divergent interval (535–602 kb) corresponded to a ∼67-kb TE-rich region exhibiting a mean nucleotide identity of only 43.1%. Both sub-telomeric regions were likewise highly divergent, while additional low-identity intervals coincided with Cluster 3-specific variants.

Because Chr17 represented the most structurally divergent chromosome, we next investigated whether it also harbored signatures of effector diversification. Genome annotation identified 198 genes coding for ANK repeat-containing proteins, of those, 45 were located on Chr17 (**Supplementary Fig. 13, 14**). Genome-wide effector prediction further identified 92 putatively secreted ANK repeat proteins, including 12 encoded by the genes located on Chr17 in the *Pb*3A reference genome and five additional accessory ankyrin effectors absent from the reference and avirulent isolates (**Supplementary Table 10**). Notably, nine of the 12 Chr17 ankyrin effectors were expressed during secondary infection at 16 and 26 dpi (**Supplementary Fig. 15**), consistent with previous studies implicating ANK-rich proteins’ importance in gall-forming plasmodiophorids for disease progression.¹⁴⍰²¹

To identify candidate effectors undergoing rapid diversification, we first screened all 964 predicted effectors for variant burden and impact class using SnpEff annotations across the full 141-isolate cohort. High-impact variants (including stop-gained, splice-site, and start-loss mutations) were mostly concentrated within a small subset of effector genes. Among those, two on Chr17 coded for PBTT_09600 (DUF3638 domain-containing protein) and PBTT_09611 (an ANK repeat- and SKP1-containing putative effector, hereafter *PbAnk2*) ranked high among the most polymorphic genome-wide. *PbAnk2* carries 70 non-synonymous mutations (69 missense and one high-impact variant) across its 1,026-bp CDS. To obtain length- and composition-normalised selection statistics, we computed per-site non-synonymous polymorphism rates using the pN/pS pipeline for Cluster 3 isolates and identified the top 15 effectors ranked by pN (**Supplementary Fig. 16**). Within Cluster 3, *PbAnk2* exhibited a non-synonymous polymorphism rate (pN) of 88.09 per 1,000 non-synonymous sites, placing it second among all 964 predicted effectors and well above both the genome-wide mean (8.09 per 1,000 sites) and the effector-class mean (13.17 per 1,000 sites). This elevation was statistically significant relative to the genome-wide background (Fisher’s exact test, FDR = 2.50 × 10⁻⁸). The synonymous polymorphism rate was similarly elevated (pS = 101.18 per 1,000 synonymous sites), yielding a pN/pS ratio of 0.87, consistent with high overall polymorphism at this locus rather than directional positive selection at non-synonymous sites specifically (**Supplementary Table 11**). Additionally, the *PbAnk2* window stood out as a co-localised peak of elevated Cluster3 nucleotide diversity (∼0.02) and elevated inter-cluster divergence (dxy up to ∼0.05 for Cluster1 vs Cluster2), consistent with the locus being a focal point of differentiation among cluster specific isolates (**Supplementary Fig. 17**).

To investigate the genomic context underlying *PbAnk2* diversification, we examined a 241-kb interval (396,622–638,105 bp) encompassing the *ANK* locus using the pangenome graph (**Fig. 8a**). This region contained six ANK repeat effectors, two additional ANK-containing non-effector genes, and a large (∼67-kb) TE-rich region downstream of *PbAnk2* dominated by LTR retrotransposons. Despite overall conservation of gene order, graph traversal revealed pronounced path divergence that distinguished virulent Cluster 1 isolates from avirulent Cluster 2 and representative Cluster 3 genomes, reflecting lineage-specific structural variation concentrated around the *ANKs* locus (**Fig. 8a**).

**Fig. 8.**
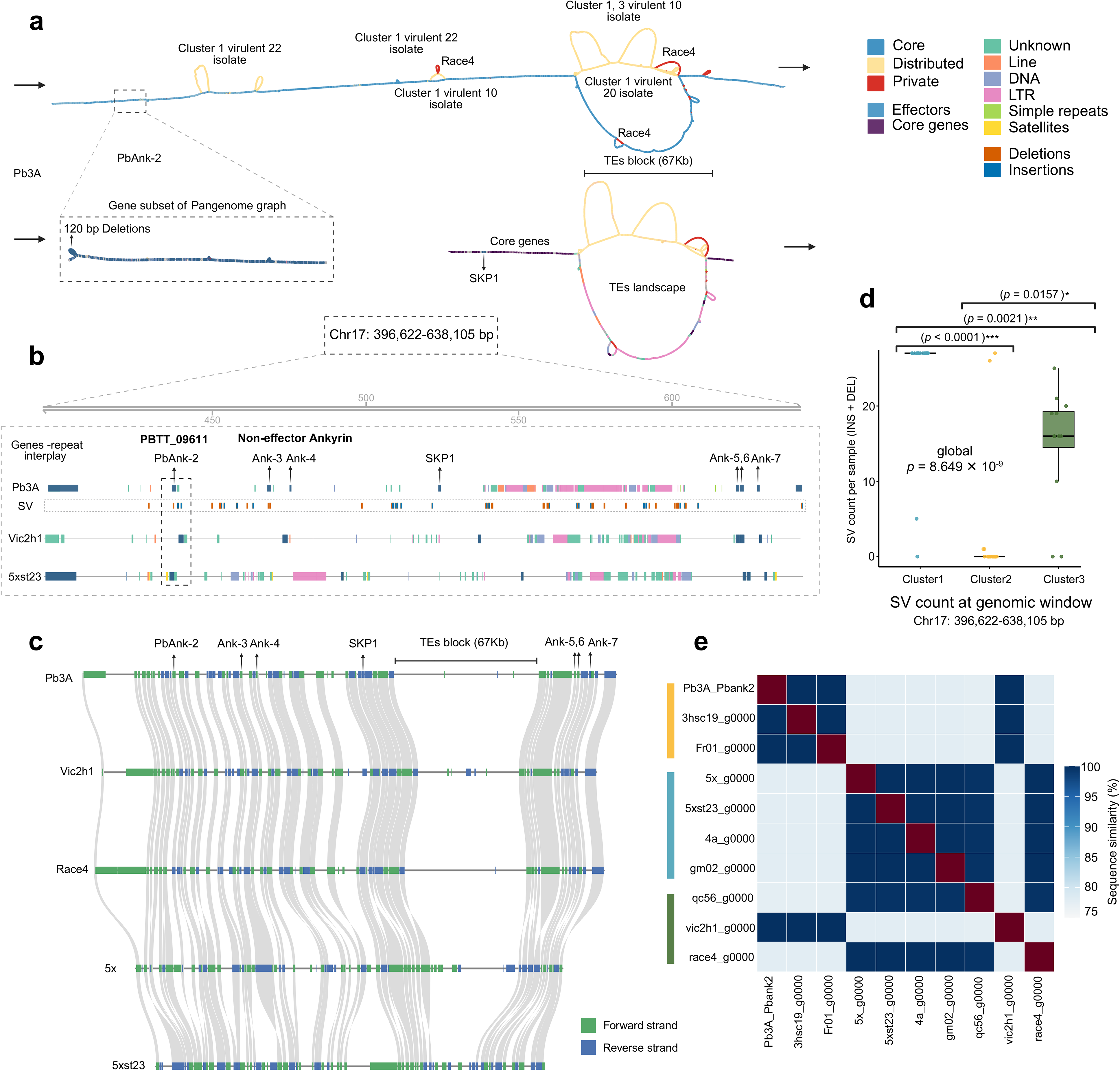
Pangenome variation at *PbAnk2* locus on Chr17. **a**, Pangenome graph traversal across the Chr17 interval (396,622–638,105 bp), showing path topology for Cluster 1 virulent isolates (10- and 22-isolate subsets), Race4, and the Cluster 3 representative. Nodes are coloured by genomic category: core (blue), accessory (orange), private (red), effectors (light blue), core genes (purple), and repeat elements (LINE, DNA, LTR, simple repeats, satellites). Structural variants are annotated as deletions (orange) and insertions (blue). Inset shows the gene-level pangenome graph subset at the *PbAnk2* locus, with a 120-bp deletion visible across traversing paths. The 67-kb transposable element block and flanking *SKP1* core gene are indicated. **b**, Schematic genome representation of the Chr17 interval across three genomes (Pb3A, SV composite track, Vic2h1, 5xst23), showing gene and repeat annotations. *PbAnk2* and flanking non-effector ankyrin-repeat genes (*Ank-3/4, Ank-5/6/7*) and *SKP1* are indicated. The dashed box highlights the structurally variable region immediately surrounding *PbAnk2*. **c**, Synteny alignment across five *P. brassicae* genome assemblies (Pb3A, Vic2h1, Race4, 5x, 5xst23) spanning the Chr17 interval. Coloured blocks indicate forward (green) and reverse (blue) strand alignments. Landmark genes and the 67-kb TE block are annotated above. Grey ribbons connect syntenic blocks between assemblies. **d**, Structural variant (SV) count per isolate including insertions and deletions (≥ 50 bp) within the Chr17 window (396,622–638,105 bp) across the three population clusters (Cluster 1, n = 21; Cluster 2, n = 22; Cluster 3, n = 12). Global difference assessed by Kruskal–Wallis test (χ² = 37.13, *p* = 8.649 × 10⁻⁹). Pairwise post-hoc comparisons by Dunn test with BH correction are indicated above (Cluster 1 vs. Cluster 3: \*\*\**p* < 0.0001; Cluster 2 vs. Cluster 3: \*\**p* = 0.0021; Cluster 1 vs. Cluster 2: \**p* = 0.0157). Boxes show interquartile range with median line; whiskers extend to 1.5× IQR. *e*, Pairwise protein (aa) sequence similarity matrix of *PbAnk2* alleles across nine isolates. Colour scale indicates percent sequence identity (75–100%), with dark red representing one-to-one identity (100%). Isolates are annotated by cluster colour of origin. Two discrete allele types are resolved at approximately 77.78% between-group identity, with within-group identities of 99.75–100%.

Linear comparisons among representative genomes further illustrated extensive TE proliferation surrounding *PbAnk2* in virulent isolates. Numerous simple repeats and satellite sequences flanked the gene, while a large LTR retrotransposon block was inserted approximately 100 kb downstream (**Fig. 8b**). Synteny analysis across representative assemblies showed that coding genes remained largely collinear whereas the internal organization of the TE-rich interval differed substantially among isolates, indicating recent or ongoing TE-mediated structural remodeling (**Fig. 8c**). Consistent with these observations, virulent Cluster 1 isolates exhibited significantly greater insertion and deletion densities across this interval than either Cluster 2 (*p* < 0.0001) or Cluster 3 (*p* = 0.0021) isolates (**Fig. 8d**). Additionally, pairwise protein sequence comparisons identified two major PbAnk2 allele classes largely corresponding to the virulent and avirulent genetic backgrounds. Sequence identity exceeded 95% within allele classes but declined to 75–85% between virulent and avirulent isolates, indicating substantial allelic divergence (**Fig. 8e**).

### Allelic diversification of PbAnk2 remodels its interaction with the host susceptibility factor DJC31

To investigate whether the structural diversification of PbAnk2 translated into functional differences, we first compared the virulent and avirulent alleles. Relative to the avirulent allele, the virulent allele contained multiple single-nucleotide deletions together with a 120-bp deletion, resulting in the loss of 60 amino acids (**Fig. 9a**). This deletion and the surrounding sequence polymorphisms were conserved across all Cluster 1 virulent isolates, whereas Cluster 3 virulent isolates retained >95% amino acid identity with the Cluster 1 allele despite additional nucleotide substitutions. AlphaFold2 structural modelling predicted that the deletion disrupts the C-terminal region spanning the SKP1-binding and ANK-repeat domains, generating an extended disordered region and eliminating one terminal structural fold (**Fig. 9b**). Because PbAnk2 combined three independent lines of evidence supporting signatures of adaptive diversification: (***i***) one of the highest per-site non-synonymous mutation burden among all predicted effectors (**Fig. 8**, **Fig. 9a**), (*ii*) strong expression during the secondary infection stage (**Supplementary Fig. 15**), and (*iii*) localization within the hypervariable Chr17 locus, we selected this effector for functional characterization.

**Fig. 9.**
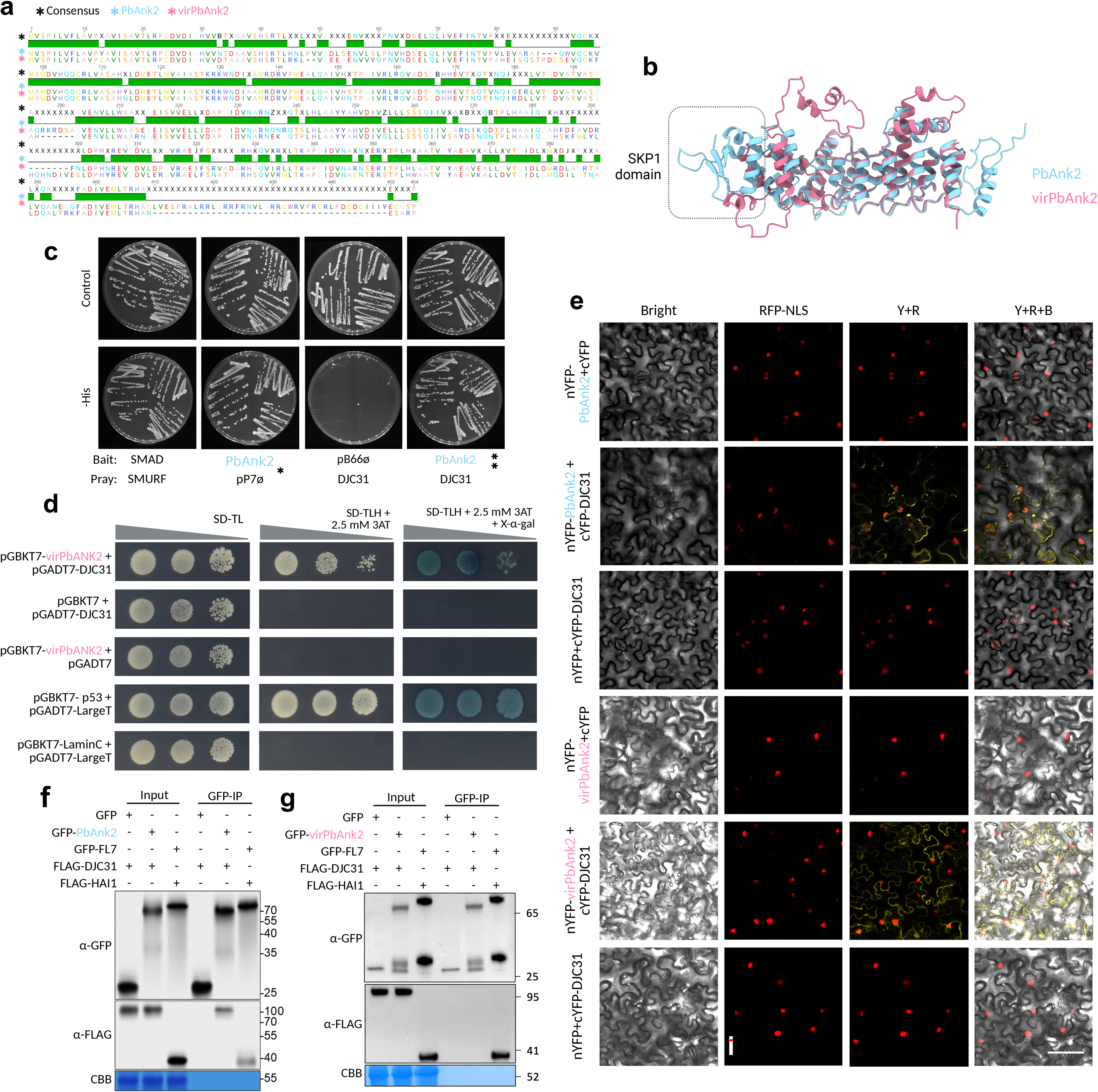
Allele-specific divergence and functional characterization of PbAnk2. **a**, Multiple sequence alignment of two PbAnk2 variants, showing consensus sequence (black), avirulent PbAnk2 variant (light blue) and virulent virPbAnk2 variant (magenta). **b**, Three-dimensional structure of PbAnk2 modelled by AlphaFold, showing the SKP1-binding domain (box) and full-length avirulent PbAnk2 (blue) and virulent virPbAnk2 (magenta) proteins. Structural disorder in the ankyrin repeat domains distinguishes the virulent variant, with the 60 amino acid deletion corresponding to complete loss of one protein fold. **c**, Y2H autoactivation assay for avirulent PbAnk2 as bait against an Arabidopsis seedling library on selective medium without tryptophan, leucine, and histidine, supplemented with 3-aminotriazole (3-AT) to suppress autoactivation. Baits tested include SMAD and SMURF as controls. pB66a and empty prey (pP7a) serve as positive and negative controls, respectively. **d**, Directed Y2H assays testing one-to-one PbAnk2 and virPbAnk2 interactions with DJC31. Top rows show virPbAnk2 interaction (positive growth); lower rows show virPbAnk2 exhibits undetectable interaction under the same stringency than PbAnk2 in **c**. Positive control pairs (p53–LargeT and LaminC–LargeT) confirm assay functionality. *e*, Bimolecular fluorescence complementation (BiFC) assays in *N. benthamiana* leaves co-expressing PbAnk2–nYFP or virPbAnk2–nYFP with DJC31–cYFP. Bright-field and fluorescence channels show RFP-NLS (red), YFP reconstitution (yellow), merged signals (Y+R+B), and overlay. Both avirulent and virulent PbAnk2 produce YFP signal when co-expressed with DJC31, localized to the membrane, whereas negative controls (bottom row) show no fluorescence. **f-g**, Co-immunoprecipitation assays in *N. benthamiana* transient expression system using GFP-tagged avirulent PbAnk2 and virulent virPbAnk2 co-expressed with FLAG-tagged DJC31. Immunoprecipitation specifically pulls down FLAG-DJC31, whereas the virulent variant exhibits undetectable interaction. Controls include GFP and FLAG-DJC31 alone, a positive control pair, GFP-FL7 with FLAG-HAI1, and input lanes confirming comparable protein accumulation.

To identify host proteins targeted by PbAnk2, we screened an *Arabidopsis thaliana* seedling cDNA library using the avirulent allele as bait in a yeast two-hybrid (Y2H) assay. The primary screen recovered 98 His⁺ colonies under stringent selection, from which the tetratricopeptide repeat protein DJC31 (AT5G12430; DNAJ Homolog Subfamily C Member 31) emerged as the strongest candidate interactor (**Supplementary Table 12**). One-to-one Y2H assays confirmed that both the avirulent (PbAnk2) and virulent PbAnk2 (virPbAnk2) variants interact with DJC31 (**Fig. 9c, d**). Mild bait autoactivation was observed for both variants but was completely eliminated by increasing the concentration of 3-amino-1,2,4-triazole (3-AT), confirming that the detected interactions were specific (**Fig. 9c, d**; **Supplementary Fig. 18**).

We next examined whether these interactions occur *in planta* using bimolecular fluorescence complementation (BiFC) in *Nicotiana benthamiana*. Co-expression of either the PbAnk2 or virPbAnk2 variants fused to nYFP with DJC31-cYFP resulted in robust fluorescence reconstitution at the plasma membrane (**Fig. 9e**, **Supplementary Fig. 19**, **Supplementary Fig.20**). No fluorescence was detected in the negative controls, whereas the positive controls produced the expected signal, confirming the specificity of the interaction (**Supplementary Fig. 18**, **Supplementary Fig.19**).

To determine whether both variants formed stable protein complexes *in planta*, we performed co-immunoprecipitation (Co-IP) assays in *N. benthamiana*. GFP-tagged PbAnk2 efficiently co-immunoprecipitated FLAG-tagged DJC31 (**Fig. 9f**), whereas virPbAnk2 failed to recover detectable DJC31 despite comparable accumulation of both proteins in the input fractions (**Fig. 9g**). Positive and negative controls behaved as expected, confirming assay specificity (**Fig. 9f, g**).

## DISCUSSION

Resistance-breaking plant pathogen populations have repeatedly emerged in agricultural systems following deployment of resistant cultivars, yet the evolutionary routes by which many obligate biotrophic plant pathogens adapt have remained unresolved^6,41–43^ . Our global population genomic analyses propose a model where novel virulence does not arise from new independent mutations within clonal populations, but instead from standing variation maintained in a recombining population followed by rapid clonal expansion of successful genotypes. This evolutionary trajectory reconciles decades of conflicting observations describing *P. brassicae* as both highly clonal and genetically diverse and suggests that occasional recombination may provide sufficient evolutionary potential for rapid adaptation under strong host selection.

The genome architecture of *P. brassicae* differs from the canonical two-speed genomes described for many fungal and oomycete pathogens^44–47^. Despite extensive geographic sampling, this species maintains a remarkably closed pangenome with highly conserved gene content. Rather than relying on extensive accessory genome evolution^45,48,49^, adaptive variation appears concentrated within a limited number of localized genomic regions enriched in structural variations and transposable elements. These findings suggest that compact genomes can achieve substantial evolutionary flexibility without undergoing genome-wide compartmentalization.

Our previous work established ankyrin-repeat proteins as the dominant effector family in gall-forming plasmodiophorids^23^. Here we provide evidence that expansion alone does not explain their evolutionary importance. Instead, expansion within dynamic genomic regions created a reservoir of functionally redundant effectors which rapidly diversified through the accumulation of point mutations and structural variations. This likely generated new host specificities while preserving core pathogenic function, favoring the establishment of resistance-breaking lineages.

These findings also challenge the current framework used for clubroot resistance breeding. Pathotype classification has traditionally served as the principal tool for monitoring a broad range of pathogen populations, despite lacking explicit evolutionary information^50–53^. Our data demonstrate that genetically related isolates often share virulence phenotypes across continents, whereas genetically distinct populations may converge on similar pathotypes. This suggests that integrating population genomics with pathotyping could provide a substantially more informative framework for predicting resistance durability, monitoring the emergence of new virulent lineages, and guiding resistance deployment. These findings may be extrapolated to breeding programs aiming to manage other destructive diseases.

Our findings add to evidence that *P. brassicae* infection involves the manipulation of host proteins as well as the evasion of immune recognition. Few host susceptibility factors have been identified in the clubroot pathosystem. These include XCP1, a cysteine protease targeted by a secreted pathogen effector to suppress immunity^54^, and GSL5, whose disruption confers broad-spectrum resistance across Brassica species through jasmonate-mediated defence⁵⁵. Our interaction assays identify the HSP40/DnaJ co-chaperone DJC31 as a candidate host target of PbAnk2, although its contribution to clubroot susceptibility remains to be tested.

Plant chaperone systems are targeted by effectors from other pathogens. The *Pseudomonas syringae* effector HopI1 engages plant HSP70 and suppresses defence^55^; the *Phytophthora sojae* effector PsAvh262 stabilizes the ER chaperone BiP and suppresses stress-triggered cell death^56^; and the *Ralstonia solanacearum* effector RipAC associates with the co-chaperone SGT1 and inhibits SGT1-dependent immunity^57^. In this context, the structural differences between *PbAnk2* alleles and the failure to detect an interaction between DJC31 and the allele from resistance-breaking isolates by co-immunoprecipitation suggest that allelic variation may alter this host association. Whether DJC31 influences clubroot development or participates in CRa-mediated recognition remains to be determined. These observations suggest that adaptation in *P. brassicae* may involve the acquisition or loss of effector functions but also the fine-tuning of host protein interactions through allelic diversification.

## METHODS

### Global collection of clubroot pathogen material

A total of 192 *Plasmodiophora brassicae* isolates were sequenced in this study, collected from 24 countries (**Supplementary Table 1**). Of those, 92 are maintained at the ClubGenoEvo collection at Université Laval (**Supplementary Table 13**). The susceptible host *Brassica napus* L. cv. Westar was used to propagate *P. brassicae* resting spores prior to DNA extraction, following a previously established protocol^58^. For Australian *P. brassicae* isolates maintained at the University of Sydney, Mizuna (*Brassica rapa* var. *niposinica*) plants were used for the propagation of the pathogen (**Supplementary Table 1**).

### DNA extraction and whole-genome sequencing

#### Université Laval isolates

Resting spores were extracted as previously described^58^. High-molecular-weight (HMW) DNA was isolated from suspensions containing 10⁸–10⁹ spores mL⁻¹ using the Nanobind CBB kit (PacBio), following the manufacturer’s recommendations with minor modifications described before^16^. DNA concentration and quality were assessed using a Qubit 4 fluorometer with the dsDNA Broad Range Assay Kit (Thermo Fisher Scientific, Canada).

For short-read sequencing, paired-end (2 × 150 bp) whole-genome shotgun libraries were prepared and sequenced on an Illumina NovaSeq 6000 platform at the Génome Québec sequencing facility. For long-read sequencing, HMW DNA integrity and fragment size distribution were first evaluated using a Femto Pulse automated pulsed-field system (Agilent Technologies, USA) at the IBIS sequencing platform (Université Laval). Genomic DNA was treated with RNase and purified by magnetic bead clean-up, including an FFPE repair step following the manufacturer’s recommendations (Plasmidsaurus Inc., USA). Oxford Nanopore Technologies (ONT) libraries were prepared using the SQK-NBD114.96 Native Barcoding Kit, targeting an approximately 10 kb insert size, and sequenced on PromethION instruments equipped with R10.4.1 flow cells. Raw signal data were base called using Dorado v7.4.12, and reads were demultiplexed with MinKNOW.

#### University of Alberta isolates

Resting spores were isolated and genomic DNA was extracted as previously described^59^. DNA concentration and quality were assessed using a NanoDrop 2000c spectrophotometer and Qubit 2.0 DNA fluorometer with the dsDNA HS Assay (Thermo Scientific). Short-read sequencing was performed as previously described^59^. Libraries were prepared using the KAPA HyperPrep Kit and sequenced on an Illumina HiSeq X platform to generate paired-end (2 × 150 bp) reads. For long-read sequencing, libraries were prepared using the SMRTbell Prep Kit 3.0 and sequenced using PacBio HiFi sequencing on a Sequel IIe system, following the manufacturer’s recommendations.

#### University of Sydney isolates

*P. brassicae* resting spores were purified from field-derived and herbarium isolates using a previously described method with minor modifications^58^. In brief, clean resting spores were centrifuged and concentrated in Milli-Q water. Spores were ground in a chilled mortar with acid-washed sand in liquid nitrogen. Ground spores were washed with 0.35M D-Sorbitol buffer multiple times until the suspension became non-viscous. After that, CTAB gDNA extraction, coupled with Proteinase K and RNase A treatment, was performed^60^. To maximize the gDNA extraction yield, overnight precipitation with 100% Ethanol at -20°C was performed.

High-fidelity (HiFi) sequencing was performed at the Australian Genome Research Facility (AGRF). DNA libraries were prepared using the SMRTbell library preparation kit and sequenced on a PacBio Revio platform using SMRT Cell 25M flow cells. For chromosome-scale scaffolding, Hi-C libraries were generated following chromatin crosslinking and DNA extraction using the Phase Genomics Proximo Hi-C Fungal Kit according to the manufacturer’s instructions. Hi-C libraries were sequenced on an Illumina MiSeq platform at the Biomolecular Research Facility, Australian National University (ANU). Illumina DNA sequencing was performed at the AGRF. Genomic DNA libraries were prepared using IDT xGen cfDNA & FFPE Library Preparation Kit and sequenced with the Illumina NovaSeq X Plus platform. To address the potential low quality of DNA extraction from herbarium isolates, all herbarium samples were first sequenced with low coverage, and the clubroot read quality and fraction were estimated. Low-coverage samples (<70x) were sequenced twice to achieve high coverage among the two runs combined.

#### Variant calling and filtering

Whole-genome short-read data from 192 isolates were quality-checked using FastQC (v0.11.8)^61^. Adapter sequences and low-quality bases were trimmed with Trimmomatic (v0.39)^62^, using the following parameters: LEADING:3; TRAILING:3; SLIDINGWINDOW:4:20; MINLEN:50. Filtered reads were aligned to the *P. brassicae* reference genome Pb3A^16^ using BWA-MEM (v0.7.19)^63^. Alignments were sorted and PCR duplicates marked with Sambamba (v1.0.1)^64^, and BAM files were indexed using SAMtools (v1.22)^65^.

Variants were called with BCFtools (v2.1)^66^ and filtered using a combination of VCFtools (v0.1.17)^67^ and custom Python scripts. To ensure high-confidence variant calls, filtering was applied at both the genotype and site levels. Genotypes were set to missing when allelic balance (ratio of reference allele reads to total reads) ranged between 10% and 90%, indicating ambiguous allele assignment. Genotypes with coverage below 15% or above 300% of the individual’s median sequencing depth were also treated as missing, and sites with a genotype quality (GQ) score <50 were removed. A minor allele count (MAC) threshold was also applied during variant filtering, retaining only sites with MAC ≥ 1. Isolates exhibiting more than 10% missing data across all SNPs were excluded from downstream analyses, and SNP sites with more than 10% missing data across retained isolates were similarly removed. Variant effects were annotated using SnpEff (v5.2c)^68^. The complete workflow, including quality control, variant calling, filtering, and annotation steps, is available at https://github.com/Edelab/Population-genomics-and-Pangenome-analysis.

### Population structure analysis

For population structure analysis, we extracted independent SNPs using LD pruning PLINK (v1.9)^69,70^ (--indep-pairwise 50 20 0.2) and estimated sample pairwise identity using --distance ibs square. Biallelic independent SNPs were converted from VCF to PLINK binary format. We then performed a Principal component analysis (PCA) on a dataset with all samples (n=141) and on a dataset excluding samples with more than 90% identity (n = 15). PCA was computed on the genotype variance–covariance matrix using prcomp in R (base stats)^71^, with the proportion of variance explained by each component visualized with a scree plot. PCA scatterplots were generated using ggplot2^72^.

We then ran ADMIXTURE (v1.3.0)⁶¹ on the unlinked-SNP dataset including all samples, with the haploid genotype model (--haploid="*"). ADMIXTURE was run for K = 1–15 with 10 replicates per K each initialized from an independent time-based random seed. The optimal number of clusters (K) was selected based on the minimum cross-validation error, following the ADMIXTURE manual. Convergence across independent runs and alignment of clusters across K were assessed with Clumppling^73^. For the >90% identity-by-state dataset, cross-validation error decreased monotonically with K and was therefore uninformative, so K was instead chosen from the mode-stability of aligned replicates in Clumppling. Admixture proportions were visualized in R using ggplot2^72^.

To test whether genetic structure was associated with population assignment, geography, or host plant, we performed a permutational multivariate analysis of variance (PERMANOVA) using the adonis2 function in R (vegan package)^74^. The analysis was conducted on a matrix of unlinked SNPs, with genetic distance as the response variable. Geography was defined at the continental scale, except for Canada, where isolates were stratified by province (Québec, Ontario, Alberta, and Saskatchewan). Host plants were classified as vegetable crops or canola. The full model was specified as G ∼ CLUSTER × GEOGRAPHY × HOST, where G represents the genetic distance matrix. Significance was assessed using 10,000 permutations with marginal tests for each term (by = "term").

### Phylogenetic inference and network analysis

To infer evolutionary relationships, phylogenetic analysis was performed by rooting with the closely related plasmodiophorid *Spongospora subterranea* as an outgroup. Single copy orthologs from *P. brassicae* Pb3A reference proteome and *S. subterranea* were identified using OrthoFinder (v2)^75^. For each of the 141 isolates, per-gene coding sequences were reconstructed by applying the isolates called variants to the corresponding Pb3A reference CDS. Protein sequences from every orthologous genes from *S. subterranea* were aligned to the corresponding *P. brassicae* coding sequences using MAFFT (v7)^76^, before being translated back to nucleotide sequences. Alignments missing a start codon or presenting an internal stop codon were removed from analysis, as well as alignments with more than 100 gaps. Genes presenting values of Pn/Ps or dn/ds higher than the 95 percentiles in at least one cluster were also discarded to avoid selection-driven artifacts. A rooted ML phylogeny was then inferred using IQ-TREE (v2)^77^, restricting the analysis to only polymorphic sites in *P. brassicae*. The best-fitting substitution model was selected with ModelFinder Plus under ascertainment-bias correction (-m MFP+ASC), and branch support was assessed with 1,000 ultrafast bootstrap replicates. Phylogenetic trees from both analyses were visualized and annotated using the Interactive Tree of Life (iTOL) (v6)^78^. To visualize conflicting phylogenetic signal not captured by a bifurcating tree, we computed a NeighborNet network from the same codon-based alignment generated using SplitsTree (v6.00)^79^. To test for recombination, we applied the Pairwise Homoplasy Index (PHI) test^80^ to each cluster separately.

### Virulence profile of *P. brassicae isolates*

#### Université Laval isolates

A subset of *P. brassicae* isolates was selected based on genetic cluster and geographic origin to assess virulence phenotypes (**Supplementary Table 1**). Virulence was evaluated using a homozygous inbred canola line carrying the *CRa* resistance gene, kindly provided by Nutrien Ag Solutions, together with the susceptible cultivar Westar. Resting spore suspensions were adjusted to 10⁸ spores mL⁻¹, and seedlings were inoculated following the protocol described previously. Plants were maintained under controlled greenhouse conditions in a randomized complete block design with three biological replicates per treatment, while mock-inoculated plants (water only) served as negative controls. The experiment was independently repeated three times. Disease severity was assessed 28 days post-inoculation (dpi). Throughout this study, virulence refers specifically to the ability of an isolate to overcome *CRa*-mediated resistance. Regardless of virulence status, all isolates induced gall formation and completed their life cycle on the susceptible cultivar Westar.

#### University of Alberta isolates

The 45 isolates processed at the University of Alberta were previously pathotyped^81^ using the 13 host genotypes of the Canadian Clubroot Differential (CCD) set^52^. Resting spore suspensions were adjusted to 10^7^ spores mL⁻¹, and inoculated seedlings were grown for 6 weeks in a greenhouse maintained at approximately 24°C during the day and 18°C at night, under natural light supplemented with artificial lighting to provide a 16-h photoperiod. Each host genotype-isolate combination was represented by 12 plants per replicate and maintained in a split-plot design, with isolates as the main plots and differential hosts as the subplots. One single-spore isolate representing each pathotype was selected and retested on the CCD set in three additional independent experiments. The index of disease (ID) and 95% confidence intervals (CIs) were used to determine host resistance or susceptibility to each isolate, using the thresholds previously described^81,82^. Isolates were classified as virulent when they were able to overcome resistance in *Cra*-bearing genotype.

### Genetic diversity and Linkage disequilibrium

Population level nucleotide diversity (π) and absolute divergence (dxy) were calculated using pixy (v1.2.7)^83^, which provides populations genetics summary statistics. Analyses were performed on the filtered VCF of 314,193 high-quality biallelic SNPs and invariant sites across 141 isolates, which was partitioned into cluster specific VCFs prior to analyses. For within population diversity estimation, π was calculated separately for each cluster using 10 kb non-overlapping sliding windows across all 20 chromosomes. Population assignments were based on previous PCA based clustering analysis, with Cluster 1 (n=54), Cluster 2 (n=65), and Cluster 3 (n=28) analyzed independently. Between population absolute divergence, dxy was computed for all three pairwise cluster combinations (C1 vs C2, C1 vs C3, C2 vs C3) using identical 10 kb windows to enable direct genetic variation comparison between population. Pixy was run with the following parameters: --sites for invariant site handling, --window_size 10000 bp, and --populations specifying cluster membership files. Statistical comparisons between clusters employed Kruskal-Wallis tests followed by Dunn’s post-hoc tests with Bonferroni correction for multiple comparisons.

Linkage disequilibrium (LD) decay was analyzed separately for each population cluster to assess reproductive mode differences. Pairwise LD (r²) was computed using PLINK (v1.90b5.3) in haploid mode. VCF files were converted to PLINK binary format using plink --vcf cluster.vcf.gz --allow-extra-chr--chr-set -20 --double-id --make-bed --out cluster_haploid. Pairwise LD computation was performed using plink --bfile cluster_haploid --allow-extra-chr --chr-set -20 --r2 dprime --ld-window-kb 100 --ld-window 999999 --ld-window-r2 0 --out cluster_ld.

LD decay curves were constructed by binning pairwise r² values by physical distance (500 bp bins) and calculating median r² per bin to provide robust central tendency estimates for right skewed r² distributions. Exponential decay models were fitted to the binned medians by non-linear least squares (NLS) model^6,84^. For each cluster both a two-parameter model (r² = a × exp(-b × distance)) and a three-parameter model accounting for a non-zero baseline LD (r² = a × exp(-b × distance) + c) were fitted, and the model with the lower Akaike Information Criterion (AIC) was retained. The half-decay distance, defined as the distance at which the predicted r² declines to half of its maximum (initial) value, was reported only where the decay rate b was significantly greater than zero (*p* < 0.05) and sufficient SNP pairs were available. For the clonal lineages (Clusters 1 and 2), which showed non-significant or non-estimable decay, no half-decay distance was reported. All curve fitting and statistical analyses were performed in R.

### Assessing the signature of natural selection

To quantify signatures of natural selection, we estimated the ratio of non-synonymous to synonymous polymorphism (Pn/Ps) within each of the three genetic clusters and divergence (dn/ds) between clusters. Individual coding sequences were generated and filtered using custom perl scripts, removing terminal stop codons and codons represented in fewer than four individuals. Within each cluster, the numbers of synonymous and non-synonymous sites were estimated using Bio++, assuming a transition-to-transversion ratio of 2. We used a custom python script to estimate the number of synonymous (Ps), and non-synonymous (Pn) polymorphic sites. Pairwise synonymous (ds) and non-synonymous fixed divergence (ds) were then counted using custom Python scripts. The Pn/Ps ratios were calculated as:

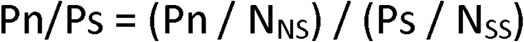

Where Pn and Ps represent the numbers of non-synonymous and synonymous polymorphisms, respectively, and N_NS_ and N_SS_ represent the numbers of non-synonymous and synonymous sites. The same formula was used for estimating dn/ds between pair of clusters, using the pairwise average N_NS_ and N value for normalisation. Scripts were adapted from the PiNPiS pipeline^85^; https://github.com/QuentinRougemont/PiNPiS), and the final versions are available at https://github.com/Edelab/Population-genomics-and-Pangenome-analysis.

### Quality control and genome assembly

ONT read quality was assessed using NanoQC (v1)^86^, and low-quality Nanopore reads shorter than 500 bp were removed using chopper (v0.6.0)^87^. Genome assemblies were generated with hifiasm (v0.25.0)^88^ in Nanopore mode using the inbred (haploid) flag (–l0). For assemblies exhibiting fragmented chromosomes, genomes were independently assembled using Canu (v2)^89^, and complete chromosomes were subsequently replaced into the hifiasm assemblies following the strategy developed for the Pb3A reference genome^16^.

To remove host (*B. napus*) and endophytic microbial contamination, assembled contigs were screened using the Foreign Contamination Screen (FCS) pipeline^90^ and further queried by BLASTN against the NCBI non-redundant nucleotide database using default parameters^15^. BLASTN hits were filtered based on ≥70% query coverage, an E-value threshold of 1 × 10⁻⁵, and GC content consistent with *P. brassicae* (57 to 61%), as previously described^16,38^.

Final genome assemblies were polished using Hapo-G (v1.2)^91^ with Illumina paired-end reads over three iterations. Assembly quality was assessed using QUAST (v2.5.0)^92^, and gene completeness was evaluated with compleasm (v0.2.5)^93^ using the eukaryota_odb10 dataset. Whole-genome alignments for dot-plot visualization were generated using D-Genies (v1.5.0)^94^.

#### University of Sydney isolates

To filter out the non-clubroot reads, raw PacBio reads were aligned to the custom database containing all available *Brassicaceae* plants and NCBI bacterial RefSeq genomes using MiniMap2 (v2.1)^95^ with the ‘-t 24 -ax asm20 --secondary=no’ settings. Aligned reads were discarded and the unaligned read sets of all, longer than 20kb, 15 kb, and 10kb PacBio HiFi reads were assembled using hifiasm (v0.19.8)^88^, with ‘-l 0’ or ‘-l 3 -hic’ flags where the HiC data was available. Each assembly was manually curated to pick the best combination of T2T chromosomes as a whole-genome representation. Chromosomes were ordered and oriented according to the Pb3A genome assembly^16^.

### Genome and transposable element annotations

Genome annotation was performed using an approach optimized for the Pb3A reference genome, as previously described^94^. Protein-coding genes were annotated using BRAKER3 (v3.0.7)^96^, which integrates transcriptomic and protein evidence to automatically annotate repeat-masked genomes. The resulting GTF files were processed using a Python script provided by the BRAKER3 developers (https://github.com/Gaius-Augustus) to retain only the longest isoform for each gene model. *De novo* repeat and transposable elements annotations were carried out using the EarlGrey pipeline (v6.0)^97^, by constructing a custom repeat library specific to *P. brassicae*.

### Pan-secretome and effector predictions

Predicted proteomes from the 55 *P. brassicae* isolates included in the pangenome analysis (**Supplementary Table 4**) were processed using an effector prediction pipeline previously applied to the reference proteome^23,98^. Signal peptides were predicted using SignalP (v6)^99^ and removed prior to downstream analyses. Transmembrane domains were identified using DeepTMHMM via the BioLib wrapper in GPU mode^100^. Candidate secreted proteins were defined as those containing an N-terminal signal peptide and lacking transmembrane helices beyond the signal peptide, following established criteria^23^. Candidate secreted protein sets from all isolates collectively constituted the pan-secretome, which was subsequently used for downstream effectorome analyses.

### Pan-gene and pan-effectorome presence/absence variation analysis

Predicted protein-coding genes from the 55 *P. brassicae* isolates included in the pangenome analysis (**Supplementary Table 4**) were clustered into orthogroups (OGs) using OrthoFinder (v2)^75^ with default settings, generating a pan-genome orthogroup catalogue. For each analysis, a PAV binary matrix was constructed at the orthogroup level across 52 isolates (3 excluded having gene completeness below 80%). Orthogroups present in all 52 isolates were classified as core (including both single-copy and paralogous families), whereas those present in ≥ 95% of isolates were classified as soft-core. Orthogroups present in 2–94% of isolates were designated accessory , and those detected in a single isolate were considered private^101^.

Single-copy orthogroups (SCOs) conserved across isolates were extracted, aligned using MAFFT (v7)^76^, and concatenated into a partitioned supermatrix for phylogenetic inference. An ML phylogeny was inferred using IQ-TREE (v2)^77^ with ModelFinder (–m MFP) and 1,000 bootstrap replicates to assess node support. For pan-effectorome analyses, all effector-associated orthogroups were extracted using custom Python script, and a corresponding PAV binary matrix was generated.

### Pangenome graphs construction and structural variant analysis

A pangenome graph was constructed using the Minigraph–Cactus pipeline, enabling chromosome splitting to generate per-chromosome graph files (GFA)^37^. Post-pangenome filtering and annotation of small and large structural variants were performed following the Human Pangenome Reference Consortium (HPRC) graph pipeline, with all variants harmonized to the reference genome Pb3A coordinate system^102^. Chromosomal graphs were visualized using Bandage-NG^103^ to identify large graph bubbles representing complex structural variation. Variant coordinates were then intersected with genome annotations using BEDTools (v2.31.1)^104^ and we used a custom Python script to classify structural variants according to genomic context, including genic and intergenic regions, transposable elements, exons, introns, and upstream or downstream regulatory regions.

Structural variants were filtered to retain events ≥ 50 bp mapping to the 20 nuclear chromosomes (INS: n = 4,161; DEL: n = 3,386; INV: n = 110; total n = 7,657). To assess whether SVs were distributed proportionally to chromosome length, chi-square goodness-of-fit tests were performed for each variant class using chromosome length proportions as expected probabilities. Per-chromosome enrichment was quantified as the ratio of observed to size-expected SV counts, where expected values were derived from each chromosome’s fractional contribution to total genome length, circumventing the density inflation artefact inherent to per-Mb normalization on smaller chromosomes. Statistical significance was evaluated using two-sided exact binomial tests, with *P*-values adjusted for multiple comparisons across 20 chromosomes using the Benjamini-Hochberg correction. Chromosomes satisfying both fold enrichment > 1 and adjusted *P* < 0.05 were classified as significantly enriched. All analyses were performed in R and visualization using ggplot2.

Orthologous gene groups across chromosome 17 of five *P. brassicae* isolates (pb3A, vic2h1, race4, 5X, and 5xst23) were extracted from OrthoFinder results. Pairwise syntenic anchors were computed using the MCscan Python implementation within JCVI (jcvi.compara.catalog ortholog), which internally invokes LAST (v1651)^105^ for sequence alignment followed by synteny analysis. Microsynteny was examined across a ∼240 kb window on chromosome 17 of the Pb3A reference, centred on effector gene PbAnk2, retaining 53 orthologous gene pairs verified across all five isolates. Syntenic blocks were visualised using jcvi.graphics.synteny.

### Pangenome saturation and gene discovery curves

Saturation curves for the pangenome graph, pan-gene set, and pan-effectorome were generated using a custom R script by computing cumulative gene counts across isolates. The resulting curves were used to evaluate gene discovery rates and assess pangenome saturation dynamics. The script is available at https://github.com/Edelab/Population-genomics-and-Pangenome-analysis.

### Yeast two-hybrid assay

Yeast two-hybrid (Y2H) screening and pairwise validation were performed by Hybrigenics Services (https://hybrigenics-services.com). The coding sequence of *P. brassicae PbAnk2* (19–460 aa, excluding the signal peptide) was synthesized (Twist Biosciences, San Fransico, California, USA), cloned into pTwist ENTR, and subcloned in-frame as a C-terminal fusion to the Gal4 DNA-binding domain in pB66 (bait), all constructs were sequence-verified. The bait was screened against a random-primed *A. thaliana* (1-week seedling) cDNA library cloned in pP6 using a mating-based approach in yeast strain CG1945. Approximately 60 million clones (∼6× library complexity) were assessed; 98 His⁺ colonies were recovered on –Trp –Leu –His medium supplemented with 50 mM 3-aminotriazole (3-AT) to suppress bait autoactivation. Prey inserts were PCR-amplified and end-sequenced, and protein identities assigned by automated BLAST against GenBank. For one-by-one assays, bait and prey plasmids were transformed into CG1945 and YHGX13, respectively, and interactions assessed via the HIS3 reporter on DO-2 (–Trp –Leu; plasmid control) and DO-3 (–Trp –Leu –His; interaction) with 3-AT titration. Negative controls included bait transformed with empty pP7 and prey transformed with empty pB66. The SMAD–SMURF interactive pair served as a positive control. Each interaction was tested as streaks of three independent clones. Experiments were performed in triplicate.

### Bimolecular fluorescence complementation

Bimolecular fluorescence complementation (BiFC) assays (ProNetBio, China) were used to test protein–protein interactions between *P. brassicae* PBTT_09611 and *A. thaliana* proteins AT5G12430 (TPR16). Coding sequences were synthesized and fused to the N- or C-terminal halves of YFP in the pCAMBIA1300-nYFP and pCAMBIA1300-cYFP vectors. Constructs were propagated in *Escherichia coli* strain TOP10 and subsequently introduced into *Agrobacterium tumefaciens* strain GV3101 by electroporation. *Nicotiana benthamiana* plants (4–6 weeks old, five-leaf stage) were grown under controlled conditions (14 h light/10 h dark photoperiod; 22–25 °C day and 18–20 °C night) and infiltrated on the abaxial leaf surface with Agrobacterium suspensions (30–50 µL per infiltration site). Leaves were harvested 36–48 h post-infiltration and imaged by laser confocal microscopy (FV4000, Olympus) to detect YFP reconstitution. All assays were performed in triplicate, with the HAI1 (AT5G59220) - MPK6 (AT2G43790) interaction used as a positive control.

### Co-immunoprecipitation assays

For co-immunoprecipitation (Co-IP) assays, *N. benthamiana* plants were grown in a greenhouse at 22–25 °C under a 16 h light/8 h dark photoperiod and used at 4–6 weeks of age (five-leaf stage). The *P. brassicae PbAnk2* was cloned into pBinGFP2 to generate a C-terminal GFP fusion, while Arabidopsis genes *TPR16* and *BG1-like* were cloned into pCAMBIA35S-4×Myc-MCS-3×FLAG. Plasmids were propagated in *E. coli* TOP10 and introduced into *A. tumefaciens* strain GV3101. Agrobacterium suspensions were syringe-infiltrated into *N. benthamiana* leaves for transient co-expression. At 36–48 h post-infiltration, leaf tissue was harvested, flash-frozen in liquid nitrogen, ground, and extracted in protein lysis buffer (1 mM DTT, 0.2% Triton X-100, 1× protease inhibitor, 50 mM HEPES, 150 mM KCl, 1 mM EDTA) at 4°C. GFP-tagged protein complexes were immunoprecipitated using GFP-trap magnetic beads at 4°C, washed with protein wash buffer (homemade, 1× PBST), and eluted with protein elution buffer (Home-made, 0.1 M Glycine and 0.15 M NaCl, pH 3.0, followed by neutralization with 1 M Tris-HCl, pH 8.0) (ProNetBio, China). Co-immunoprecipitated proteins and input controls were resolved by SDS–PAGE, transferred to PVDF membranes (Merck, Cat. No. 301004000) and detected by immunoblotting using anti-GFP (bait) (Abmart, Cat. No. M20004S) and anti-FLAG (Abmart, Cat. No. M20008S) antibodies. Chemiluminescent signals were recorded for detection.

### Statistical analysis and visualization

Non-parametric statistical analyses, including Wilcoxon, Mann–Whitney and Kruskal–Wallis tests, were performed in R and Python. Where applicable, multiple comparisons were corrected using the Benjamini–Hochberg or Bonferroni procedure, as described for individual analyses. Genomic data preprocessing, including header normalization, table joins, and QC was conducted with custom Python scripts using pandas, numpy, scikit-allele, and pysam libraries. Heatmaps and summary graphics were produced in R (ComplexHeatmap, ggplot2) and Python (matplotlib/Plotly). Complete pipelines, including reproducible code and BASH scripts are available in the EdeLab GitHub repository GitHub: https://github.com/Edelab/Population-genomics-and-Pangenome-analysis.

## ACKNOWLEDGMENTS

We gratefully acknowledge the bioinformatics and sequencing personnel at the Institut de biologie intégrative et des systèmes (IBIS), Université Laval, for their continuous support and access to essential research infrastructure throughout this project. We thank Professor Jacquie van der Waals of the University of Pretoria for generously providing *P. brassicae* isolates from South Africa. They were the first international isolates acquired by EdeLab and marked the beginning of this long and challenging research journey. We also thank our collaborators worldwide for providing pathogen isolates, and Dr. Jordan Bailey of the New South Wales Department of Primary Industries (NSW DPI) Herbarium for sharing herbarium specimens. We are grateful to the Fonds de recherche du Québec (FRQ) for the graduate funding awarded to A.J. and J.W. Finally, we thank the many undergraduate students who contributed to this project, with support from NSERC, Mitacs, and Canadian canola grower associations.

## AUTHORS CONTRIBUTION

M.A.J. led the bioinformatic analyses and wrote the original manuscript draft with F.S. and E.P.-L. F.S. performed bioinformatic analyses and contributed to data analysis and interpretation of the results. S.M. supported data analyses and contributed to the interpretation of the results. R.S. and A.-S.B. prepared the isolates for the study. E.I.F. contributed to laboratory analyses. S.M.V.C., J.W., T.M.H., A.S., Y.A., L.G.-G., S.-F.H., P.H., M.L.T.R., N.Z.-N., G.C.N., R.K., L.A., A.F., A.M., D.R.T., A.T.P.L., S.-I.I., C.A.d.S., M.G.F.d.C., U.M., G.C., J.J., M.A.C., P.T., J.D., S.C. and B.N.K. provided biological materials used in the study. M.P. and S.S. contributed to the conception and design of the study, contributed to genome sequencing and secured funding. E.P.-L. conceived and designed the study, secured funding, led and coordinated the project, supervised the research and contributed to the interpretation of the results. All authors contributed to manuscript review and editing and approved the final version.

## DATA AVAILABILITY

*Plasmodiophora brassicae* isolates are available upon reasonable request to the EdeLab by contacting. Whole-genome sequencing and genome assembly data have been deposited in the GenBank under Bioproject PRJNA1537693 and will be made publicly available upon acceptance of the manuscript. For the purpose of peer review, editors and reviewers will access the complete dataset.

## CODE AVAILABILITY

The genome analysis workflow is also available on GitHub: https://github.com/Edelab/Population-genomics-and-Pangenome-analysis. Genome raw reads and annotations will be available at NCBI after publication.

## REFERENCES

1. Stukenbrock, E. H. et al. The making of a new pathogen: Insights from comparative population genomics of the domesticated wheat pathogen *Mycosphaerella graminicola* and its wild sister species. Genome Res. 21, 2157–2166 (2011).

2. Möller, M. & Stukenbrock, E. H. Evolution and genome architecture in fungal plant pathogens. Nat. Rev. Microbiol. 15, 756–771 (2017).

3. McDonald, B. A. & Linde, C. pathogen population genetics, evolutionary potential, and durable resistance. Annu. Rev. Phytopathol. 40, 349–379 (2002).

4. Stam, R. & McDonald, B. A. When resistance gene pyramids are not durable—the role of pathogen diversity. Mol. Plant Pathol. 19, 521–524 (2018).

5. Singh, N. K., Karisto, P. & Croll, D. Population-level deep sequencing reveals the interplay of clonal and sexual reproduction in the fungal wheat pathogen *Zymoseptoria tritici*. Microb. Genom. 7, 000678 (2021).

6. Latorre, S. M. et al. Genomic surveillance uncovers a pandemic clonal lineage of the wheat blast fungus. PLoS Biol. 21, e3002052 (2023).

7. Coomber, A., Saville, A. & Ristaino, J. B. Evolution of Phytophthora infestans on its potato host since the Irish potato famine. Nat. Commun. 15, 6488 (2024).

8. Li, Y. et al. Population dynamics of Phytophthora infestans in the Netherlands reveals expansion and spread of dominant clonal lineages and virulence in sexual offspring. *G3* (*Bethesda**) 2*, 1529–1540 (2012).

9. Matthews, C. A., Watson-Haigh, N. S., Burton, R. A. & Sheppard, A. E. A gentle introduction to pangenomics. Brief Bioinform. 25, bbae588 (2024).

10. Badet, T., Oggenfuss, U., Abraham, L., McDonald, B. A. & Croll, D. A 19-isolate reference-quality global pangenome for the fungal wheat pathogen *Zymoseptoria tritici*. BMC Biol. *2020* 18:*1* *18*, 1–18 (2020).

11. Fouché, S., Oggenfuss, U., Chanclud, E. & Croll, D. A devil’s bargain with transposable elements in plant pathogens. Trends in Genet. 38, (2022).

12. Abraham, L. N., Oggenfuss, U. & Croll, D. Population-level transposable element expression dynamics influence trait evolution in a fungal crop pathogen. mBio. 15, e0284023 (2024).

13. Grandaubert, J., Dutheil, J. Y. & Stukenbrock, E. H. The genomic determinants of adaptive evolution in a fungal pathogen. Evol. Letters 3, 299–312 (2019).

14. Schwelm, A. et al. Not in your usual Top 10: protists that infect plants and algae. Mol. Plant Pathol. 19, 1029–1044 (2018).

15. Schwelm, A. et al. The Plasmodiophora brassicae genome reveals insights in its life cycle and ancestry of chitin synthases. Sci. Rep. 5, 11153 (2015).

16. Javed, M. A., Mukhopadhyay, S., Normandeau, E., Brochu, A.-S. & Pérez-López, E. Telomere-to-telomere Genome Assembly of the Clubroot Pathogen *Plasmodiophora Brassicae*. Genome. Biol. Evol. 16, evae122 (2024).

17. Javed, M. A. et al. The clubroot pathogen Plasmodiophora brassicae: A profile update. Mol. Plant Pathol. 24, 89–106 (2023).

18. Hwang, S. F., Howard, R. J., Strelkov, S. E., Gossen, B. D. & Peng, G. Management of clubroot (*Plasmodiophora brassicae*) on canola (*Brassica napus L*.) in western Canada. Can. J. Plant Pathol. 36, 49–65 (2014).

19. Zamani-Noor, N. Variation in pathotypes and virulence of Plasmodiophora brassicae populations in Germany. Plant Pathol. 66, 316–324 (2017).

20. Hollman, K. B., Hwang, S. F., Manolii, V. P. & Strelkov, S. E. Pathotypes of Plasmodiophora brassicae collected from clubroot resistant canola (*Brassica napus L*.) cultivars in western Canada in 2017-2018. Can. J. Plant Pathol. 43, 622–630 (2021).

21. Wu, J., Mukhopadhyay, S. & Pérez-López, E. Déjà vu in clubroot resistance: same genes, new names. Trends in Genet. 42, 11–13 (2026).

22. Zamani-Noor, N. et al. Pathotype Characterization of *Plasmodiophora brassicae*, the cause of clubroot in central Europe and Sweden (2016–2020). Patho. 11, 1440 (2022).

23. Mukhopadhyay, S., Javed, M. A., Wu, J. & Perez-Lopez, E. Structure-guided secretome analysis of gall-forming microbes offers insights into effector diversity and evolution. eLife 14, RP105185 (2025).

24. Liu, L. et al. Refining the life cycle of *Plasmodiophora brassicae*. Phytopathol. 110, 1704–1712 (2020).

25. Tommerup, I. C. & Ingram, D. S. The life-cycle of *Plasmodiophora brassicae* woron. in *brassica* tissue cultures and in intact roots. New Phytol. 70, 327–332 (1971).

26. Li, H. et al. Pathogen protein modularity enables elaborate mimicry of a host phosphatase. Cell 186, 3196–3207 (2023).

27. Figueroa, M., Ortiz, D. & Henningsen, E. C. Tactics of host manipulation by intracellular effectors from plant pathogenic fungi. Curr. Opin. Plant Biol. 62, 102054 (2021).

28. Fabro, G. Oomycete intracellular effectors: specialised weapons targeting strategic plant processes. New Phytol. 233, 1074–1082 (2022).

29. Toruño, T. Y., Stergiopoulos, I. & Coaker, G. Plant-pathogen effectors: cellular probes interfering with plant defenses in spatial and temporal manners. Annu. Rev. Phytopathol. 54, 419–441 (2016).

30. Bauters, L., Stojilković, B. & Gheysen, G. Pathogens pulling the strings: Effectors manipulating salicylic acid and phenylpropanoid biosynthesis in plants. Mol. Plant Pathol. 22, 1436–1448 (2021).

31. Liu, Y., Zhang, Y.-M., Tang, Y., Chen, J.-Q. & Shao, Z.-Q. The evolution of plant NLR immune receptors and downstream signal components. Curr. Opin. Plant Biol. 73, 102363 (2023).

32. Martel, A. et al. The ETS-ETI cycle: evolutionary processes and metapopulation dynamics driving the diversification of pathogen effectors and host immune factors. Curr. Opin. Plant Biol. 62, 102011 (2021).

33. Sperschneider, J. et al. Advances and challenges in computational prediction of effectors from plant pathogenic fungi. PLOS Patho. 11, e1004806 (2015).

34. Sánchez-Vallet, A. et al. The genome biology of effector gene evolution in filamentous plant pathogens. Annu. Rev. Phytopathol. 56, 21–40 (2018).

35. Stjelja, S., Fogelqvist, J., Tellgren-Roth, C. & Dixelius, C. The architecture of the *Plasmodiophora brassicae* nuclear and mitochondrial genomes. Sci. Rep. 9, 15753 (2019).

36. Tettelin, H. et al. Genome analysis of multiple pathogenic isolates of *Streptococcus agalactiae*: implications for the microbial ‘pan-genome’. Proc. Natl. Acad. Sci. 102, 13950–13955 (2005).

37. Hickey, G. et al. Pangenome graph construction from genome alignments with Minigraph-Cactus. Nat. Biotechnol. 42, 663–673 (2024).

38. Skiadas, P. et al. Pangenome graph analysis reveals extensive effector copy-number variation in spinach downy mildew. PLoS Genet. 20, e1011452 (2024).

39. Brefort, T., et al. Characterization of the largest effector gene cluster of *Ustilago maydis*. PLoS Patho. 10, e1003866 (2014).

40. Skiadas, P. et al. Pangenome graph analysis reveals evolution of resistance breaking in spinach downy mildew. PLoS Biol. 24, e3003596 (2026).

41. Fontyn, C. et al. Evolution within a given virulence phenotype (pathotype) is driven by changes in aggressiveness: a case study of French wheat leaf rust populations. Peer Community Journal 3, e39 (2023).

42. Jones, J. D. G. & Dangl, J. L. The plant immune system. Nature *2006* 444:*7117 444*, 323–329 (2006).

43. Yoshida, K. et al. The rise and fall of the Phytophthora infestans lineage that triggered the Irish potato famine. Elife 2, e00731 (2013).

44. Dong, S., Raffaele, S. & Kamoun, S. The two-speed genomes of filamentous pathogens: waltz with plants. Curr Opin Genet Dev 35, 57–65 (2015).

45. Torres, D. E., Oggenfuss, U., Croll, D. & Seidl, M. F. Genome evolution in fungal plant pathogens: looking beyond the two-speed genome model. Fungal Biol. Rev. 34, 136–143 (2020).

46. Feurtey, A. et al. Genome compartmentalization predates species divergence in the plant pathogen genus Zymoseptoria. BMC Genomics 21, 588 (2020).

47. Frantzeskakis, L., Kusch, S. & Panstruga, R. The need for speed: compartmentalized genome evolution in filamentous phytopathogens. Mol. Plant Pathol. 20, 3–7 (2019).

48. Li, J. et al. Variability in Chromosome 1 of Select Moroccan *Pyrenophora teres f. teres* Isolates overcomes a highly effective barley chromosome 6H source of resistance. *Mol*. Plant-Microbe Interactions 37, 676–687 (2024).

49. Rocafort, M. et al. The Venturia inaequalis effector repertoire is dominated by expanded families with predicted structural similarity, but unrelated sequence, to avirulence proteins from other plant-pathogenic fungi. BMC Biol. 20, 246 (2022).

50. Pang, W., Liang, Y., Zhan, Z., Li, X. & Piao, Z. Development of a sinitic clubroot differential set for the pathotype classification of *Plasmodiophora brassicae*. Front. Plant Sci. 11, 1360 (2020).

51. Williams, P. H. A system for the determination of races of *Plasmodiophora brassicae* that infect cabbage and rutabaga. Phytopathol. 56, 624–626 (1966).

52. Strelkov, S. E. et al. Virulence and pathotype classification of *Plasmodiophora brassicae* populations collected from clubroot resistant canola (*Brassica napus L.*) in Canada. Can. J. Plant Pathol. 40, 284–298 (2018).

53. Some, A. et al. Variation for virulence on *Brassica napus L*. amongst *Plasmodiophora brassicae* collections from France and derived single-spore isolates. Plant Pathol. 45, 432–439 (1996).

54. Pérez-López, E., Hossain, M. M., Wei, Y., Todd, C. D. & Bonham-Smith, P. C. A clubroot pathogen effector targets cruciferous cysteine proteases to suppress plant immunity. Virulence 12, 2327–2340 (2021).

55. Jelenska, J., Van Hal, J. A. & Greenberg, J. T. *Pseudomonas syringae* hijacks plant stress chaperone machinery for virulence. Proc. Natl. Acad. Sci. U.S.A. 107, 13177–13182 (2010).

56. Jing, M. et al. A *Phytophthora sojae* effector suppresses endoplasmic reticulum stress-mediated immunity by stabilizing plant Binding immunoglobulin Proteins. Nat. Commun. 7, 11685 (2016).

57. Nakano, M., Ichinose, Y. & Mukaihara, T. *Ralstonia solanacearum* Type III effector RipAC targets SGT1 to suppress effector-triggered immunity. Plant and Cell Physiol. 61, 2067–2076 (2021).

58. Salih, R., et al. A Basic Guide to the Propagation and Manipulation of the Clubroot Pathogen, Plasmodiophora brassicae. Curr. Protoc. 4, e1039 (2024).

59. Askarian, H. et al. Genetic structure of *Plasmodiophora brassicae* populations virulent on clubroot resistant canola (*Brassica napus L*.). Plant Dis. 105, 3694–3704 (2021).

60. Jones, A. & Schwessinger, B. Sorbitol washing complex homogenate for improved DNA extractions v1. Preprint at 10.17504/protocols.io.beuvjew6 (2020).

61. LaMar, D. FastQC. https://qubeshub.org/resources/fastqc (2015).

62. Bolger, A. M., Lohse, M. & Usadel, B. Trimmomatic: a flexible trimmer for Illumina sequence data. Bioinformatics 30, 2114–2120 (2014).

63. Li, H. & Durbin, R. Fast and accurate short read alignment with Burrows-Wheeler transform. Bioinformatics 25, 1754–1760 (2009).

64. Tarasov, A., Vilella, A. J., Cuppen, E., Nijman, I. J. & Prins, P. Sambamba: fast processing of NGS alignment formats. Bioinformatics 31, 2032–2034 (2015).

65. Li, H. et al. The Sequence Alignment/Map format and SAMtools. Bioinformatics 25, 2078–2079 (2009).

66. Danecek, P. & McCarthy, S. A. BCFtools/csq: haplotype-aware variant consequences. Bioinformatics 33, 2037–2039 (2017).

67. Danecek, P. et al. The variant call format and VCFtools. Bioinformatics 27, 2156–2158 (2011).

68. Cingolani, P. et al. A program for annotating and predicting the effects of single nucleotide polymorphisms, SnpEff: SNPs in the genome of Drosophila melanogaster strain w1118; iso-2; iso-3. *Fly 6*, 80–92 (2012).

69. Chang, C. C. et al. Second-generation PLINK: rising to the challenge of larger and richer datasets. Gigascience 4, 7 (2015).

70. Purcell, S. et al. PLINK: a tool set for whole-genome association and population-based linkage analyses. Am. J. Hum. Genet. 81, 559–575 (2007).

71. Venables, W. N. & Ripley, B. D. *Modern Applied Statistics with S*. (Springer New York, New York, NY, 2002). doi:10.1007/978-0-387-21706-2.

72. Wickham, H. ggplot2. WIREs Computational Stats 3, 180–185 (2011).

73. Liu, X., Kopelman, N. M. & Rosenberg, N. A. *Clumppling*: cluster matching and permutation program with integer linear programming. Bioinformatics 40, btad751 (2024).

74. Dixon, P. VEGAN, a package of R functions for community ecology. J. Vegetation Sci. 14, 927–930 (2003).

75. Emms, D. M. & Kelly, S. OrthoFinder: phylogenetic orthology inference for comparative genomics. Genome Biol. 20, 238 (2019).

76. Katoh, K. & Standley, D. M. MAFFT multiple sequence alignment software version 7: improvements in performance and usability. Mol. Biol. and Evol. 30, 772–780 (2013).

77. Minh, B. Q. et al. IQ-TREE 2: New Models and Efficient Methods for Phylogenetic Inference in the Genomic Era. Mol. Biol. Evol. 37, 1530–1534 (2020).

78. Letunic, I. & Bork, P. Interactive Tree of Life (iTOL) v6: recent updates to the phylogenetic tree display and annotation tool. Nucleic Acids Res. 52, W78–W82 (2024).

79. Huson, D. H. & Bryant, D. The SplitsTree App: interactive analysis and visualization using phylogenetic trees and networks. Nat. Methods 21, 1773–1774 (2024).

80. Bruen, T. C., Philippe, H. & Bryant, D. A Simple and Robust Statistical Test for Detecting the Presence of Recombination. Genetics 172, 2665–2681 (2006).

81. Tso, H. H., Galindo-González, L., Locke, T. & Strelkov, S. E. Protocol: rhPCR and SNaPshot assays to distinguish *Plasmodiophora brassicae* pathotype clusters. Plant Methods 18, 91 (2022).

82. Askarian, H. et al. Virulence Spectrum of Single-Spore and Field Isolates of *Plasmodiophora brassicae* Able to Overcome Resistance in Canola (*Brassica napus* L.). Plant Disease 105, 43–52 (2021).

83. Korunes, K. L. & Samuk, K. PIXY: Unbiased estimation of nucleotide diversity and divergence in the presence of missing data. *Mol*. Ecol. Res. 21, 1359–1368 (2021).

84. Amezrou, R. et al. Quantitative pathogenicity and host adaptation in a fungal plant pathogen revealed by whole-genome sequencing. Nat. Commun. 15, 1933 (2024).

85. Rougemont, Q. et al. Demographic history shaped geographical patterns of deleterious mutation load in a broadly distributed Pacific Salmon. PLoS Genet. 16, e1008348 (2020).

86. De Coster, W., D’Hert, S., Schultz, D. T., Cruts, M. & Van Broeckhoven, C. NanoPack: visualizing and processing long-read sequencing data. Bioinformatics 34, 2666–2669 (2018).

87. De Coster, W. & Rademakers, R. NanoPack2: population-scale evaluation of long-read sequencing data. Bioinformatics 39, btad311 (2023).

88. Cheng, H., Concepcion, G. T., Feng, X., Zhang, H. & Li, H. Haplotype-resolved de novo assembly using phased assembly graphs with hifiasm. Nat. Methods 18, 170–175 (2021).

89. Koren, S. et al. Canu: scalable and accurate long-read assembly via adaptive k-mer weighting and repeat separation. Genome Res. 27, 722–736 (2017).

90. Astashyn, A. et al. Rapid and sensitive detection of genome contamination at scale with FCS-GX. Genome Biol. 25, 60 (2024).

91. Aury, J.-M. & Istace, B. Hapo-G, haplotype-aware polishing of genome assemblies with accurate reads. NAR Genomics and Bioinformatics 3, lqab034 (2021).

92. Gurevich, A., Saveliev, V., Vyahhi, N. & Tesler, G. QUAST: quality assessment tool for genome assemblies. Bioinformatics 29, 1072–1075 (2013).

93. Huang, N. & Li, H. compleasm: a faster and more accurate reimplementation of BUSCO. Bioinformatics 39, btad595 (2023).

94. Cabanettes, F. & Klopp, C. D-GENIES: dot plot large genomes in an interactive, efficient and simple way. PeerJ 6, e4958 (2018).

95. Li, H. Minimap2: pairwise alignment for nucleotide sequences. Bioinformatics 34, 3094–3100 (2018).

96. Gabriel, L. et al. BRAKER3: Fully automated genome annotation using RNA-seq and protein evidence with GeneMark-ETP, AUGUSTUS, and TSEBRA. Genome Res. 34, 769–777 (2024).

97. Baril, T., Galbraith, J. & Hayward, A. Earl Grey: A Fully Automated User-Friendly Transposable Element Annotation and Analysis Pipeline. Mol. Biol. Evol. 41, msae068 (2024).

98. Lovelace, A. H. et al. Effector Identification in Plant Pathogens. Phytopathol. 113, 637–650 (2023).

99. Teufel, F. et al. SignalP 6.0 predicts all five types of signal peptides using protein language models. Nat. Biotechnol. 40, 1023–1025 (2022).

100. Hallgren, J. et al. DeepTMHMM predicts alpha and beta transmembrane proteins using deep neural networks. bioRxiv 10.1101/2022.04.08.487609 (2022).

101. Zhao, K. et al. Pangenome analysis reveals structural variation associated with seed size and weight traits in peanut. Nat. Genet. 57, 1250–1261 (2025).

102. Wang, T. et al. The Human Pangenome Project: a global resource to map genomic diversity. Nature 604, 437–446 (2022).

103. Wick, R. R., Schultz, M. B., Zobel, J. & Holt, K. E. Bandage: interactive visualization of de novo genome assemblies. Bioinformatics 31, 3350–3352 (2015).

104. Quinlan, A. R. & Hall, I. M. BEDTools: a flexible suite of utilities for comparing genomic features. Bioinformatics 26, 841–842 (2010).

105. Kiełbasa, S. M., Wan, R., Sato, K., Horton, P. & Frith, M. C. Adaptive seeds tame genomic sequence comparison. Genome Res. 21, 487–493 (2011).

