## Supplementary Figures for "Global population genomics and pangenome of the clubroot pathogen link clonal expansion and effector diversification to resistance breakdown in canola"

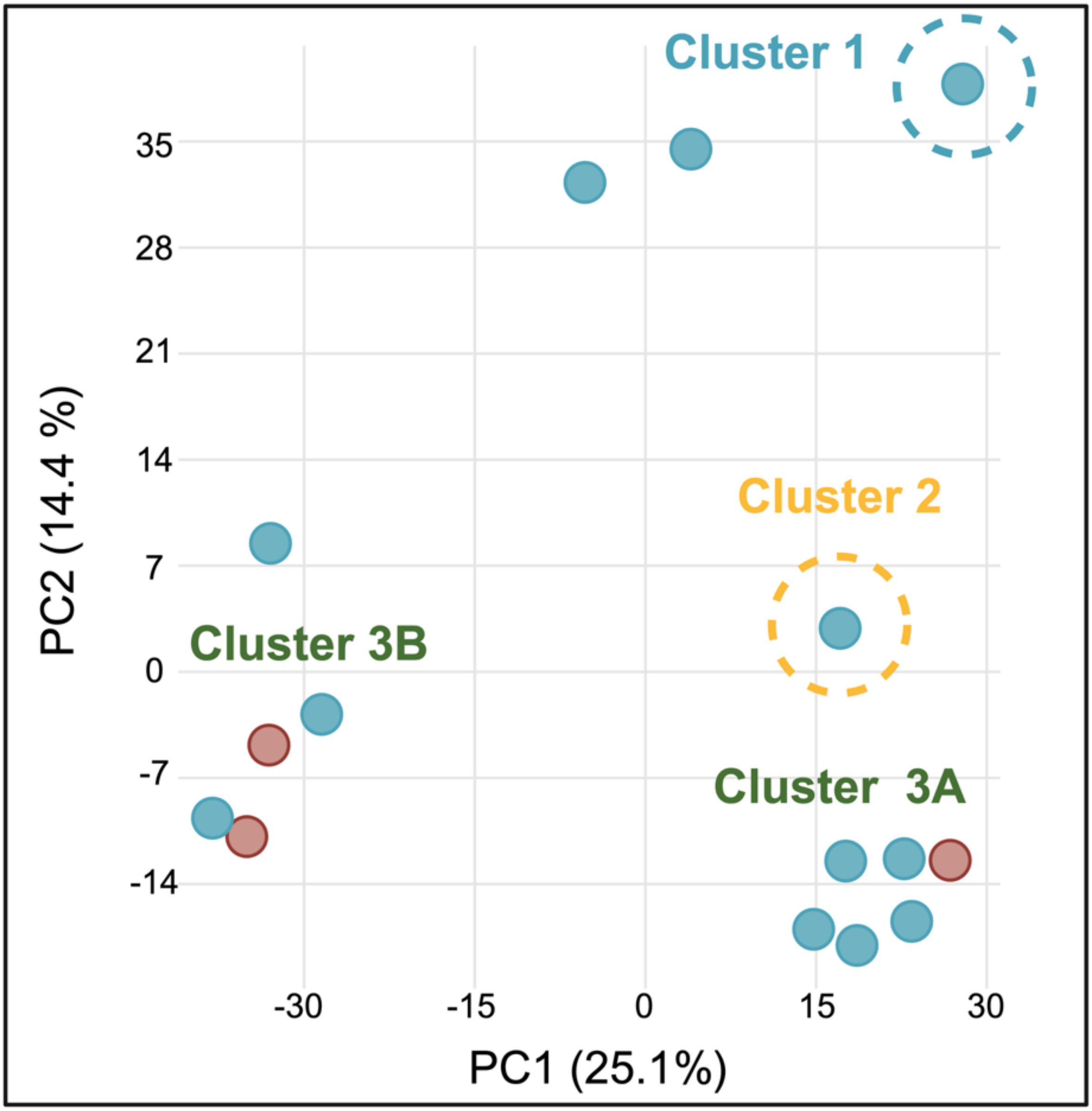


**Supplementary Fig. 1. Principal component analysis of the unrelated isolates panel**. PCA of 15 *P. brassicae* isolates retained after removing near-identical isolates (>90% IBS), based on LD-pruned genome-wide dataset. Axes show PC1 and PC2, with the percentage of variance explained. Clusters 1, 2, 3A and 3B are indicated; dashed circles mark the single representatives of the clonally expanded Cluster 1 and Cluster 2 lineages.

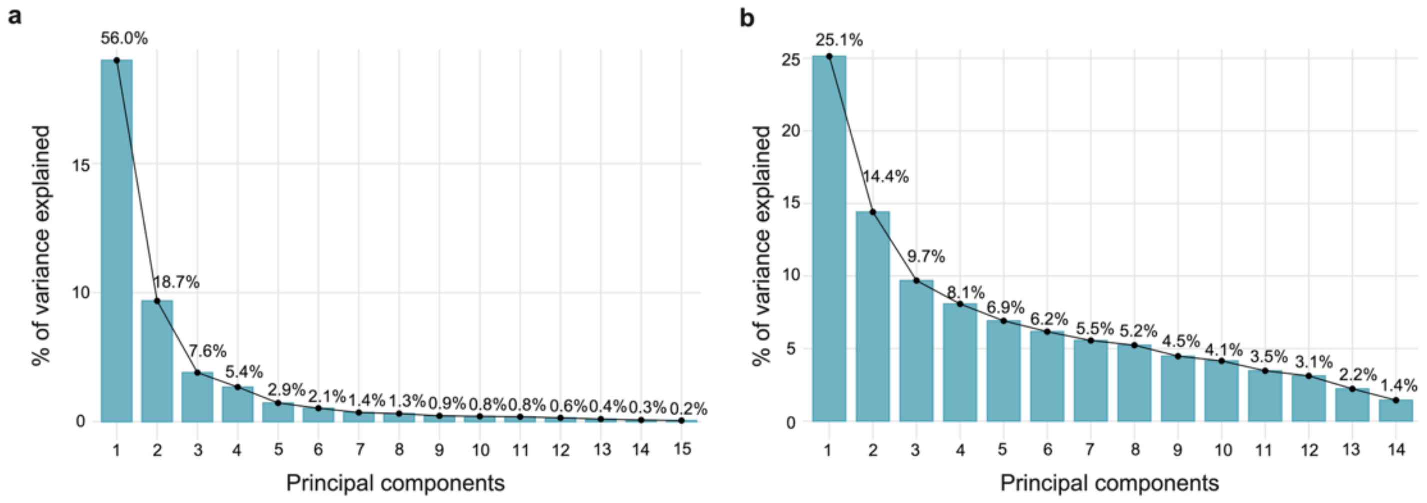


**Supplementary Fig. 2. Scree plots of variance explained by principal components.** Percentage of genetic variance captured by successive principal components for (**a**) the full 141-isolate panel and (**b**) the 15-isolate unrelated panel (after removing near-identical isolates, >90% IBS), both computed from LD-pruned SNPs. Bars and the overlaid line show per-PC variance. PC1 declines from 56.0% to 25.1% after removing over representative isolates, reflecting the removal of clonal redundancy.

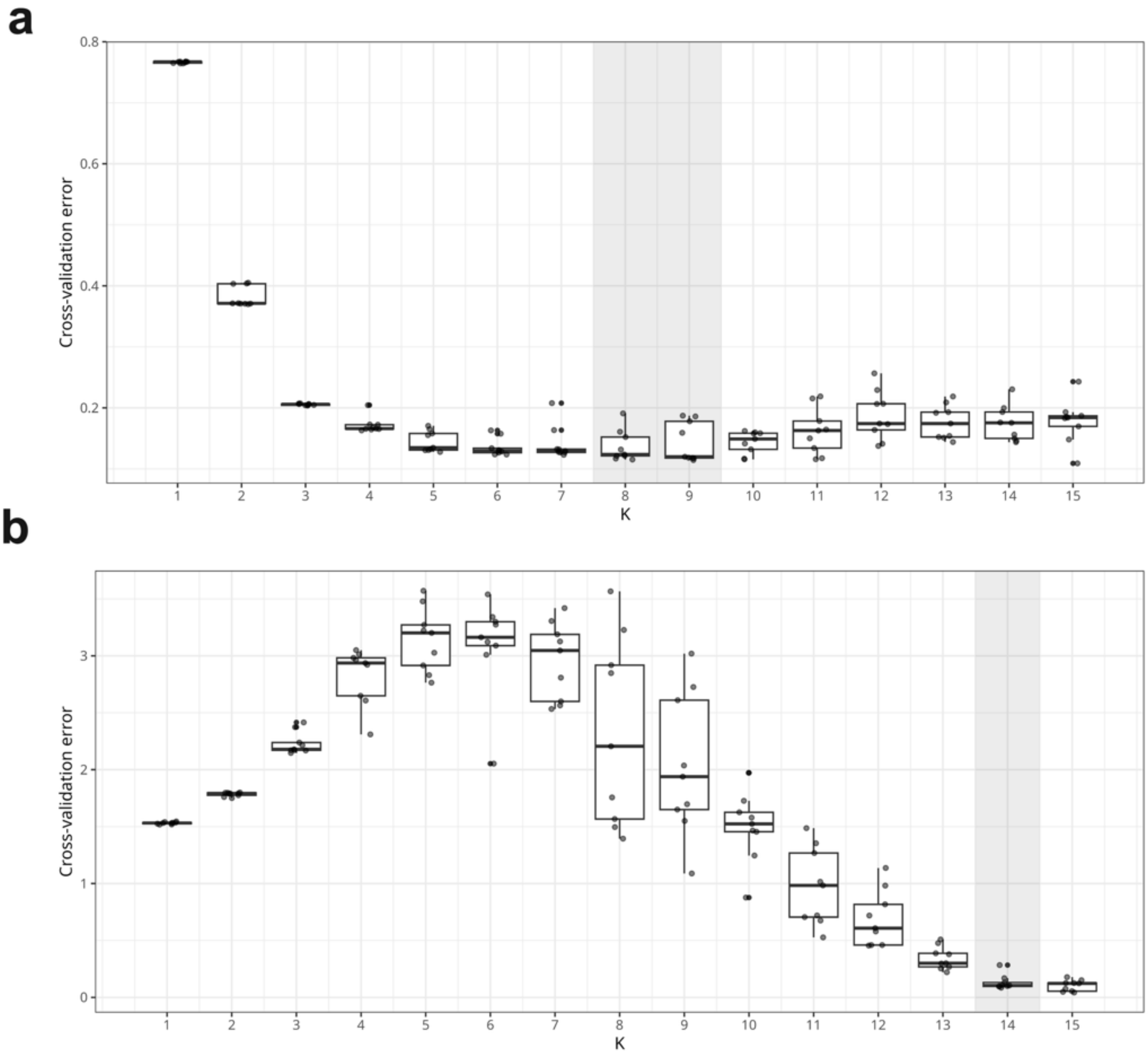


**Supplementary Fig. 3. ADMIXTURE cross-validation error. Cross-validation (CV) error across *K* = 1–15 for (a)** the 141-isolate panel and **(b)** the 15-isolate clone-corrected panel; box plots summarise ten replicate runs per *K*, with individual replicates shown as points. In (a), CV error is minimised at *K* = 8, with *K* = 9 essentially tied (shaded), and *K* = 8 was used for fine-scale structure. In (b), CV error does not identify an interpretable optimum (rising, then falling to a minimum at *K* = 14, shaded); *K* was instead selected on the basis of mode stability across replicates.

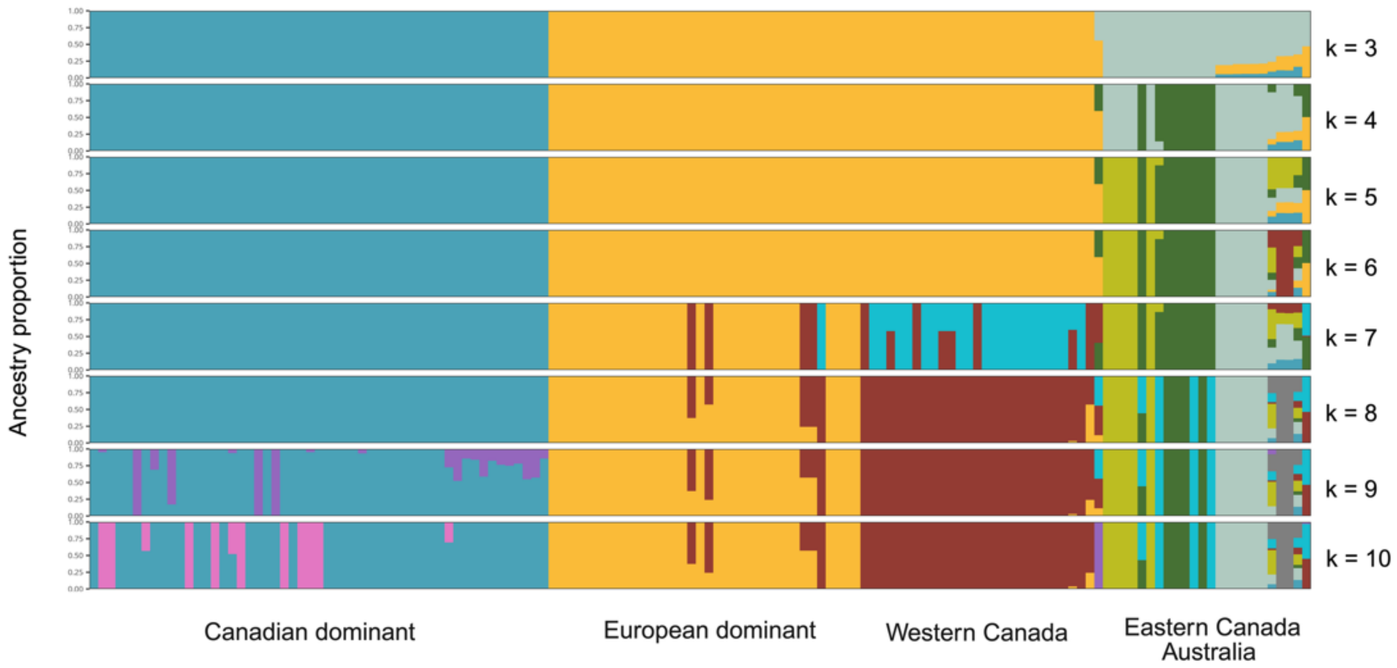


**Supplementary Fig. 4. ADMIXTURE ancestry proportions for K = 3 to K = 10.** Each horizontal bar represents an individual, partitioned into K colored segments showing the proportion of its genome inferred to derive from each of K ancestral clusters, estimated using ADMIXTURE for K = 3 through K = 10 (top to bottom). Individuals are ordered identically across all panels and grouped according to four broad population categories (Canadian dominant, European dominant, Western Canada, and Eastern Canada/Australia), indicated below the plot. At low K, isolates resolve into two major ancestral components broadly corresponding to Canadian- and European-dominant groups, with additional structure emerging within and among the Western Canada and Eastern Canada/Australia groups at higher K values.

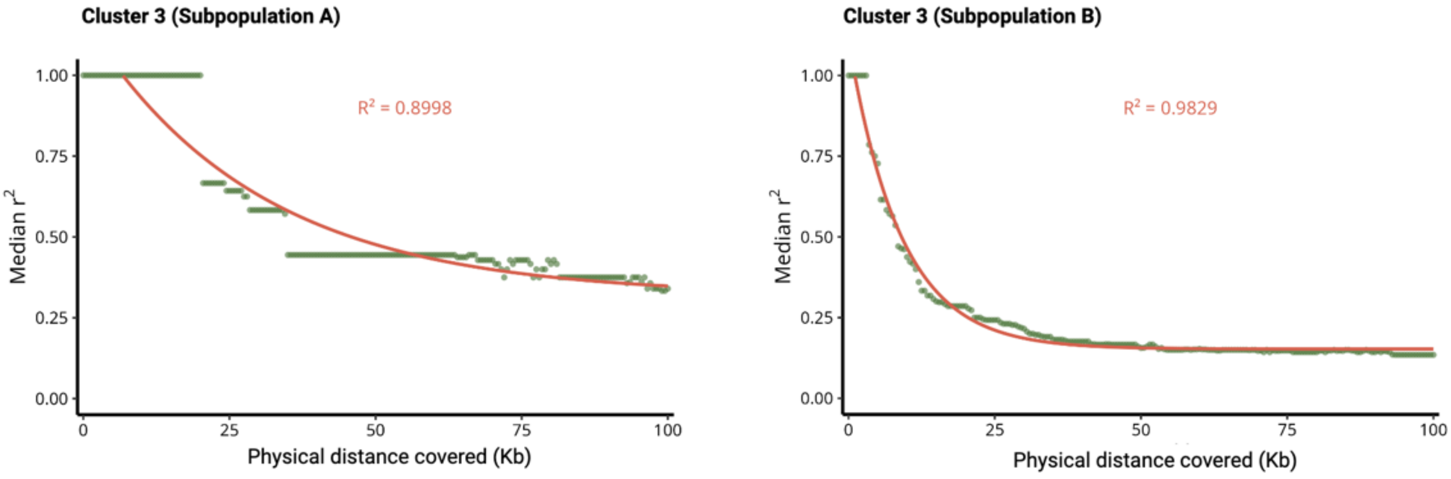


**Supplementary Fig. 5. Linkage disequilibrium decay within the Cluster 3 subpopulations.** Pairwise linkage disequilibrium ($r^2$) plotted with median $r^2$ as a function of physical distance for the two subpopulations identified within Cluster 3 (Subpopulation A and Subpopulation B). The fitted nonlinear decay curve (red line) and corresponding coefficient of determination ($R^2$) shown for each subpopulation. LD decays more rapidly in Cluster 3 Subpopulation B than in Subpopulation A.

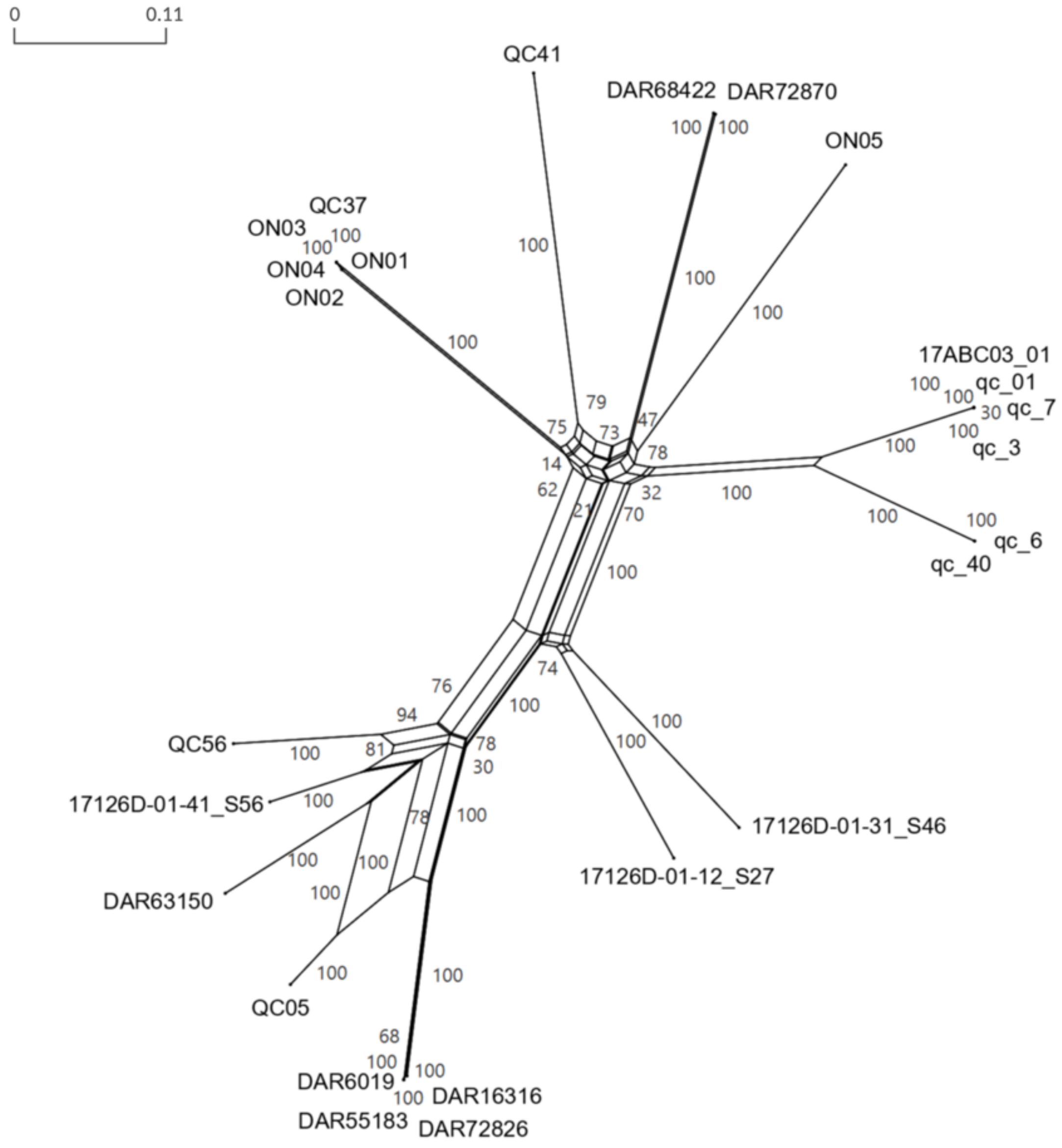


**Supplementary Fig. 6. Phylogenetic network of Cluster 3 isolates showing evidence of recombination.**Neighbor-net phylogenetic network constructed from genome-wide SNP data for isolates within Cluster 3, generated using SplitsTree (fit = 99.9%). Reticulations (parallel edges and boxed/netted structures) rather than a strictly bifurcating tree topology indicate signals of recombination among isolates, in contrast to a purely clonal. Branch lengths are proportional to genetic distance, as indicated by the scale bar (0–0.099). Isolates form several distinct subgroups connected through reticulate edges, consistent with ongoing or historical recombination within this cluster.

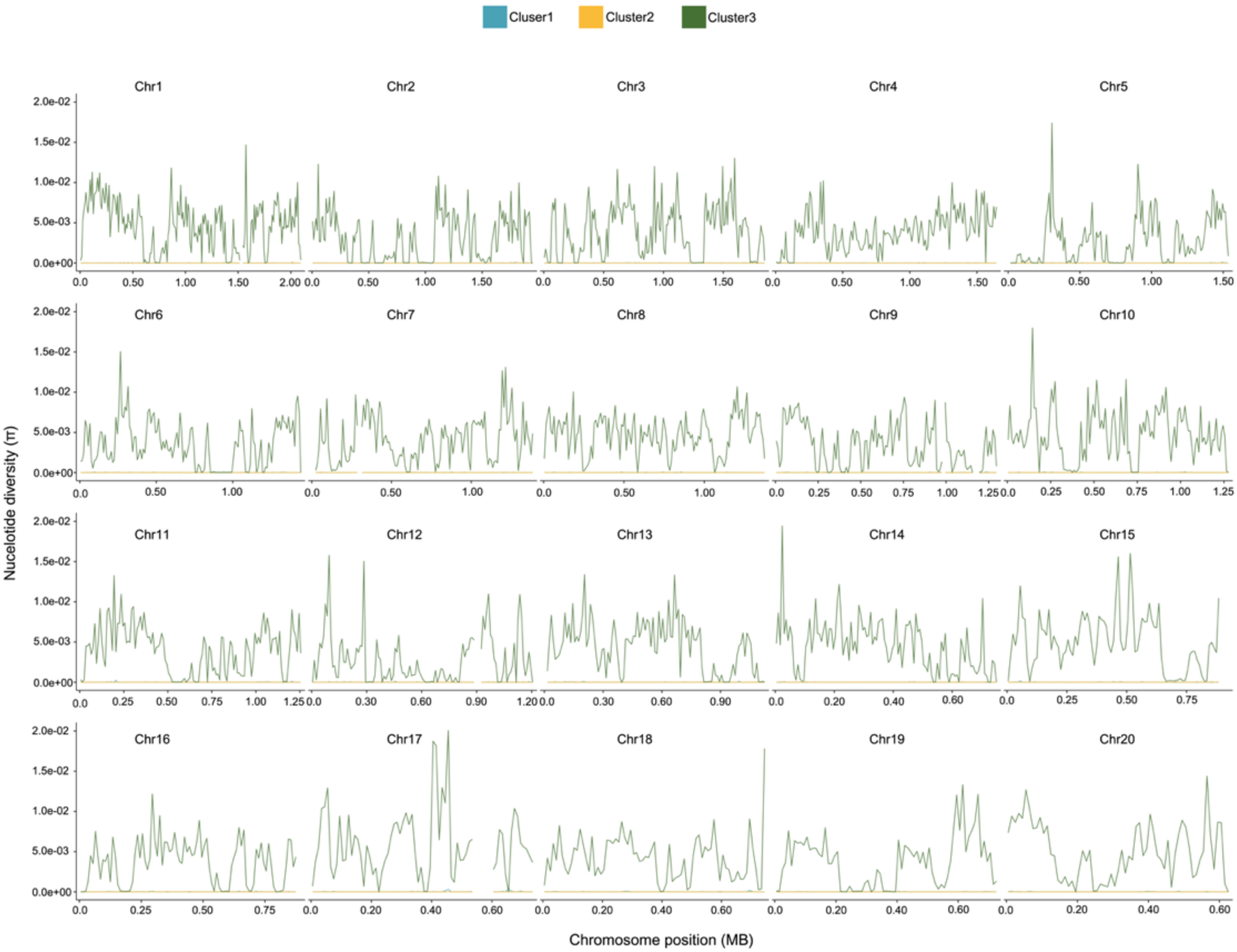


**Supplementary Fig. 7. Genome-wide nucleotide diversity (π) across all 20 chromosomes for three population clusters.** Line plots showing nucleotide diversity (π) calculated in sliding windows across all 20 chromosomes for isolates belonging to Cluster 1 (teal), Cluster 2 (yellow), and Cluster 3 (dark green). Each panel represents one chromosome, with chromosome position (Mb) on the x-axis and π on the y-axis (scale 0–2.0 × $10^{-2}$). Cluster 3 consistently exhibits the highest π across all chromosomes, reflecting substantially greater within-cluster genetic diversity relative to Clusters 1 and 2. Cluster 1 and Cluster 2 show near-zero π genome-wide, consistent with low within-cluster diversity and indicative of recent clonal expansion in these lineages. Localized peaks of elevated diversity in Cluster 3 are distributed across all chromosomes, with Chr17 showing a disproportionate concentration of diverse regions.

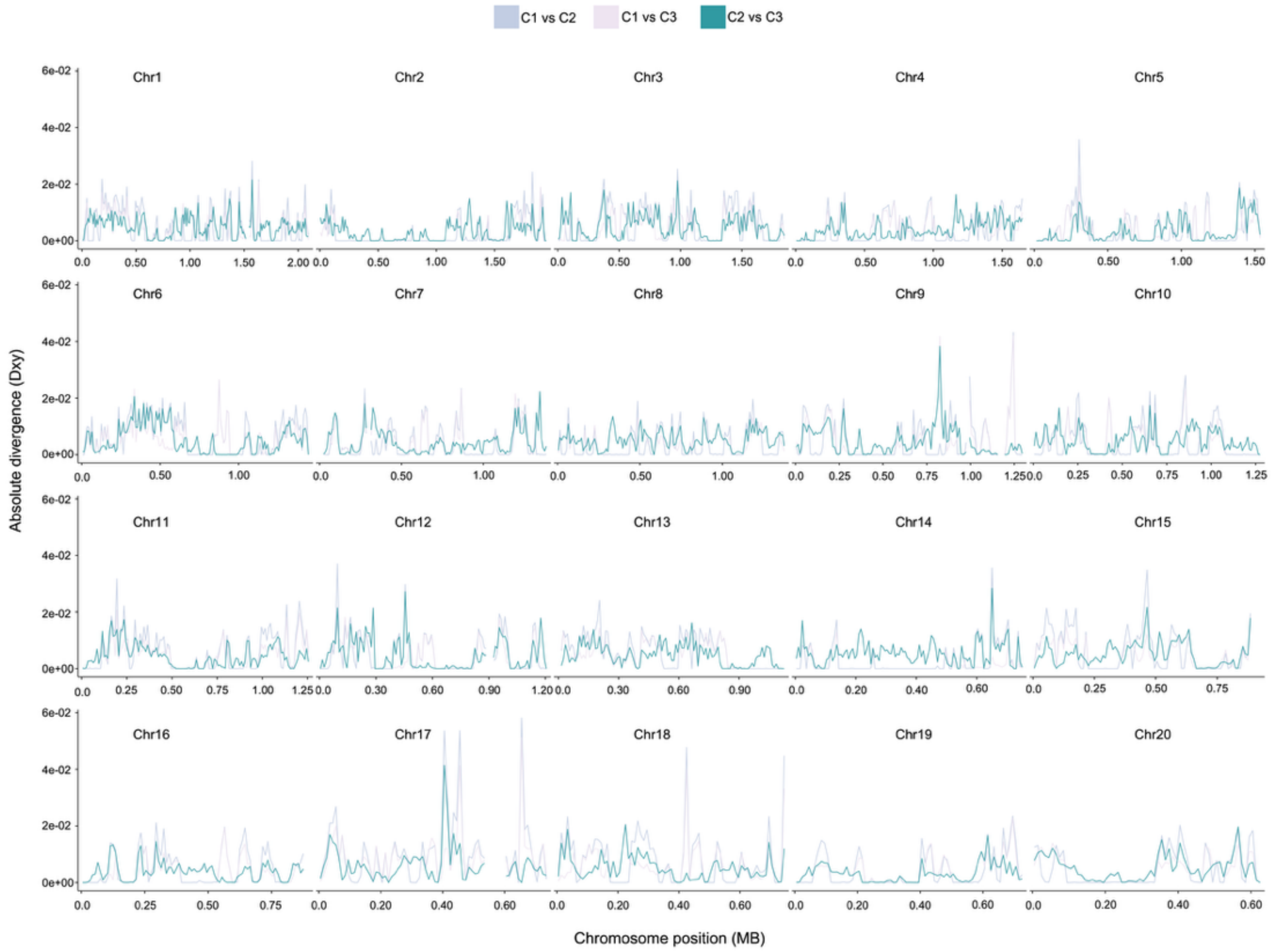


**Supplementary Fig. 8. Genome-wide absolute sequence divergence (dxy) across all 20 chromosomes for pairwise cluster comparisons.** Line plots showing absolute divergence (dxy) calculated in sliding windows across all 20 chromosomes for each pairwise comparison among the three population clusters: Cluster 1 vs Cluster 2, Cluster 1 vs Cluster 3, and Cluster 2 vs Cluster 3. Each panel represents one chromosome, with chromosome position (Mb) on the x-axis and dxy on the y-axis (scale 0–6.0 × $10^{-2}$). The Cluster 2 vs Cluster 3 comparison consistently shows the highest dxy across most chromosomes, indicating the greatest absolute divergence between these two clusters genome wide. Cluster 1 vs Cluster 2 and Cluster 1 vs Cluster 3 comparisons show broadly similar and overlapping dxy values across most chromosomes, suggesting comparable divergence of Cluster 1 from both Cluster 2 and Cluster 3. Prominent localized peaks of elevated dxy are visible across multiple chromosomes, particularly on Chr9, Chr12, Chr14, and Chr17, potentially indicating regions of elevated differentiation among clusters.

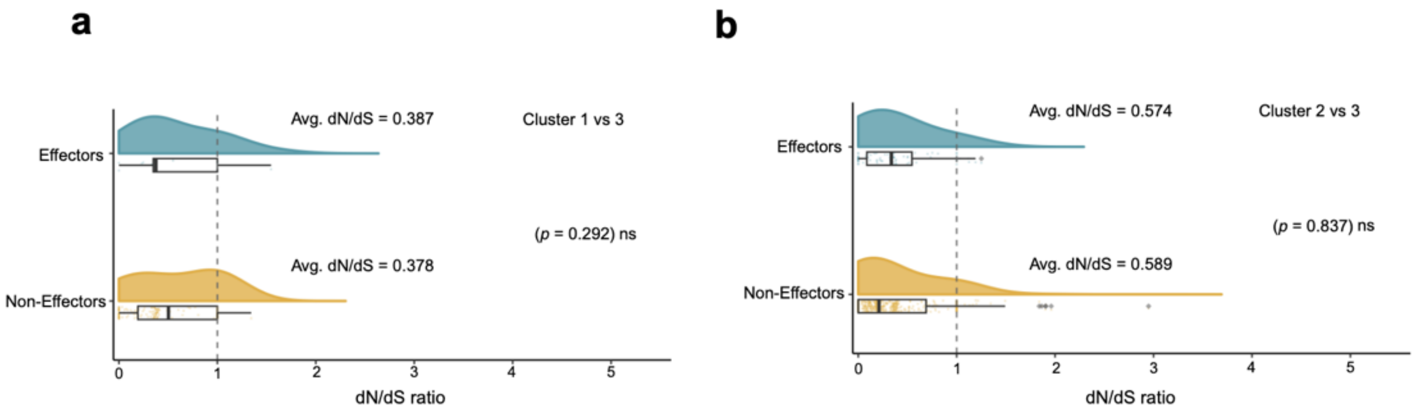


**Supplementary Fig. 9. Selective pressure (dN/dS) on effector and non-effector genes between population clusters.** Raincloud plots (combined density distribution, boxplot, and jittered individual data points) showing the distribution of pairwise dN/dS ratios for predicted effector and non-effector genes, compared between **(a)** Cluster 1 vs Cluster 3 and **(b)** Cluster 2 vs Cluster 3. The dashed vertical line at dN/dS = 1 indicates the neutral expectation; values below 1 indicate purifying selection. Mean dN/dS values are shown for each gene category, and *P*-values from comparison between effector and non-effector distributions are reported (ns, not significant; Mann-Whitney U test or equivalent). Effector and non-effector genes show similarly low average dN/dS in both comparisons, with no significant difference between gene categories.

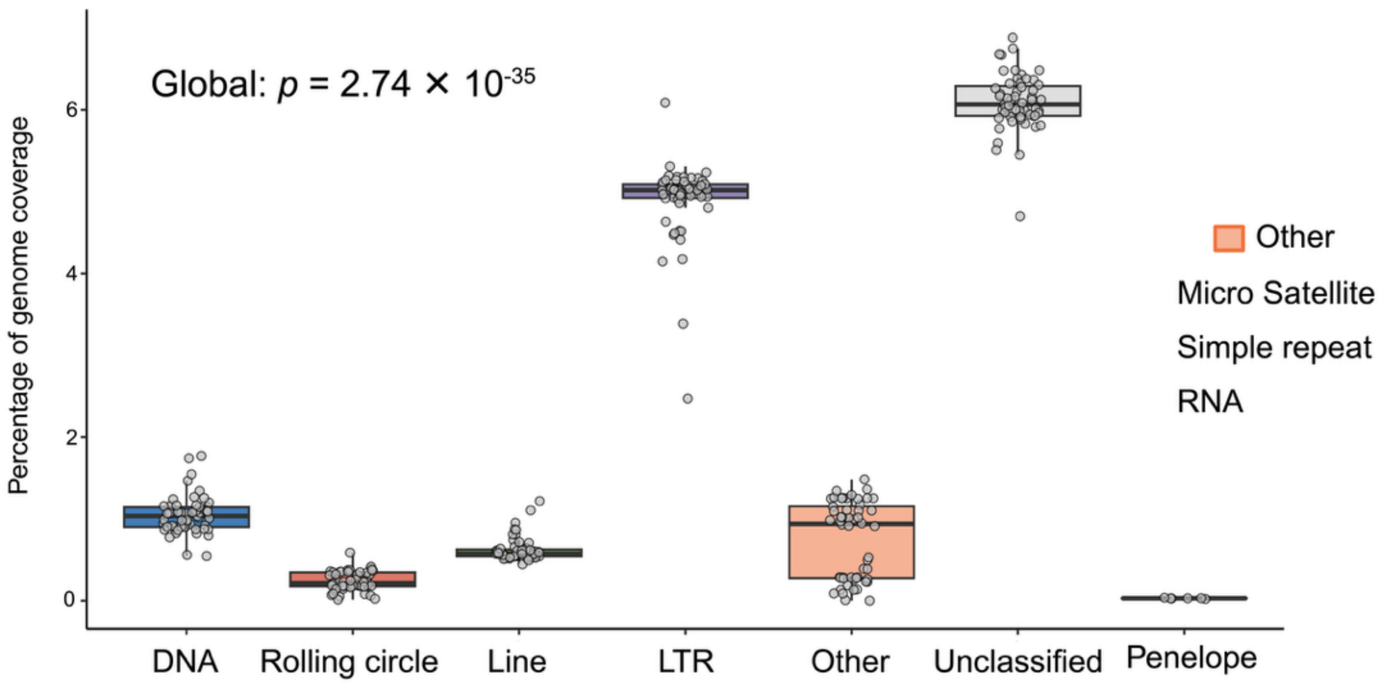


**Supplementary Fig. 10. Genome-wide proportions of transposable element (TE) classes among pangenome isolates**. Boxplots showing the percentage of genome coverage attributed to each major transposable element class (DNA transposons, rolling circle, LINE, LTR retrotransposons, Other, Unclassified, and Penelope) across all isolates, with individual isolate values overlaid as jittered points. A global Kruskal-Wallis test indicates highly significant differences in genome coverage among TE classes ($P = 2.74 \times 10^{-35}$). Unclassified repeats and LTR retrotransposons together account for the largest share of genome coverage.

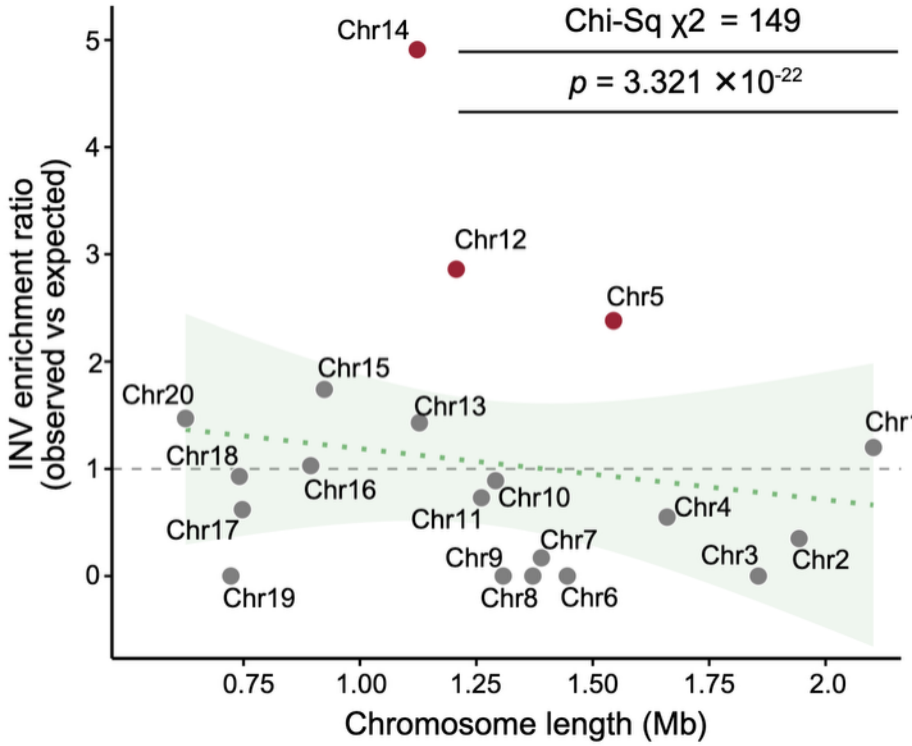


**Supplementary Fig. 11. Chromosomal enrichment of inversions relative to chromosome length.** Scatter plot showing the structural variant (SV) enrichment ratio (observed/expected number of inversions, INV) for each of the 20 chromosomes as a function of chromosome length (Mb), based on a total of 110 inversions detected genome-wide. The dashed horizontal line at ratio = 1 indicates the null expectation of proportional distribution by chromosome length; the green dotted line and shaded band show the linear regression trend and 95% confidence interval across chromosomes. Chromosomes significantly enriched for inversions (Chr5, Chr12, and Chr14; highlighted in red) show 2- to 5-fold more inversions than expected given their length. A chi-squared test indicates the overall distribution of inversions across chromosomes deviates significantly from the length-proportional null expectation ($\chi^2 = 149.06$, $P = 3.321 \times 10^{-22}$).

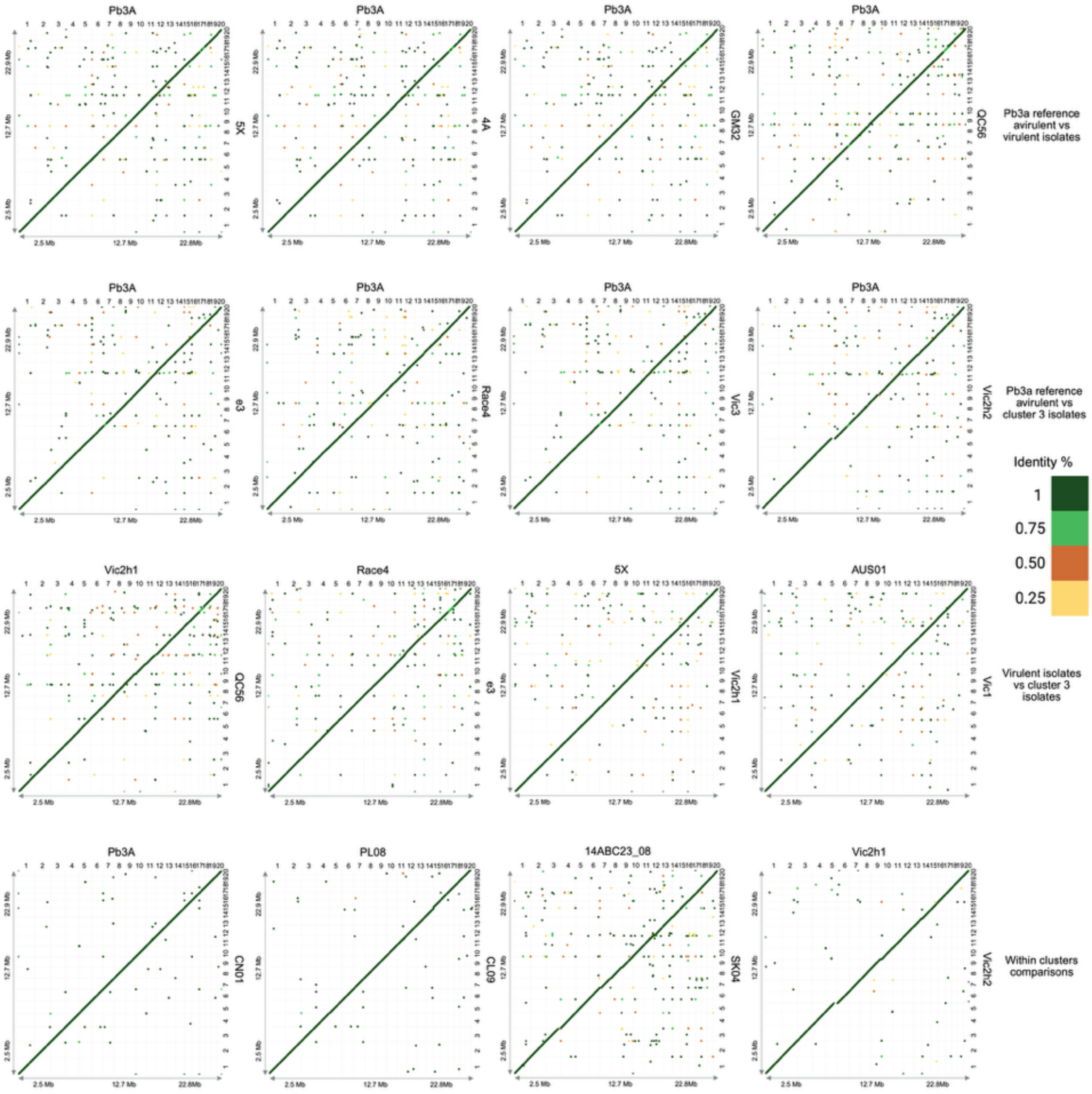


**Supplementary Fig. 12. Whole-genome alignment dot plots among *P. brassicae* isolates.** Pairwise whole-genome alignments are displayed as dot plots for representative assemblies, arranged in four comparisons. In each panel, the target assembly is plotted on the horizontal axis (top label), and the query assembly on the vertical axis (right label); both axes are scaled in genomic coordinates (2.5–22.8 Mb) with chromosomes numbered 1–20. Aligned segments are coloured by sequence identity (key, right): dark green ≈1.0, light green ≈0.75, orange ≈0.50, yellow ≈0.25. The continuous diagonal indicates collinear, one-to-one correspondence, whereas off-diagonal points denote inversions, translocations, and interruptions along the diagonal mark presence/absence variation. **Top row**, avirulent reference Pb3A versus virulent Cluster 1 isolates (5X, 4A, GM32, QC56). **Second row**, Pb3A versus Cluster 3 isolates (e3, Race4, Vic2h1, Vic2h2). **Third row**, virulent versus Cluster 3 isolates (Vic2h1 × QC56, Race4 × e3, 5X × Vic2h1, AUS01 × Vic1). **Bottom row**, within-cluster comparisons (Pb3A × CN01, PL08 × CL09, 14ABC23_08 × SKO4, Vic2h1 × Vic2h2). The recurrent chromosome 17 difference in the avirulent-versus-virulent tier (top row) marks a candidate structurally divergent region associated with the resistance-breaking Cluster 1 lineage.

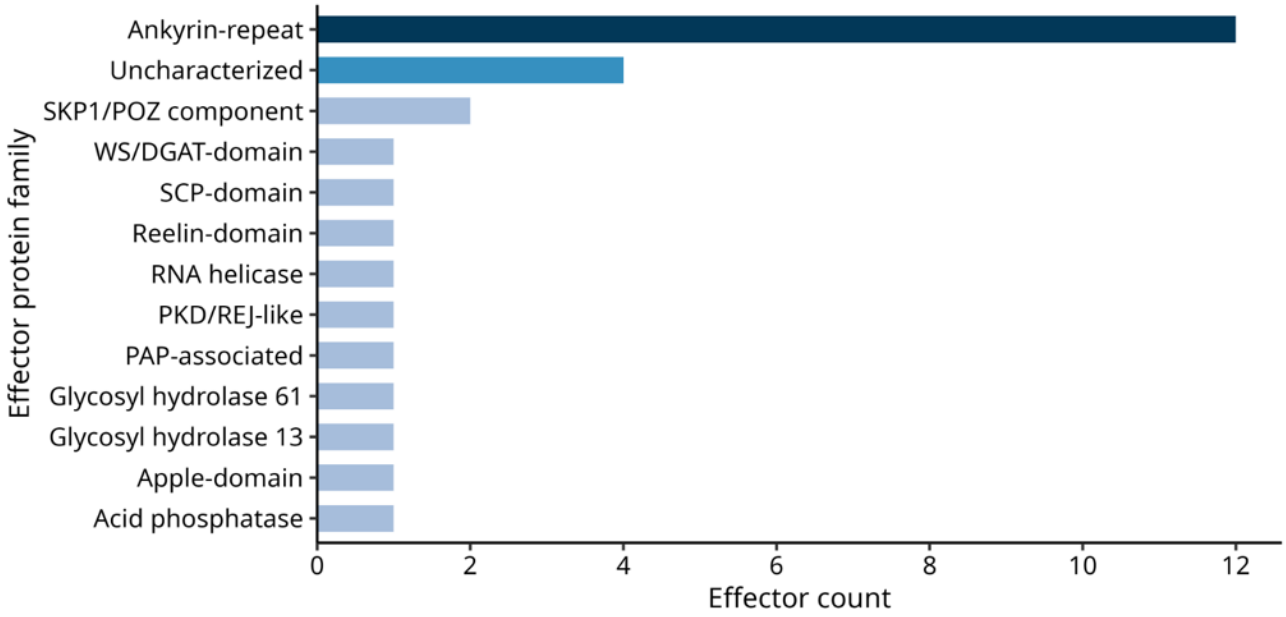


**Supplementary Fig. 13. Functional protein family composition of effector gene cluster of the Chr 17.** Horizontal bar chart showing the number of predicted effector proteins assigned to each protein family (n = 28 effectors across 13 families). Ankyrin-repeat proteins are the dominant family, accounting for 12 of 28 effectors (43%), followed by uncharacterized proteins (n = 4) and SKP1/POZ-domain proteins (n = 2); the remaining ten families are each represented by a single effector.

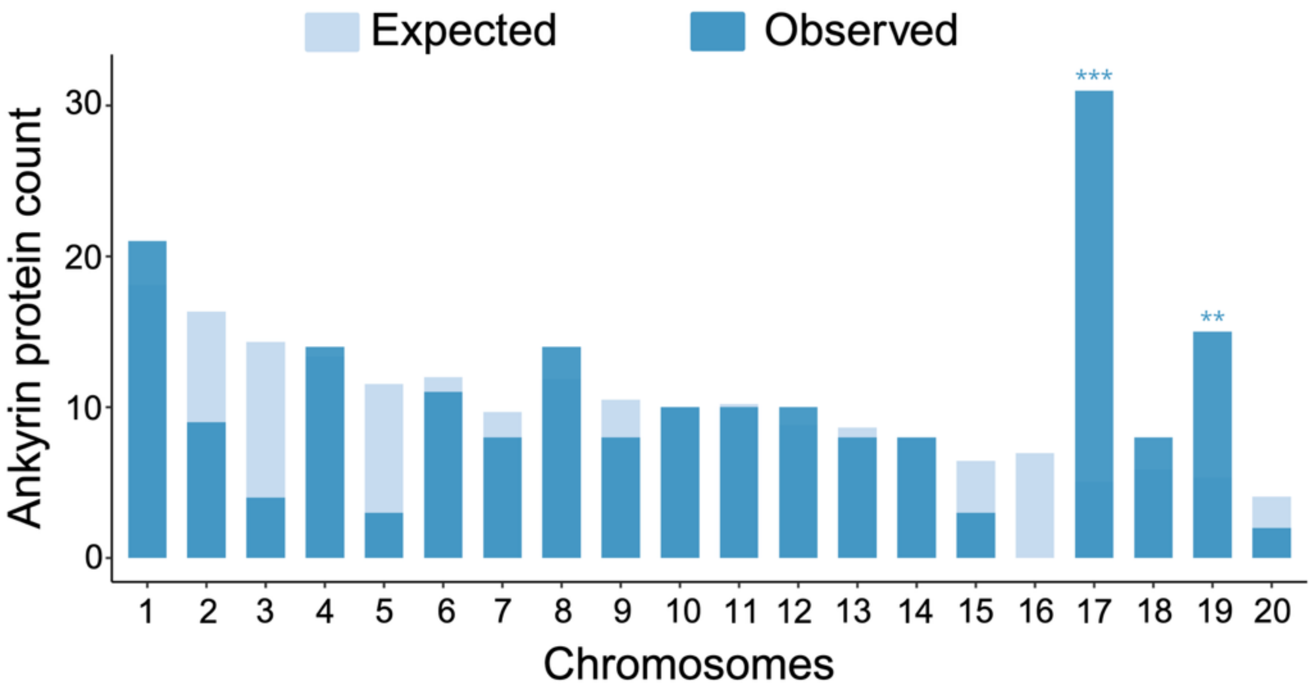


**Supplementary Fig. 14. Observed versus expected genomic distribution of ankyrin-repeat (ANK) genes across chromosomes.** Bar chart showing the observed number of ankyrin-repeat (ANK)-domain-containing genes (solid bars) compared to the number expected under a null model of uniform distribution proportional to chromosome length/gene content (light blue bars) for each of the 20 chromosomes. Asterisks indicate statistical significance of deviation from the expected distribution (Benjamini-Hochberg corrected; *$p < 0.05$, **$p < 0.01$, ***$p < 0.001$). Chromosome 17 shows a highly significant excess of ANK genes relative to expectation ($p < 0.001$), with Chromosome 19 also significantly enriched ($p < 0.01$), while Chromosome 3 shows a significant depletion relative to the null expectation ($p < 0.01$).

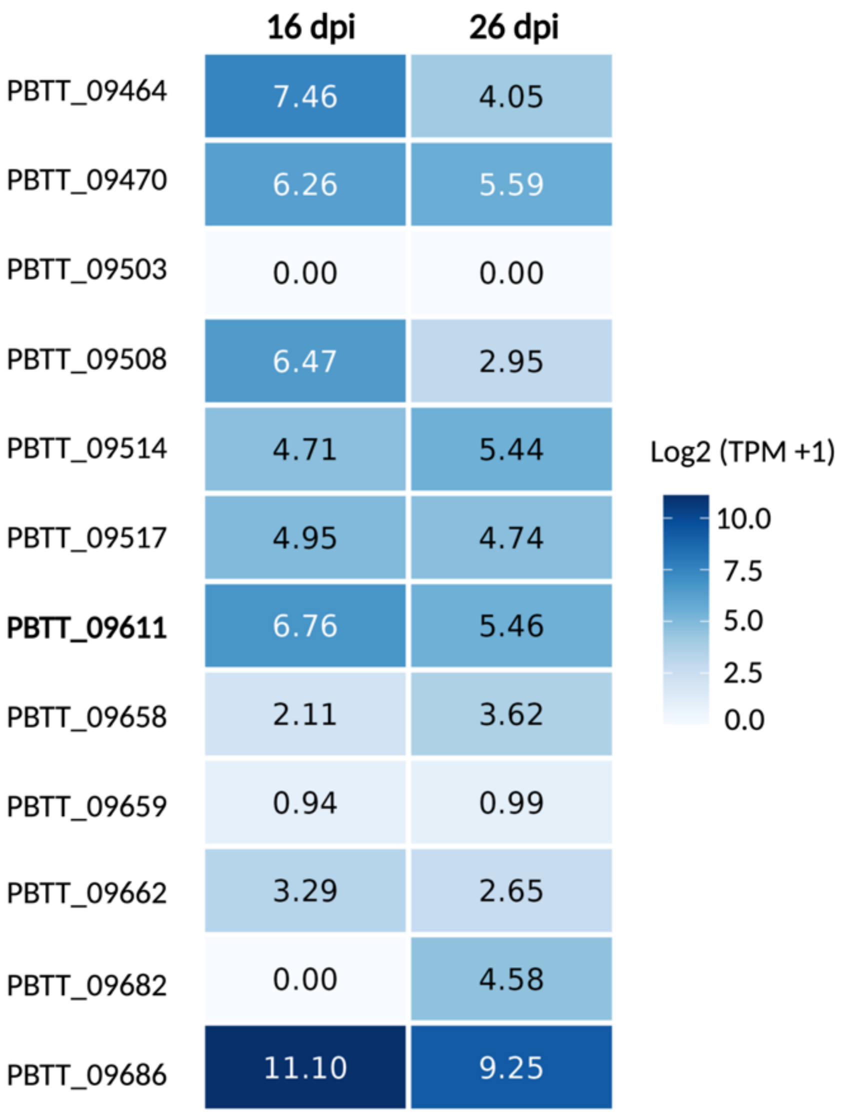


**Supplementary Fig. 15. Expression of effector candidate genes at specific infection time points.** Heatmap showing expression levels (log2[TPM+1]) of 12 effector candidate genes at 16 and 26 days post-inoculation (DPI), ranked by expression level at 16 DPI. Gene *PBTT_09611* (outlined in bold) is highlighted as the gene of interest. Most genes show decreased expression from 16 to 26 DPI, consistent with a shift in effector deployment over the time of infection.

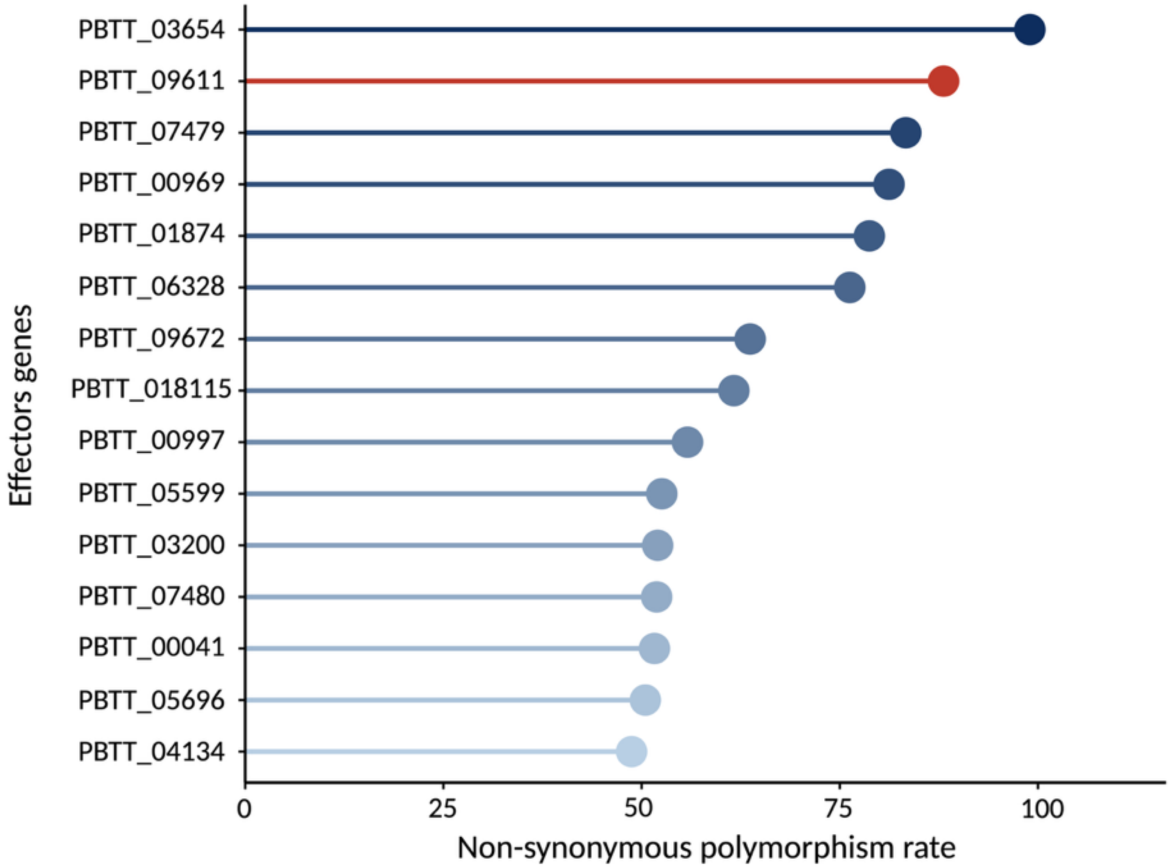


**Supplementary Fig. 16. Non-synonymous polymorphism rate (pN) across top effector candidate genes.** Lollipop plot of per-gene non-synonymous polymorphism rate (pN, expressed per 1,000 non-synonymous sites) for the 15 effector candidates with the highest pN in Cluster 3 (n = 28), ranked from highest to lowest. The functionally validated focal effector PbANK2 (PBTT_09611; red) shows the second-highest pN (88.1 per 1,000 non-synonymous sites). Colour gradient (dark navy to light blue) tracks descending pN.

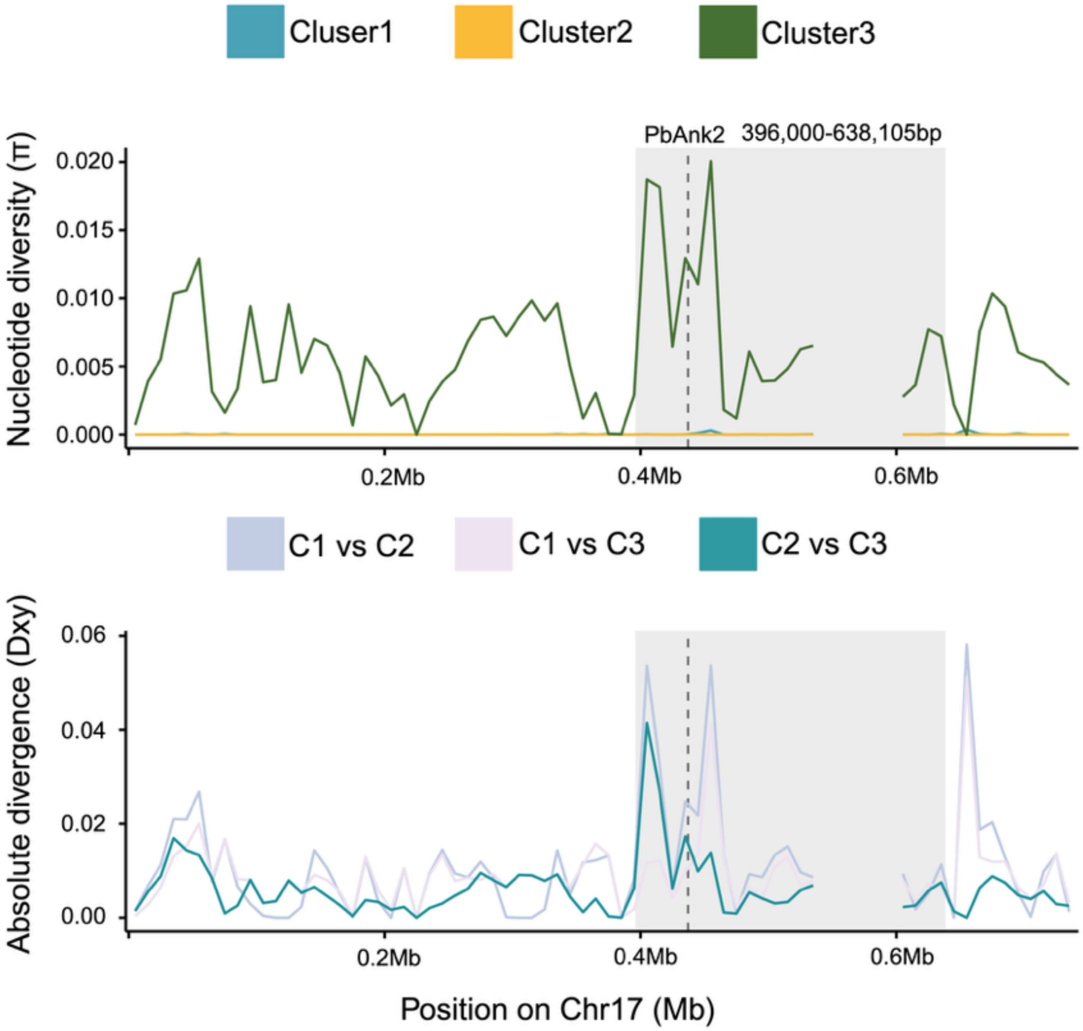


**Supplementary Fig. 17. Nucleotide diversity and absolute divergence across Chr17.** Population genetic statistics are plotted in sliding windows along Chr17 (x-axis, Mb) for the three *P. brassicae* clusters. **Top**, within-cluster nucleotide diversity (π) for three Clusters. Cluster 3 maintains substantial diversity along the chromosome (π up to ≈0.02), whereas Clusters 1 and 2 are near-monomorphic (π ≈ 0) throughout, consistent with clonal expansion of a single dominant genotype in these lineages. **Bottom**, absolute divergence (D*xy*) for the three pairwise cluster contrasts. The grey shaded band marks the *PbAnk2* region (Chr17:396,000–638,105 bp) and the dashed vertical line indicates the position of the *PbAnk2* gene. Both diversity and divergence rise sharply at the proximal boundary of this region (~0.4 Mb), Cluster 3 π peaks within the interval, and all three d*xy* comparisons show a coincident divergence peak, with the strongest signal in the contrasts involving Cluster 3. The gap near ~0.55–0.6 Mb reflects masked repeats gap. Together these profiles pinpoint the *PbAnk2* locus as a focal point of both intra-Cluster 3 diversity and between-cluster divergence on the Chr17 region that distinguishes virulent from avirulent lineages.

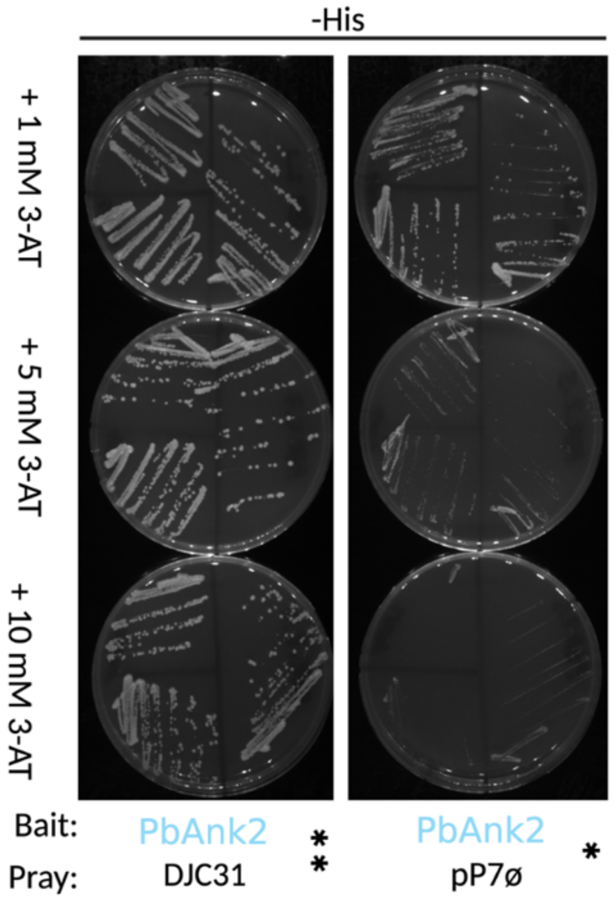


**Supplementary Fig. 18. Targeted yeast two-hybrid assay confirms PbAnk2–DJC31 interaction.** Yeast co-transformants were streaked on histidine-dropout medium (–His) with 3-AT at 1, 5 and 10 mM (rows) to suppress background His3 activation and increase assay stringency. **Left**, PbAnk2 bait with DJC31 prey; **right**, PbAnk2 bait with empty prey vector (pP7ø) as an autoactivation control (quadrants = independent streaks). PbAnk2 + DJC31 grew robustly at all 3-AT concentrations, whereas the empty-vector control was abolished at 5 and 10 mM. Sustained growth of the PbAnk2–DJC31 pairing under conditions that eliminate bait autoactivation confirms DJC31 as a direct interacting partner of the PbAnk2 effector.

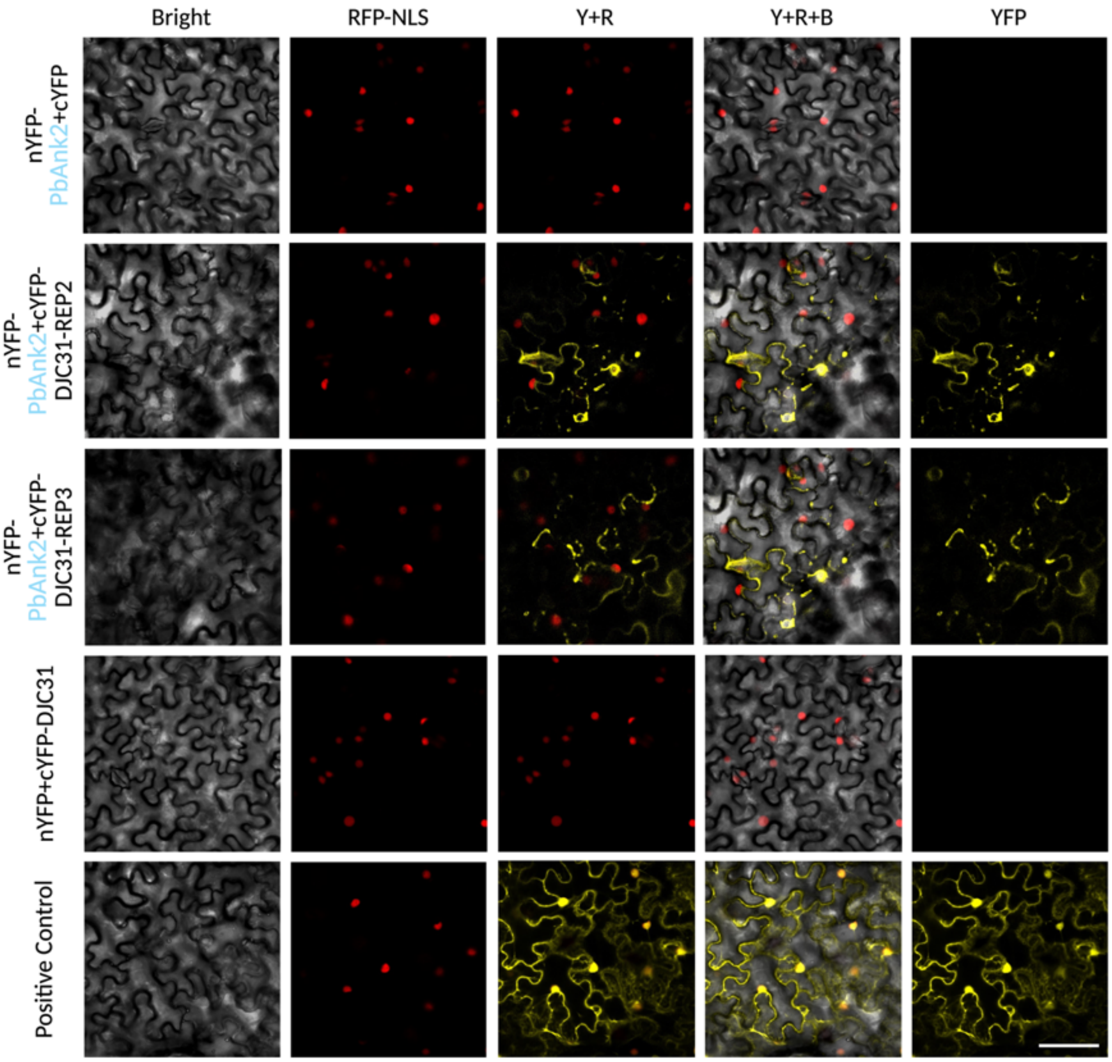


**Supplementary Fig. 19. Bimolecular fluorescence complementation (BiFC) confirms the PbAnk2–DJC31 (Avirulent allele) interaction *in planta*.** Co-expression of nYFP-PbAnk2 with cYFP-DJC31 reconstituted strong YFP fluorescence in the cell cortex and cytoplasm (rows 2–3), comparable to the positive control, whereas neither empty-vector control (rows 1, 4) produced YFP signal, demonstrating a specific PbAnk2–DJC31 interaction.

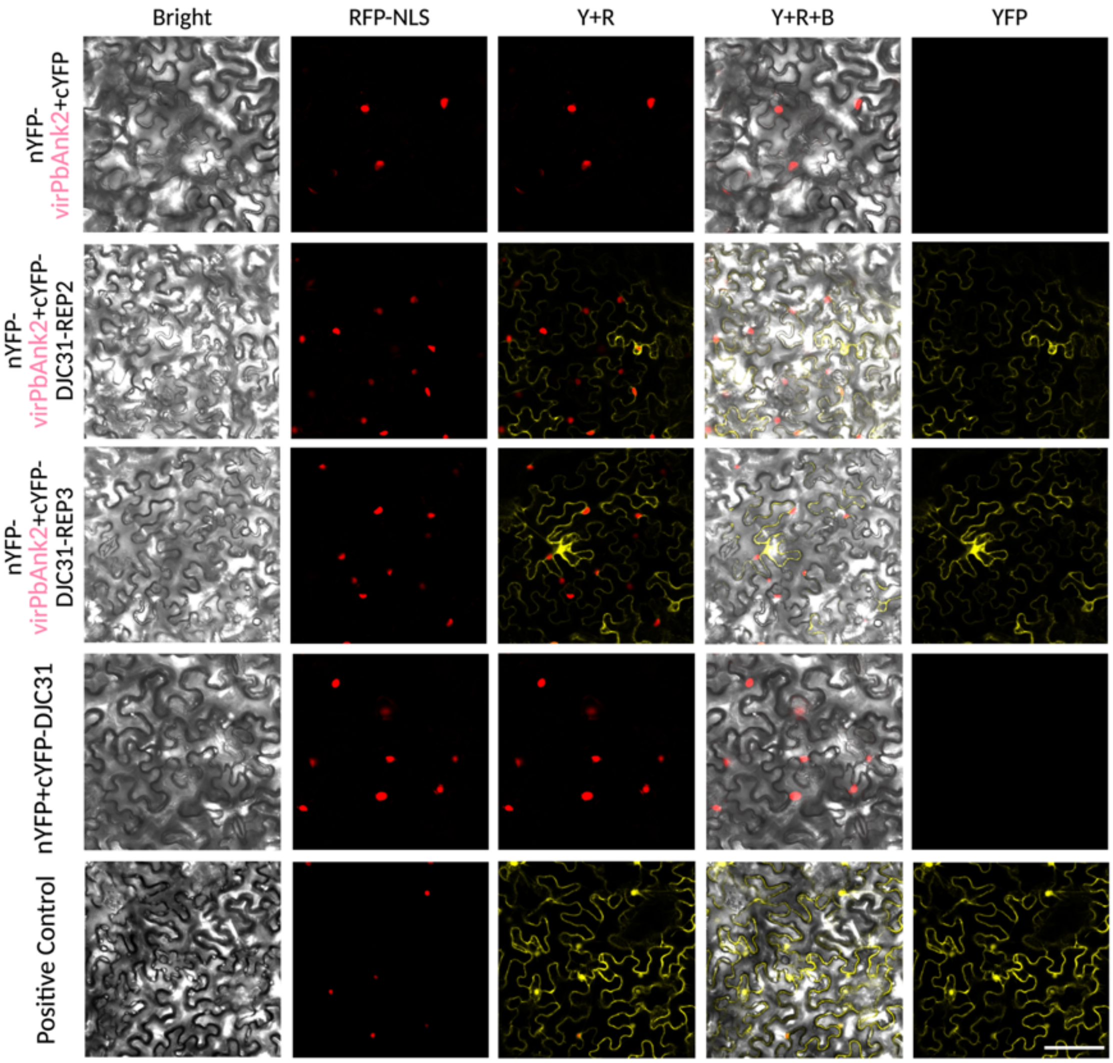


**Supplementary Fig. 20. Bimolecular fluorescence complementation (BiFC) confirms the virPbAnk2–DJC31 (Virulent allele) interaction *in planta*.** nYFP-virPbAnk2 co-expressed with cYFP-DJC31 yielded markedly weaker and more restricted YFP reconstitution (rows 2–3) relative to the avirulent allele, while all negative controls (rows 1, 4) remained YFP-negative and the positive control was unaffected. This indicates that the virulent PbAnk2 variant retains the DJC31 interaction.
